# Genetic effects on migration behaviour contribute to increasing spatial differentiation at trait-associated loci in Estonia

**DOI:** 10.1101/2023.10.25.564036

**Authors:** Ivan A. Kuznetsov, Estonian Biobank Research Team, Mait Metspalu, Uku Vainik, Luca Pagani, Francesco Montinaro, Vasili Pankratov

## Abstract

There is emerging evidence that migration behaviour can be selective with respect to individuals’ genotypes. As a result, it can generate genotype-environment correlations at migration-associated loci that cannot be corrected by standard methods used in genetic association studies. Here, we explore this phenomenon by looking into the dynamics of the spatial distribution of polygenic scores (PGSs) in the Estonian Biobank. We show that contemporary migrations intensify inter-regional differences in PGSs for many traits, with the effect of educational attainment (EA) PGS being the strongest and, to a large extent, explaining inter-regional differences in other PGSs. This differentiation is mainly driven by the migration of individuals with relatively high PGS for EA to the two largest cities from the rest of the country. Importantly, we replicate this pattern within families: individuals migrating to the major cities have, on average, higher PGS for EA relative to their siblings residing in other regions of Estonia. This trend is observed for the period starting from the mid-20th century to the present, despite substantial changes in Estonian society during this period. Thus, we show that there is increasing genetic differentiation at trait-associated loci between most urbanized regions of Estonia versus the rest of the country, leading to genotype-environment correlations that cannot be fully corrected using standard approaches. We also provide evidence for direct genetic effects on migration behaviour based on sibling analysis and discuss potential links between migration and EA.

## Introduction

Spatial genetic structure, i.e., differences in allele frequencies across geographic locations^1^, has been observed in human populations from global^2,3^ to fine scales^4–8^. It is driven by various demographic phenomena, including migrations and admixture as well as isolation due to geographic and cultural factors^9–13^. Spatial genetic structure may cause spurious genotype-phenotype associations and is routinely corrected for in genetic studies^14–17^. At a fine scale, such structure is being blurred by migration that has largely intensified in the past century^18–21^. If migration is equally likely to happen in any direction, it should mutually randomize environment and allele frequencies, reducing environmental confounding in genome-wide association studies (GWAS) and downstream analyses^17,22–24^. In practice, however, migration patterns may be associated with individuals’ genotype at certain loci. In the case of directional migrations (for instance, migrations to more economically developed areas) over generations, this will lead to increasing differentiation in allele frequencies at migration-associated loci between regions.

Indeed, Abdellaoui et al. have shown that migrants and non-migrants from the same economically deprived areas in Great Britain differ in their average genetic profiles, with the strongest difference observed for alleles associated with educational attainment (EA)^25^. As a result, this newly emerging genetic structure might generate genotype-environment correlations, which are the source of bias for the genetic effect estimates and complicate the interpretability of the results of genetic studies^26^. Recent works demonstrate that traits related to socioeconomic status (SES) are particularly prone to such complications^25–27^.

Despite the potential practical implications of such non-random changes in fine-scale spatial genetic structure due to recent human migrations, little is still known about how widespread and how recent they are. Most of the observations to date come from the UK Biobank^28^, raising the question of whether these effects are country- or cohort-specific. Additional analyses are also required to further characterize this phenomenon in terms of affected phenotypes, effects of confounding, and temporal dynamics.

Here, we aim to analyse the Estonian biobank dataset^29,30^ in order to assess the genetic consequences of recent migrations within Estonia, a country which is characterised by different genetic background, and demographic and socio-economic aspects when compared to Britain. Estonia has a centuries-old population structure that aligns with the broader European context but carries unique local patterns^7,10^. Moreover, Estonia has one of the largest internal migration rates in Europe, with approximately half of the population moving at least once in their lifetime^31^. The recruitment strategy of the EstBB also differs from that of the UK Biobank. The EstBB includes data on more than 210,000 participants, which represents approximately 20% of the current adult population of all ages and a relatively uniform geographic coverage^30^.

In this work, we explore how contemporary migrations (defined as the difference between an individual’s place of birth and place of current residence) change the spatial distribution of polygenic scores (PGSs) for 169 complex traits. We show that migrations lead to increasing differentiation between regions of Estonia (specifically the two major cities versus the rest of the country) in most of the tested PGSs, with the strongest effect for PGS for educational attainment. Using the within-family approach, we demonstrate that this association between genotype and migration profile reflects direct genetic effects and cannot be fully accounted for by parental effects or confounding. Next, we reveal that the inter-regional PGS differences accumulate over generations regardless of substantial changes in society. Finally, we discuss the potential implications of such migration-driven genetic structure for genetic studies.

## Results

### Data overview

We investigated the distribution of genetic principal components and polygenic scores for complex traits across geographic areas and between different migration groups in the Estonian Biobank (EstBB)^29,30^. We used genome-wide single-nucleotide polymorphism (SNP) data from 183,576 adults of European genetic ancestry who were born in Estonia, resided there at the time of joining the EstBB, and indicated ‘Estonian’ (172,376) or ‘Russian’ (11,200) when answering a question about ethnicity in the EstBB questionnaire. We treated these two cohorts separately, and we refer to them as Estonians and Russians, respectively. We stress that this division was based on self-identity and is made primarily to control for potential historical and cultural differences in migration patterns between these groups. The cohort of self-reported Estonians was used for all the main analyses. To control for group-specific effects and to provide a comparison across subgroups, we repeated some analyses in partially overlapping subsamples based on demography (sex and age) and time of the biobank enrolment, which occurred in two different recruitment campaigns. We also repeated the analyses on the Estonian subcohort after excluding relatives up to and including the second degree to confirm the observations using independent data points. Most of the analyses were replicated in the cohort of self-reported Russians, the second largest group in the EstBB. Detailed subdivision information and a description of the groups can be found in Supplementary Note 1.

### Effect of recent migrations on regional differences in genome-wide ancestry and polygenic scores

Among all the EstBB participants involved in this study, 41% (75,384 out of 183,576) have their current county-level place of residence (POR) different from their place of birth (POB). Hence, we investigated whether, and to what extent, contemporary migrations affect the present-day genetic structure in Estonia. In doing so, we compared the extent of genetic differentiation across the 15 Estonian counties when grouping individuals based on POB versus POR. Specifically, we calculated coordinates for the top 100 genomic principal components (PCs) for each individual and then compared *Var_county_* – the proportion of variance for each principal component explained by POB or POR. POB and POR explain a non-zero proportion of variance for 100 and 98 PCs, respectively (Figure 1A, Supplementary Table 4), consistent with previous reports of population structure in Estonia^10^. Importantly, for all the first 100 PCs, *Var_county_* for POB is significantly higher than for POR, suggesting that contemporary migrations attenuate genetic structure on a genome-wide scale.

**Figure 1.**
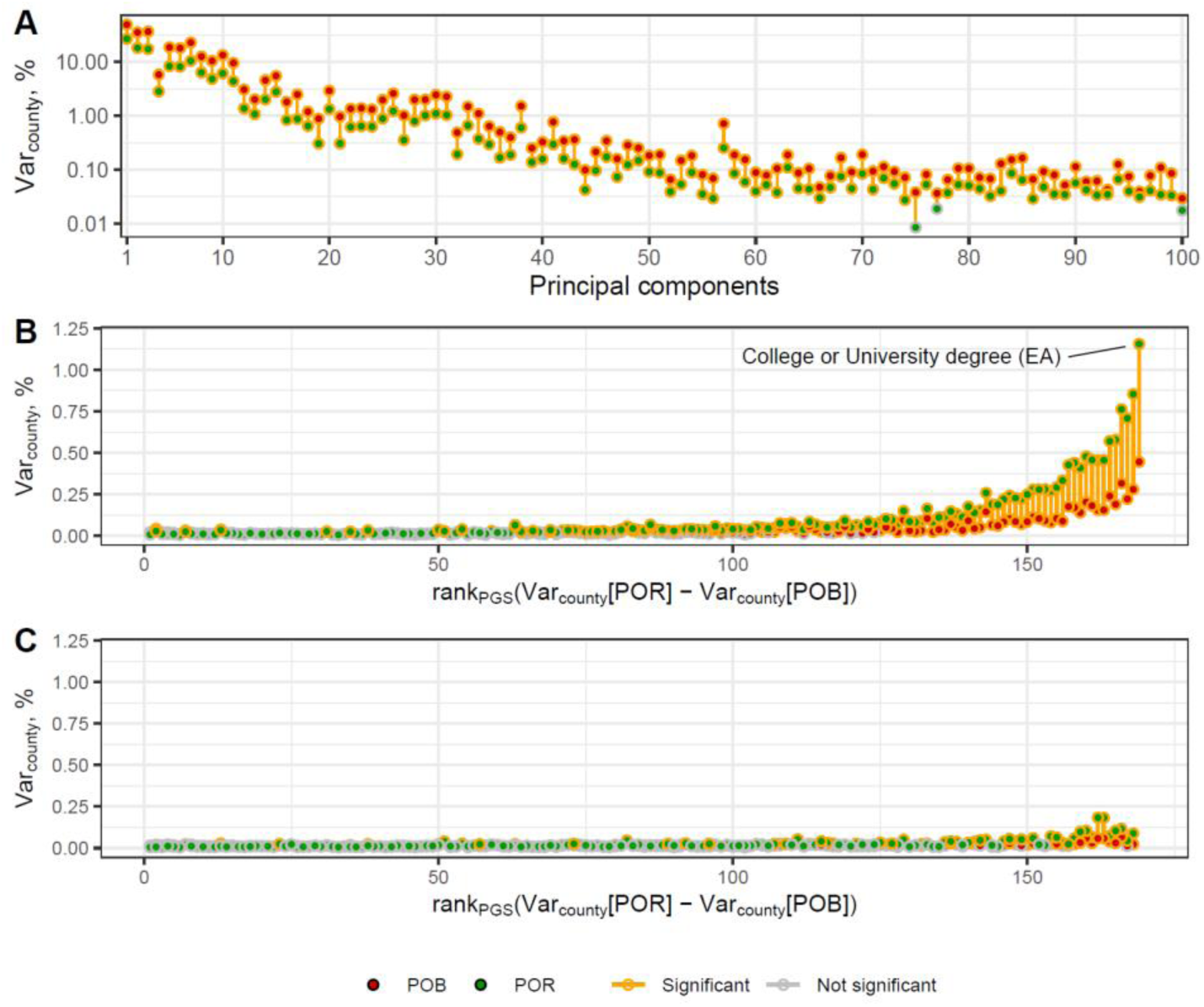
Fraction of the inter-individual variance of (A) PCs, (B) PGSs and (C) PGSs additionally adjusted for PGS_EA_, explained by county of birth (POB) and county of residence (POR). PGSs are preliminary adjusted for the top 100 PCs and demographic covariates. The y-axis in panel A has a logarithmic scale. The PGSs on the x-axis in panels B and C are ordered in the same way according to the difference between POR and POB *Var_county_*in panel B. Red and green dots refer to the POB and POR, correspondingly. Estimates significantly different from zero are outlined in yellow. The line connecting the two points is yellow when the variance explained by POB and POR together is significantly larger than the variance explained by only the weaker predictor (if significant) or when the stronger predictor is significant. The significance level is 0.05, after Bonferroni correction.

Certain genetic loci or sets thereof, however, might exhibit patterns different from the genome-wide ones. Indeed, contemporary migrations have been shown to be able to enhance regional differences in PGS for certain traits^25^. To verify these findings in the EstBB, we explored the spatial distribution of PGS for 169 diverse phenotypes, enriched with traits related to behaviour and SES (Supplementary Table 1). These PGS were calculated using summary statistics from GWAS in the UK Biobank subcohort of European ancestry^28,32^ and adjusted for demographic covariates and the first 100 PCs. Regional differences in POB and POR explain a statistically significant non-zero proportion of variance for 73 and 112 PGSs, respectively (Figure 1B, Supplementary Table 5). Unlike for the PCs, for 107 PGSs, the *Var_county_* values for POR are significantly higher than for POB. The effect was reversed for only 3 PGSs. Therefore, most PGSs show a geographic structure remaining after regressing out the first 100 PCs, and this structure is enhanced by contemporary migrations.

As the tested PGSs are intercorrelated (Supplementary Table 6), these results might reflect a single underlying phenomenon. Assuming a set of loci with a concordant pattern of allele frequency change across Estonian counties due to migrations, we expect multiple PGSs to have a non-zero *Var_county_*, with this statistic being higher for PGSs that better capture alleles with stronger frequency differentiation. To see whether different PGSs capture the same genetic structure pattern and change thereof we repeated the analysis after correcting the tested PGSs for the PGS for ‘College or university degree’ (PGS_EA_), which is the PGS with the highest *Var_county_* for both POB and POR (0.45% and 1.16%, respectively). Among PGSs with *Var_county_*significantly greater than zero before the adjustment, the adjustment for PGS_EA_ leads to a decrease of *Var_county_* in all 72 cases for POB and for 110 out of 111 cases for POR (Figure 1C, Supplementary Table 7). *Var_county_*remains significantly higher than zero for 32 and 48 adjusted PGSs, not exceeding 0.07% and 0.19%, for POB and POR, respectively. This result suggests that the loci associated with university education, or more broadly with EA, contribute to a substantial fraction of the signal of non-random distribution of the other PGSs in space.

If the signal described above is indeed mostly linked to the EA-associated loci, we expect that correcting for a more powerful PGS for EA will reduce the signal for other PGSs even further. We indeed confirmed this by using summary statistics from a meta-analysis GWAS (PGS_EA4_)^33^. *Var_county_* for POB (0.50%) and POR (1.53%), and the difference between them are higher for PGS_EA4_ than for PGS_EA_. Adjustment for PGS_EA4_ reduces the *Var_county_*for other PGSs to a greater extent than adjustment for PGS_EA_ (21 and 37 remain significant with a maximum of 0.05% and 0.14% for POB and POR, respectively) (Supplementary Figure 23, Supplementary Table 8). These results support the hypothesis that most of the inter-regional differences in the PGSs and the increase of these differences due to contemporary migrations can be considered as a single phenomenon best captured by variants associated with EA, regardless of the mechanism of these associations. PGS_EA4_ is more powerful but potentially more confounded than PGS_EA_ as it is based on a meta-analysis of many relatively small cohorts in which the adjustment for population structure can be less effective^34,35^. Thus, throughout the rest of our analyses, we mainly focused on PGS_EA_, which captures most of the signal of non-random distribution of the tested PGSs, yet carries potentially less confounding than PGS_EA4_.

Next, we explored whether the increase of *Var_county_*for PGS_EA_ can be an artifact due to some unaccounted properties of the EstBB sample. We showed that the increase in *Var_county_* for PGS_EA_ remains significant if we a) stratify the sample by sex, age, or recruitment phase; b) filter out relatives up to second degree included; c) repeat the analysis in the cohort of self-reported Russians; d) adjust PGS_EA_ for the complete genetic relatedness matrix (GRM) in a leave-one-chromosome-out (LOCO) approach. We also showed that adjusting PGS_EA_ for PCs has a minor effect on the difference in *Var_county_* between POR and POB, with a weak dependence on the number of PCs used. Additionally, e) the polygenic score derived from the summary statistics of within-sibship GWAS for EA^36^ also demonstrates a significant increase in *Var_county_*. See Supplementary Notes 4 and 5 (Supplementary Figure 17) for details.

Finally, f) the relatively large number of sibships in the EstBB allowed us to repeat the analysis for PC or PGS deviations from their within-sibship mean values. Such sibling design randomizes genotypes and environment, allowing differentiation of genetic effects from associations due to environmental confounding^37,38^. As we used only sibships, in which all the members were born in the same county, and thus *Var_county_*for POB is zero by design, we compared *Var_county_* for POR to zero and calculated empirical p-values. For the PC coordinates, *Var_county_*for POR is in the range between 7.3×10^−5^ and 5.5×10^−4^ with one out of 100 PCs reaching significance (PC76: Var_county_ = 5.5×10^−4^, p-value_Bonf_ = 0.02; Figure 2A; Supplementary Table 9). In addition to the 169 UKB-based PGSs, we include PGS_EA4_ in the analysis as the sibling design is robust to environmental confounding inherited by a PGS from the corresponding GWAS^39^. *Var_county_*for POR is significantly different from zero for 12 PGSs after correction for multiple testing (Figure 2B). These PGSs correspond to the phenotypes related to SES or cognitive skills, with EA4 having the highest *Var_county_* (*Var_county_* = 1.4×10^−3^, p-value_Bonf_ = 0.017). When we repeated the analysis using PGSs adjusted for PGS_EA4_, *Var_county_* for POR did not exceed 5×10^−4^ and never reached the significance level (Figure 2C). The sibling design is underpowered compared to our main analysis due to reduced sample size and because of ignoring cases of identical migration of individuals from the same sibship. Nevertheless, these results confirm that migration behaviour is associated with a specific genetic component that, out of the PGSs we tested, is best captured by PGSs for EA-related phenotypes. These associations lead to a non-random spatial distribution of the corresponding alleles and are at least partly driven by direct genetic effects.

**Figure 2.**
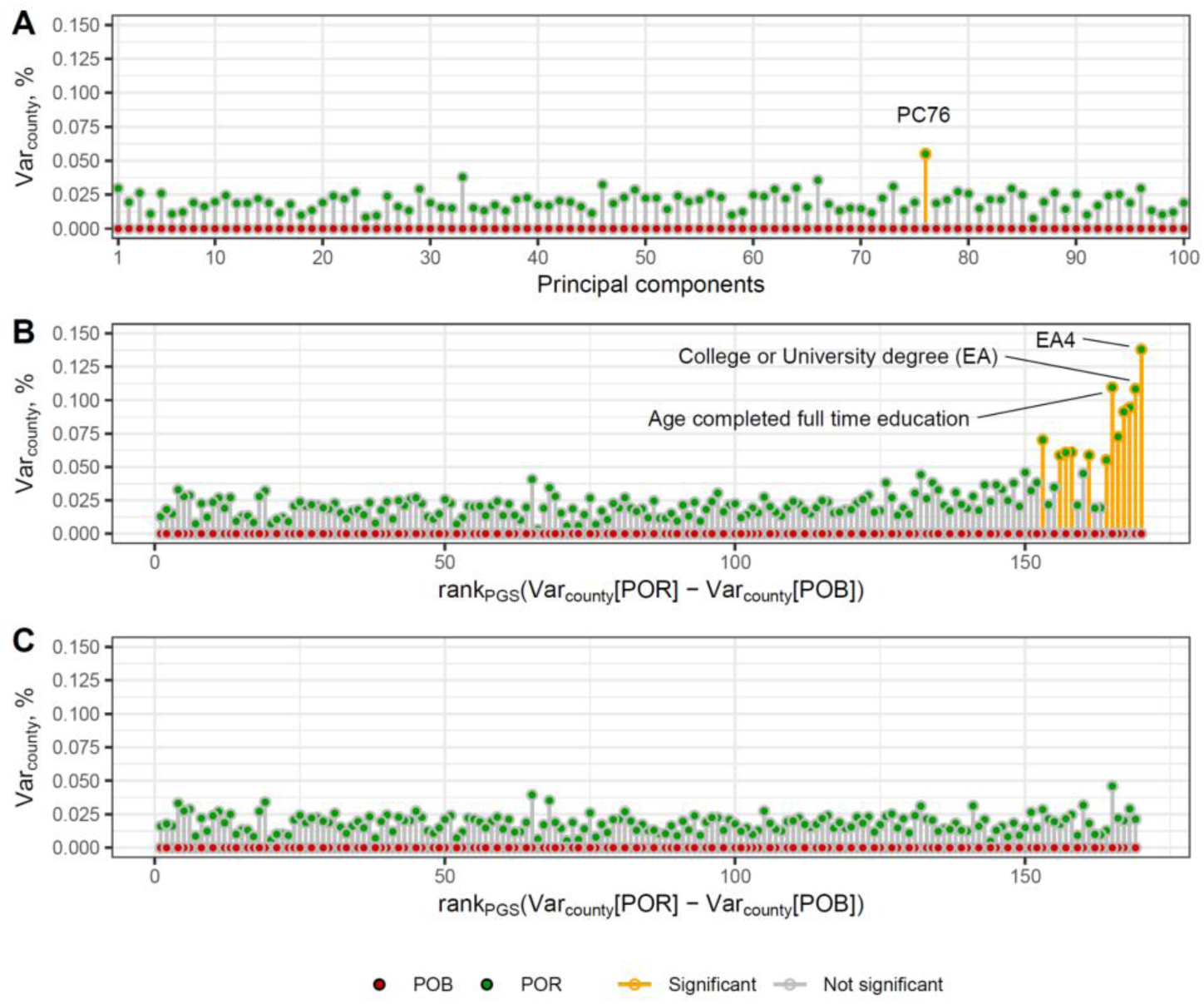
Fraction of the inter-individual variance of the deviation of individual’s value from the sibship’s mean for (A) PCs, (B) PGSs, and (C) PGSs additionally adjusted for PGS_EA4_, explained by county of birth (POB) and county of residence (POR). PGSs are preliminary adjusted for the top 100 PCs and demographic covariates. The PGSs on the x-axis in panels B and C are ordered as in Figure 1B but with PGS_EA4_ added as the right-most datapoint. Red and green dots refer to the POB and POR, correspondingly. Estimates significantly different from zero are outlined in yellow. *Var_county_* for POB is always zero by design of the analysis. The significance level is 0.05, after Bonferroni correction.

### Geographical distribution of PGS_EA_

To explore whether the increasing between-county variability of PGS_EA_ reported above is driven by specific Estonian regions, we mapped the mean values of PGS_EA_, adjusted for PCs and demographic covariates, for every county in Estonia (Figure 3A and B). For both POB and POR, two counties have values significantly higher than the country average: Harju (FDR-adjusted p-value 4.1×10^−77^ and 1.1×10^−168^, correspondingly) and Tartu (FDR-adjusted p-value 4.8×10^−12^ and 2.8×10^−14^, correspondingly). These counties are where Tallinn and Tartu, the two most populated Estonian cities, are located, together making up more than 40% of the country’s population^40^. Most other counties have values significantly lower than the country’s average.

**Figure 3.**
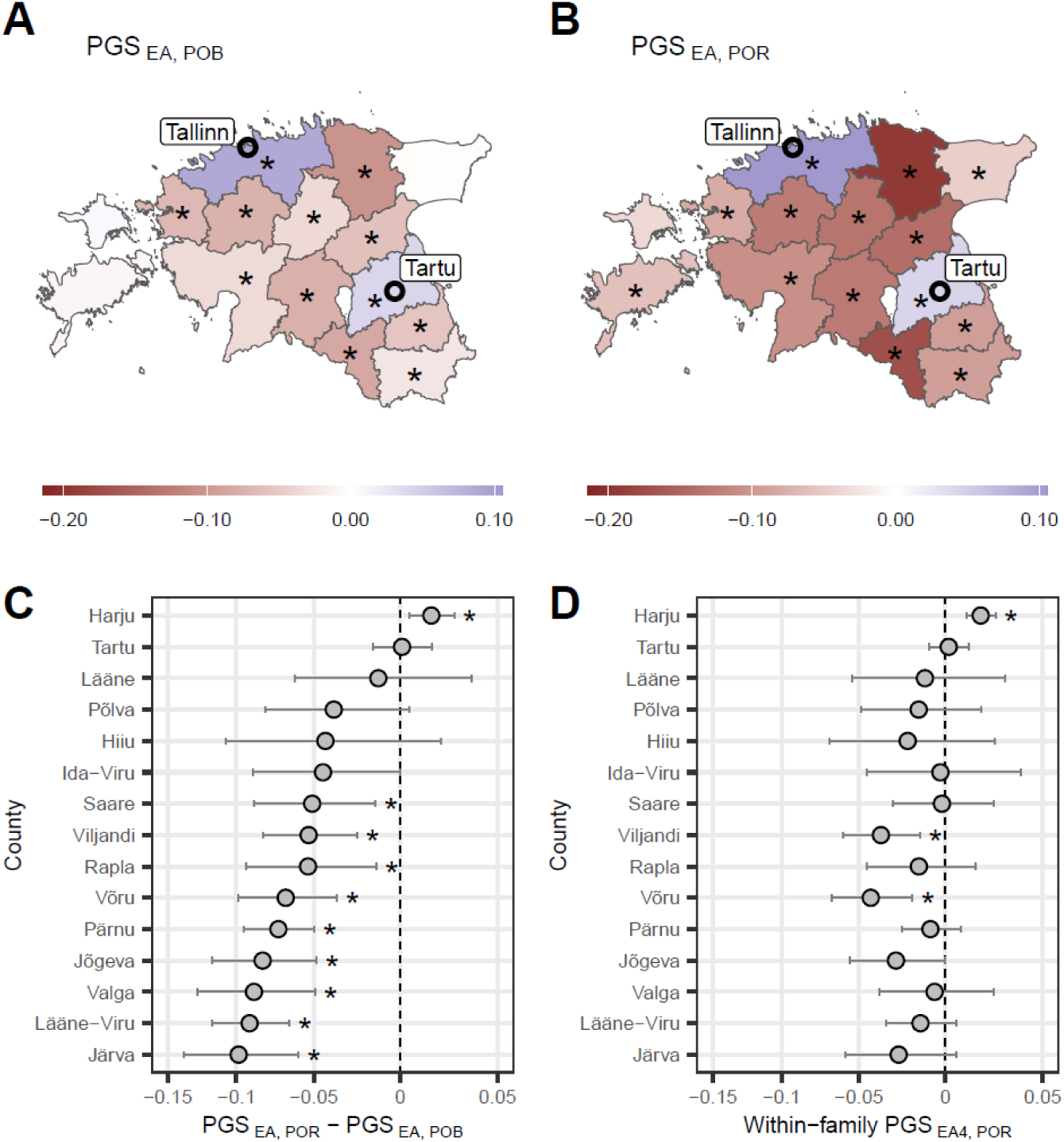
PGS_EA_ (PGS_EA4_) landscape in Estonia. Mean PGS_EA_ of individuals (A) born or (B) residing in each county. (C) Difference between values in panels B and A per county. In panels A-C PGS_EA_ is adjusted for the top 100 PCs and demographic covariates. (D) Mean value of PGS_EA4_ adjusted for sibship mean per county of residence. Only siblings born in the same county are included making the corresponding statistic for county of birth being zero for all counties. Counties with the corresponding value being significantly different from zero after FDR correction at the 0.05 significance level are marked with an asterisk (*). PGSs are measured in standard deviations.

To see how the mean PGS_EA_ changed due to contemporary migrations, we subtracted the mean values of PGS_EA_ individuals born in a corresponding county from the mean values of PGS_EA_ of the county’s residents (Figure 3C). This change is significantly positive for Harju County, which includes the capital Tallinn, significantly negative for nine counties, and is not significant for the remaining five. Notably, for Tartu County, the estimate has a narrow CI_95%,_ suggesting that recent in- and out-migrations counterbalance each other. These patterns are generally consistent across cohorts of Russians and unrelated Estonians, as well as in subcohorts stratified by sex, year of birth, and year of biobank enrollment (Supplementary Figures 19, 24, 37-40). A significant increase in average PGS_EA_ in Harju County is also observed on the within-sibship level, while the point estimates of change in the average PGS_EA_ in all other regions but Tartu County are negative (Figure 3D).

### PGS_EA_ values in groups with different migration profiles

Next, we compared the mean PGS_EA_ between groups with different migration profiles (Figure 4). For this, we divided Estonia into three areas: Harju County (including Tallinn), Tartu County (including Tartu City), and other regions of Estonia (referred to as ‘ORE’ below). All the individuals were classified into 9 groups based on their place of birth and residence. This classification was motivated by the results above and by the fact that Harju and Tartu Counties are the major destinations of migration in Estonia^41,42^, see also Supplementary Note 2. In all cases, migration within the defined areas (for instance, between counties defined as the ORE) was ignored.

**Figure 4.**
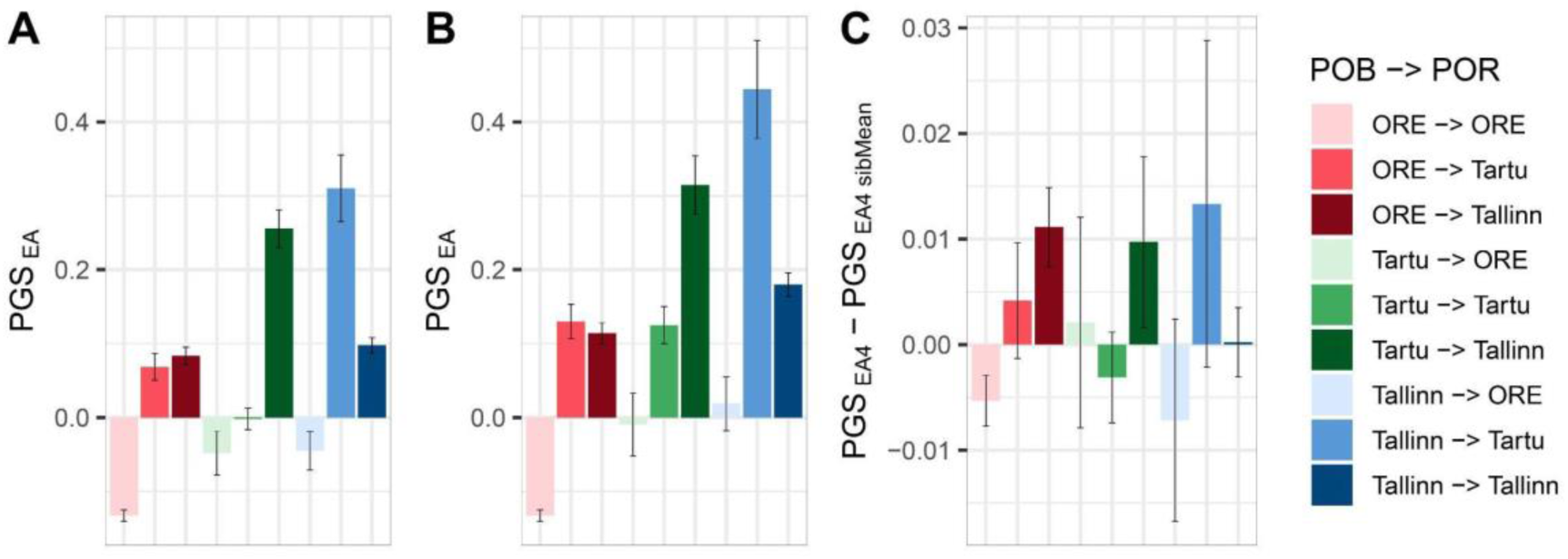
PGS_EA_ (PGS_EA4_) in migration groups defined by combination of place of birth (POB) and residence (POR). (A) County-based analysis where ‘Tartu’ and ‘Tallinn’ refer to Tartu County and Harju County respectively while ‘ORE’ refers to other counties. (B) City-based analysis, where ‘Tartu’ and ‘Tallinn’ refer to the respective cities while ‘ORE’ refers to other counties as in A. In A and B PGS_EA_ is adjusted for the top 100 PCs and demographic covariates. (C) County-based analysis for PGS_EA4_ adjusted for sibship-average. In all panels error bars correspond to 95% confidence intervals.

Individuals who moved to Harju or Tartu Counties from ORE have higher PGS_EA_ in comparison with those who stayed in ORE, explaining the decrease of PGS_EA_ in most ORE counties (Figure 4A). Among individuals born in Harju or Tartu Counties, those migrating to ORE have the lowest PGS_EA,_ while those moving between Tartu and Harju Counties have the highest PGS_EA_. Tallinn and Tartu are the two biggest cities in Estonia, the main hotspots of urbanisation, centres of education, and economic development. Therefore, we questioned if our results are also driven by those cities. To check this, we repeated the analysis, keeping only participants born/residing in Tallinn or Tartu City instead of the entire corresponding counties (Figure 4B). The results demonstrate an even larger contrast between those who were born in or moved to Tallinn or Tartu City and those who stayed in ORE. These patterns replicate in unrelated Estonians (Supplementary Figure 20), in subcohorts divided by sex, year of biobank enrollment, and some of the groups divided by year of birth (Supplementary Figures 49-51). They are also consistent with observations coming from the remaining subcohorts (Supplementary Figures 48, 50). That supports the hypothesis on the important role of the cities in the increasing contrast between the counties. PGS_EA_ adjusted for the sibship average also demonstrates significant differences between those who stayed in ORE and those who moved to Harju (p-value = 2.2×10^−13^) or Tartu County (p-value = 9.3×10^−4^) (Figure 4C) or to Tallinn (p-value = 9.6×10^−11^) or Tartu City (p-value = 3.4×10^−3^) (Supplementary Figure 26). It is also higher among migrants from Tartu County to Harju County than among those who stayed in Tartu County (p-value = 3.1×10^−3^).

We next investigated whether the PGS_EA_ of migrants to Tallinn and Tartu City varies based on an individual’s POB in a destination-specific manner. We calculated differences in mean PGS_EA_ between residents of Tallinn and Tartu City born outside those two cities, grouped by their county of birth (Figure 5). Individuals who migrated to Tallinn from counties surrounding Tartu City show, on average, higher PGS_EA_ compared to individuals born in the same counties and migrated to Tartu City. The opposite is true for the counties surrounding Tallinn. Subcohort analyses produce results generally in line with these observations, though they suffer from low power (Supplementary Figures 21, 53-56).

**Figure 5.**
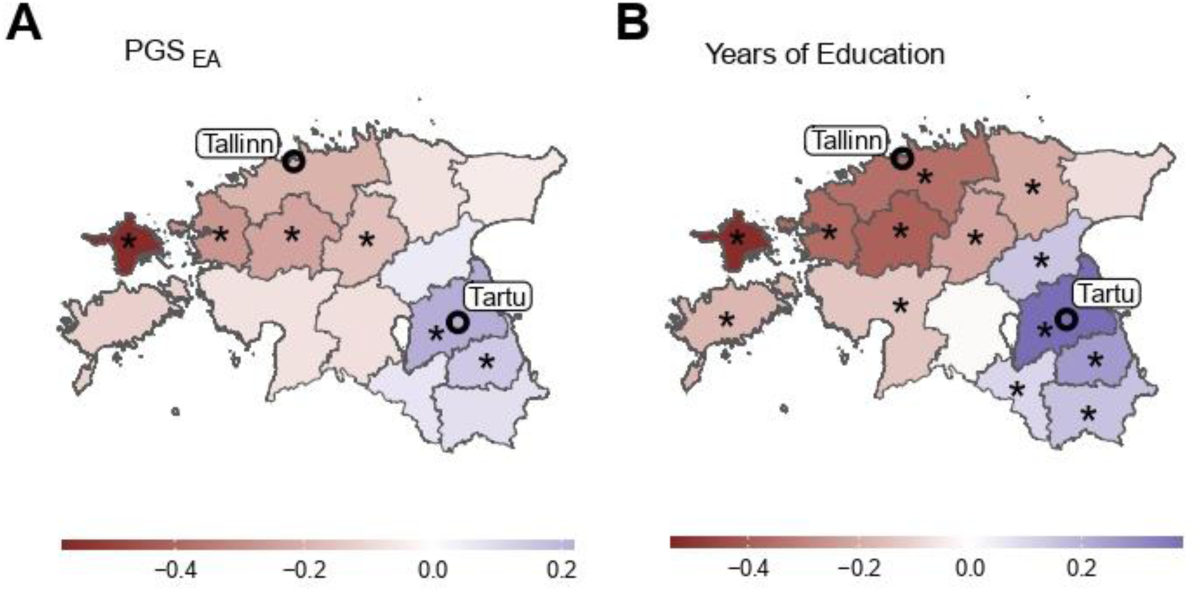
The difference in mean PGS_EA_ and EA (years of education) between residents of Tallinn and Tartu City by county of birth. (A) The value for each county corresponds to the mean PGS_EA_ of individuals born in that county and residing in Tartu City subtracted from the mean PGS_EA_ of individuals born in the same county and residing in Tallinn. Individuals born in Tallinn or Tartu City are excluded from the analysis. PGS_EA_ is adjusted for the top 100 PCs and demographic covariates. (B) The same but for the ‘years of education’ phenotype. Counties with the difference being significant after FDR correction at level 0.05 are marked with an asterisk (*).

### Genetic predictors of ORE-to-cities migration

The results above suggest that the pattern observed in Figures 1B and C is mostly driven by selective migration out of ORE to Tallinn and Tartu City. Thus, we explored the genetic differences between those moving out of ORE to the two major cities (‘cases’) versus those born and staying in ORE (‘controls’) in more detail. First, a SNP-based heritability estimate of 0.13 (CI_95%_: 0.10 - 0.16) (Table 1) confirmed that there are systematic genetic differences between migrants and non-migrants. Next, we tested the 169 UKB-based PGSs and PGS_EA4_ as predictors for out-of-ORE migration to Tallinn or Tartu City in unrelated individuals (29,306 cases and 14,028 controls) and in siblings (5,931 cases and 15,281 controls; 11,078 sibships)^43^. The latter approach allowed us to estimate between- and within-family effects separately. Within-family estimates of PGS effects are not confounded by genotype-environment correlations and parental indirect effects. We note, however, that this approach doesn’t control for indirect genetic effects of siblings, which are expected to correlate positively with direct effects^44^, thus introducing downward bias to the within-sibship effect estimates. In this analysis, we applied two models: a) mixed effects logistic regression, with sibship modeled as a random effect, and b) fixed effects logistic regression. The mixed effects model explicitly accounts for unexplained inter-sibship variability and is hence more appropriate, while the fixed effects model enabled us to compare estimates from the sibling subsample to the population-level effects derived from the subcohort of unrelated Estonians. The effects of PGS_EA4_ on the migration phenotype at the within-sibship, between-sibship, and population levels are significantly higher than zero and are the strongest among the tested PGSs, followed by PGS_EA_ (Figure 6A, Supplementary Figures 31, 32, 33A). All the PGSs with significantly non-zero within-sibship effects are related to SES. Estimates of the population effects for the PGSs from the unrelated Estonians and from siblings are consistent with each other, demonstrating an absence of strong biases in the sibling subsample in comparison with the whole sample (Supplementary Figures 32, 33). PCs do not show significant within-sibship effects on ORE-to-cities migration (Supplementary Figures 34, 35). Next, we regressed PGS_EA4_ out of other PGSs and found no significant within-sibship effects for the adjusted PGSs (Figure 6B, Supplementary Figure 33B). This result supports the idea that the loci associated with EA substantially contribute to the set of loci that directly influence an individual’s probability of ORE-to-city migration.

**Figure 6.**
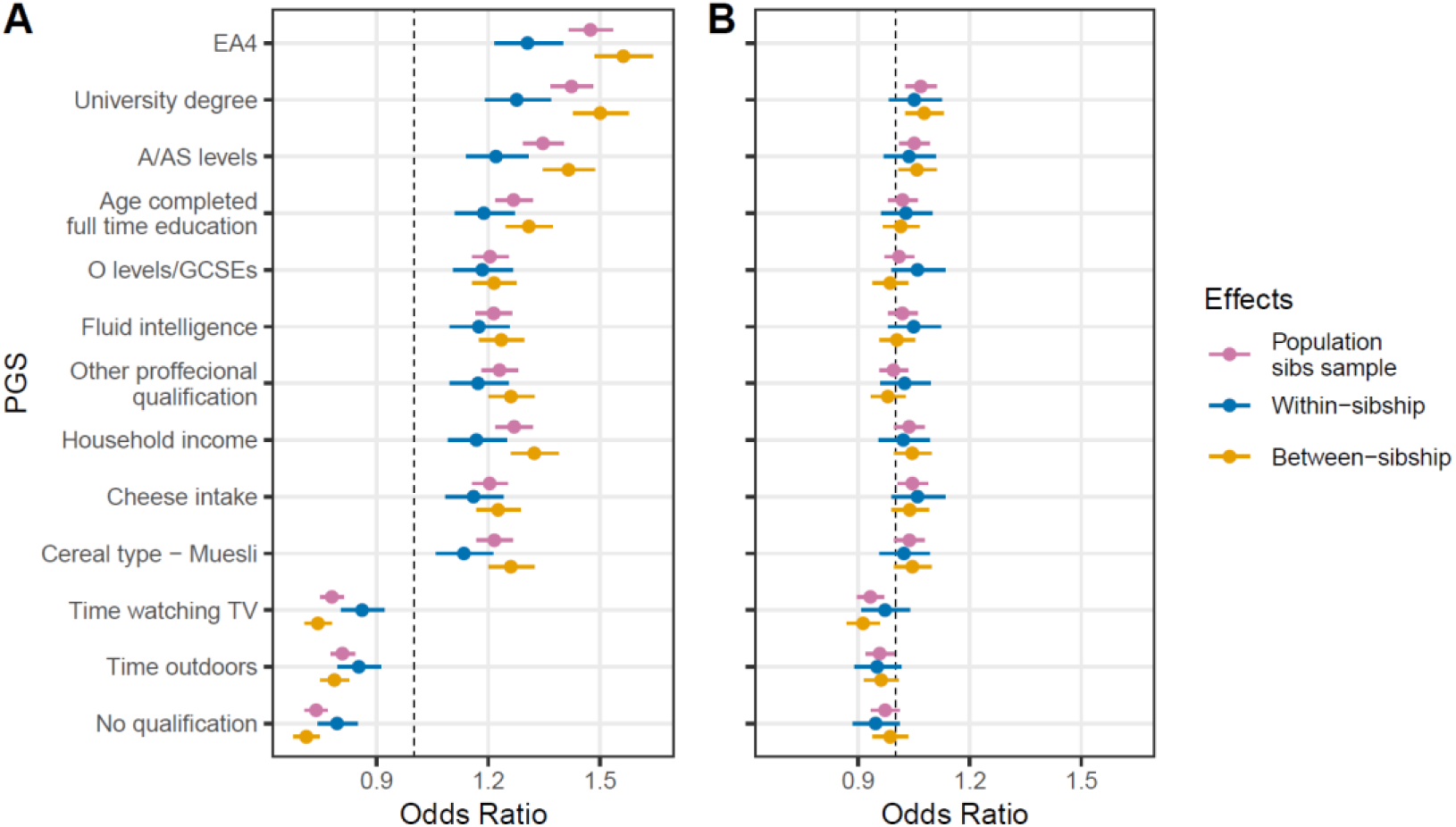
PGSs as predictors of migration from ORE to the major cities (Tallinn or Tartu). All the effects are estimated in a sample of siblings with only siblings born in the same county being included. The estimates are obtained using mixed effects logistic regression with a random intercept for sibship. (A) Effect sizes for PGSs; (B) Effect sizes for PGSs additionally adjusted for PGS_EA4_. All PGSs are preliminary adjusted for top 100 PCs and the demographic covariates. Results are shown for PGSs with a significant within-sibship effect before adjusting for PGS_EA4_ and after Bonferroni correction. Vertical dashed line indicates Odds Ratio equal to 1. Error bars correspond to 95% confidence intervals.

**Table 1.** Genetic aspects of the migration phenotype. The migration phenotype is defined for individuals born in ORE and residing in either Tallinn or Tartu City (cases) or in ORE (controls). The logistic regression section provides the odds ratio for PGS_EA_ as a migration predictor in a model without or with EA. Two models with EA as a covariate were tested: years of education translated from the reported categories of EA (Supplementary Table 3) and the reported categorical EA. Fixed effects model was used for within-sibship logistic regression for comparability with population-level effects. GREML-GCTA section tabulates heritability estimates for binary educational attainment - university degree (h^2^_EA_) and migration (h^2^_Migr_) as well as the genetic correlation between them in the corresponding cohort.

| **Population-level logistic regression, $\text{OR}_{\text{PGS(EA)}}$** |  |  |
| --- | --- | --- |
| | Estimate, $\text{CI}_{95}$ | P-value |
| $\text{PGS}_{\text{EA}}$ | 1.31 [1.29; 1.33] | $4.9 \times 10^{-258}$ |
| $\text{PGS}_{\text{EA}} + \text{Years of education}$ | 1.15 [1.13; 1.17] | $5.4 \times 10^{-64}$ |
| $\text{PGS}_{\text{EA}} + \text{EA (categories)}$ | 1.14 [1.12; 1.16] | $2.0 \times 10^{-57}$ |
| **Within-sibship logistic regression, $\text{OR}_{\text{PGS(EA)}}$** |  |  |
| | Estimate, $\text{CI}_{95}$ | P-value |
| $\text{PGS}_{\text{EA}}$ | 1.22 [1.15; 1.29] | $2.1 \times 10^{-10}$ |
| $\text{PGS}_{\text{EA}} + \text{Years of education}$ | 1.11 [1.04; 1.18] | $1.5 \times 10^{-3}$ |
| $\text{PGS}_{\text{EA}} + \text{EA (categories)}$ | 1.10 [1.03; 1.17] | $4.1 \times 10^{-3}$ |
| **GREML-GCTA** |  |  |
| | Estimate, $\text{CI}_{95}$ | P-value |
| $h^2_{\text{EA}}, \%$ | 25.9 [23.1; 28.6] | $8.8 \times 10^{-77}$ |
| $h^2_{\text{Migr}}, \%$ | 12.9 [10.2; 15.6] | $2.7 \times 10^{-21}$ |
| $r_g, \%$ | 79.9 [69.9; 89.9] | $2.3 \times 10^{-55}$ |

### Difference in mean PGS_EA_ between cities and ORE accumulated over time

Above, we showed that contemporary migration increases the PGS_EA_ differentiation between Tallinn/Tartu City and ORE. We next set out to explore if this effect accumulated over the last century and if there has been any change in the genetic makeup of migrants over this period of time. We compared the mean PGS_EA_ in Estonians grouped by place of birth and residence and the birth decade, while the PGS_EA_ was adjusted and normalised in the entire Estonian cohort (Figure 7). We used wider birth year bins for the oldest and the youngest participants due to their smaller sample sizes. The comparison between groups of individuals born in Tallinn/Tartu City and ORE shows that individuals born in the cities on average have significantly higher PGS_EA_ than those born in ORE starting from the 1940s (p-value 4.2×10^−3^). Furthermore, the contrast between these groups tends to increase over time (Figure 7A). Consistently, PGS_EA_ is significantly higher in the group of migrants from ORE to the cities than in the group of participants who stayed in ORE. This difference is significant already in the earliest bin (p-value 1.4×10^−3^) and persists in all subsequent bins (Figure 7B). These patterns persist in the analysis of unrelated subsample and with PGS_EA4_ (Supplementary Figures 22 and 28, respectively).

**Figure 7.**
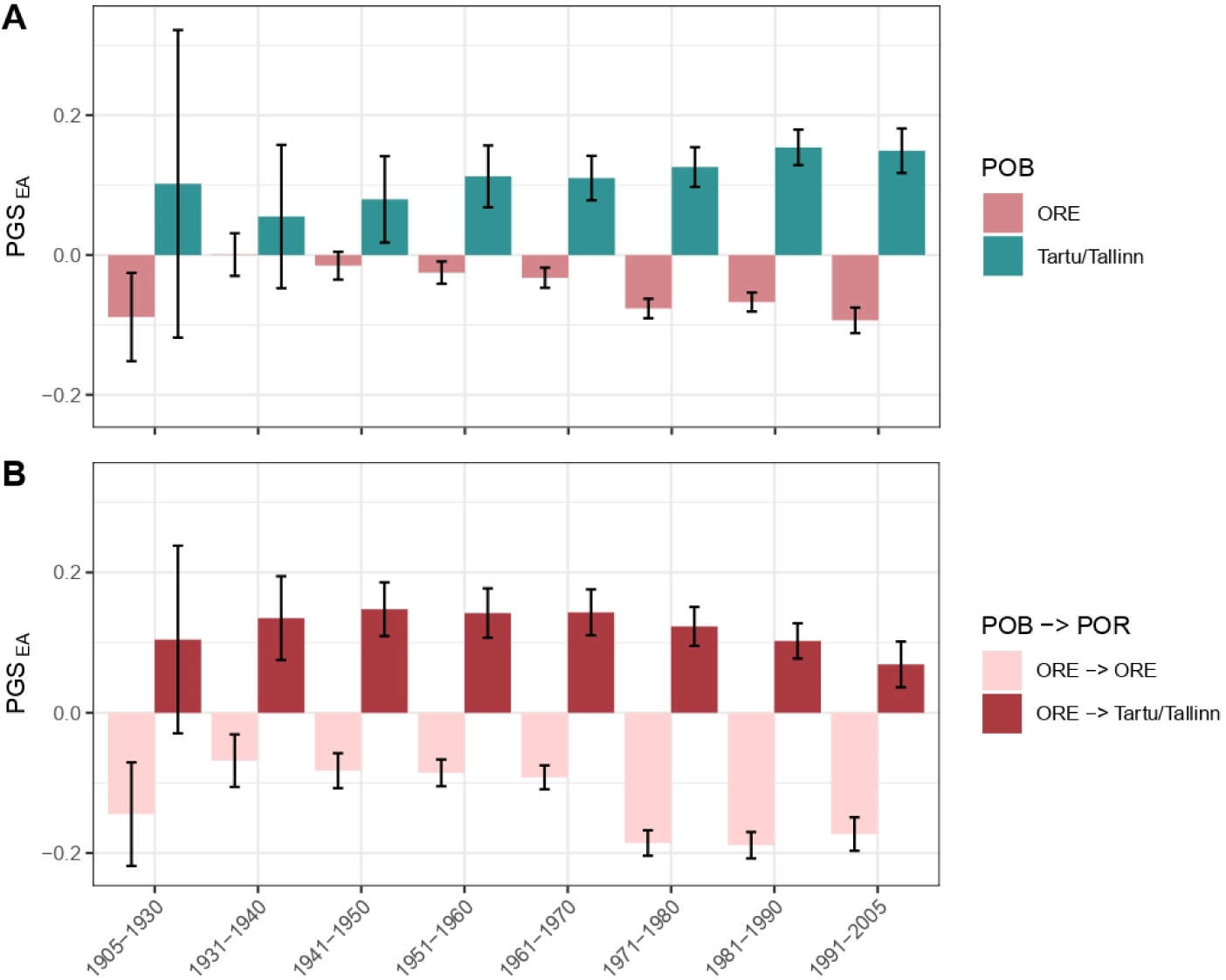
Difference in average PGS_EA_ between cities (Tallinn and Tartu combined) and ORE across birth year bins. (A) Mean PGS_EA_ of individuals born in either ORE or Tallinn/Tartu; (B) mean PGS_EA_ of individuals born in ORE and residing in either ORE or Tallinn/Tartu. PGS_EA_ is adjusted for the top 100 PCs and demographic covariates. Error bars correspond to 95% confidence intervals.

### The relation between genetic factors of educational attainment and migration

It has been previously shown that a higher EA is associated with higher migration activity^45–47^. Hence, the patterns we reported above for PGS_EA_ can merely reflect migration patterns of individuals with various EA levels. This is supported by the observation that EA shows a similar geographic distribution as well as a similar distribution between different migration-profile groups (Supplementary Figures 41-46, 57-68).

To test whether the results for PGS_EA_ can be entirely explained by the trait itself, we first compared PGS_EA_ between different migration groups after controlling for the EA phenotype. With either binary and continuous measures of EA (university degree and years of education, respectively), regressed out of PGS_EA_, the differences between the migration groups become less pronounced but remain significant in most cases (Supplementary Figures 69-82).

We showed above that PGS_EA_ is a significant predictor for migration out of ORE in a logistic regression model. Here, we used logistic regression to test whether PGS_EA_ predicts migration in a joint effect model including EA as a predictor (Table 1). As above, we estimated both population as well as within-sibship effects. Years of education attenuate the regression coefficient of PGS_EA_, yet it remains statistically significant, which is in agreement with the results of another recent study on Swedish twins^48^. Treating EA as a continuous variable doesn’t allow for different effects of different EA categories in the regression model. To relax this condition, we tested an alternative model with EA included as a categorical covariate. In this case, the effect of PGS_EA_ on migration is close to that with years of education as a covariate and is still significant. Remarkably, despite wider confidence intervals and generally lower regression coefficient estimates than at the population level, within-sibship estimates of PGS_EA_ effects remain significant in the joint models.

We also show using GREML-GCTA that migration to the cities has a genetic correlation of 0.8 (CI_95%_: 0.7 - 0.9) with having versus not having a university degree. This suggests the two traits have largely but not fully overlapping genetic backgrounds.

## Discussion

In this work, we harness a sample of more than 180 thousand individuals from the Estonian Biobank^29,30^ to explore the genetic correlates and consequences of contemporary migrations in Estonia. We show that contemporary migrations intensify inter-regional differences in polygenic scores (PGS) for many traits, especially those related to socioeconomic status (SES). The strongest effect is observed for the PGS for educational attainment (PGS_EA_), which is consistent with previous observations in the UK^25^. Moreover, correlation with PGS_EA_ explains a substantial fraction of the inter-regional differences for other PGSs. We demonstrate that spatial PGS differentiation in Estonia is mainly driven by the migration of individuals with relatively high PGS_EA_ to the two largest cities from the rest of the country. Through sibling comparison, we show that such migration and the resulting increase in inter-regional PGS heterogeneity can be in part explained by direct genetic effects. We also demonstrate that the accumulation of inter-regional PGS differences began no later than the mid-20th century and has continued into the 21st century, despite significant societal changes. Our findings shed light on the interplay between genetics and social factors, providing deeper insights into the processes driving contemporary changes in population structure. Furthermore, they illustrate a type of genotype-environment correlation that is likely to be widespread and should be considered in genetic studies.

First, we demonstrate that contemporary migration in Estonia amplifies inter-regional differences in most of the tested PGSs, contrasting with the trend observed for the genome-wide population structure reflected by PCs. PGS_EA_ exhibits the largest increase in *Var_county_* and explains a big fraction of the signal for other PGSs. This motivated us to treat the PGS_EA_ as a genomic variable that most effectively captures genetic loci associated with migration behaviour in the EstBB. The analysis of the geographic distribution of PGS_EA_ and differences between migration groups in PGS_EA_, as well as the comparison of the intensity of the migration paths, indicates that selective migration from ORE to Tallinn and Tartu City is the major process driving the genetic differences between the regions (Supplementary Note 2).

This selective migration might reflect either some causal genetic effects on migration behaviour or be driven by environmental confounding. For example, culturally driven differences in migration rates from different localities of ORE might co-occur with genetic differences between localities due to the recent fine-scale population structure that is challenging to fully adjust for^17^. Indirect parental or dynastic genetic effects, known to particularly influence SES-related traits, may also be considered a form of environmental confounding. However, we accumulate evidence for a substantial role of causal direct genetic effects on migration behaviour. First, PGSs for the SES-related traits are associated with migration behaviour in Estonians and Russians (this study) as well as in the British^25^ populations. Similarly to our observations, in the UK Biobank, PGS for EA is strongly associated with migration from less to more economically developed areas. It is unlikely that the fine-scale genetic structure and migration patterns overlap in the same way between those three populations. Second, and most importantly, we observe consistent patterns when exploring this phenomenon in siblings, where we don’t expect any association between local or family environment and the genotype at birth. Importantly, demonstrating the causal link between genotype and migration behaviour, we still face the problem of genetic confounding. It refers to long-range LD driven mostly by fine-scale population structure and assortative mating^39^. Genetic confounding makes it problematic to highlight certain traits that share direct genetic effects with migration behaviour. Nevertheless, it does not call into question the presence of direct effects. Although the PC76 also has a significant increase in *Var_county_*on the within-sibship level, it may serve as an example of a PC capturing some genetic component with a direct effect on the phenotype^49^. Alternatively, it may be a false positive. We make separate corrections for multiple testing for PCs and PGSs. However, the p-value increases up to 0.054 when correcting jointly for the number of PCs and PGSs tested, while the p-value for PGS_EA4_ (0.027) remains lower than the threshold α = 0.05.

One potential explanation for the observed changes in the population structure may lie in the link between migration and EA. Indeed, there is rich evidence that obtaining an education or applying the acquired qualification is a strong motivation for migration and that education level is associated with migration activity. This might explain part of the association between PGS_EA_ and migration. However, first, these two traits are not perfectly genetically correlated in our sample, thus having some non-overlapping genetic component. Second, in agreement with a study of mobility in Sweden^48^, PGS_EA_ is associated with migration behavior even after including the EA phenotype in both population and within-sibship regression models.

In the 20th century, Estonia went through a series of political transitions related to drastic changes in economic and social organisation^50^. It first gained independence in 1918 and lost it during the Soviet period from 1940 to 1991, which was interrupted by German occupation from 1941 to 1944. Despite this, we observe a consistent trend for increasing differences between the cities and ORE in the PGS_EA_ during almost one century, largely caused by the genetically biased migration from ORE to the cities. This trend is comparable to the genetically selective migration from coal mining areas in the UK. These findings make us suggest that the effect of recent migrations on PGS distribution may be a more general phenomenon for urbanised societies, largely independent of political and economic aspects and probably shared with other countries, at least within Europe. We also replicated the patterns of PGS_EA_ distribution in sex-, age-, and recruitment strategy-based subcohorts and in self-reported Russians, further supporting these patterns to be genuine and general. This hypothesis should be verified using data from more countries, including non-European ones.

PGS_EA_ and the EA phenotype are associated not only with the mere fact of migration but also with migration distance: city residents born in more distant regions have, on average, higher PGS_EA_ and EA than those born in nearby regions. This is consistent with previous studies from other countries^24,51^. A similar pattern has already been observed on the phenotype level in the early 20th century in Estonia, where students from farther away from Tartu City had on average higher scores on an intelligence test than students born closer to the city^52^. Although the test used in that study is considered outdated, factors affecting the result are in line with those currently affecting EA^53^. Since EA is largely transmitted through the family environment, the correlation between EA (and, more broadly, SES) and genetic ancestry, once originated from such selective migration, will persist across subsequent generations^27^.

No matter the relative contribution of causal effects and confounding in the genetic component of the migration behaviour we discuss here, it has clear implications: such migration leads to the differentiation between more and less urbanized and economically developed regions in allele frequencies at a specific set of loci. Thus, such migrations create both active (for the migrating individuals) and passive (for their offspring) genotype-environment correlation^54^. We show that this correlation is not UK-specific, is amplified over generations, and cannot be corrected for either using the top 100 PCs or the whole common SNPs-derived GRM in a LOCO approach. Thus, it might contribute to the inflation of heritability estimates and marginal SNP effects in genetic studies, especially those focusing on socio-economic traits, including EA itself^26,36^. This may also cause inflation in the estimates of assortative mating, confusing it with mating by proximity (Supplementary Note 7)^33^. While here we focus on differences between major cities and other areas, the same phenomenon might be observed on the finer geographic (e.g., neighborhood) level or even on the level of professional or SES groups. The latter would happen even if SES is mostly culturally heritable, but there is some genetic component (not necessarily causally) associated with the chance and direction of SES change. A similar process can create correlations between allele frequencies and SES within cities^27^.

To reduce such confounding on the population level, one would ideally want to control for the parental environment, but at least controlling for place of birth could be a partial solution to the problem^26^. Another solution is to move from population-based cohorts towards large-scale family-based studies to be able to separate the different sources of the genetic associations^55^.

Despite the clear messages delivered by our results, it is important to stress that the present study has several limitations. First, although the biobank data includes information on approximately 20% of the adult population in Estonia, it has been shown not to be a completely representative population cohort^29^ (Supplementary Note 1). Second, the information on the places of birth or residence may contain inaccuracies^56^. The reported place of birth may, in some cases, correspond to the settlement where the maternity hospital was located and not to the actual place where the family lived at the time of birth. For this reason, it is safer to consider counties than individual cities, and our main conclusions are not sensitive to this potential issue. The information on the place of residence is updated regularly, synchronising with the population register. However, people do not always report their movements to the register. Third, in reporting the results for separate age groups, we consider the age of the dead individuals to be fixed at the time of death. Given that migration behaviour can change both with an individual’s age and across historical periods, the desynchronisation of the year of birth and age can obscure some patterns conditioned on either factor. Still, this effect is expected to be negligible (Supplementary Note 1).

Finally, we would like to make a cautionary note about interpreting our results within a broader sociological framework. In most analyses, we used the polygenic score based on population-based GWAS for EA, which captures a substantial fraction of the inter-regional differentiation signal of other PGSs. It has been shown by many studies that it is influenced by lots of diverse confounders^25,26,33,36,44,57–61^. Thus, PGS_EA_ cannot be interpreted as a cumulative genetic factor directly affecting EA outcome. It is rather a correlate of EA, only in part determined by direct genetic effects. One should also note that the differences in mean PGS_EA_ between migration groups are subtle despite being statistically significant. Moreover, the corresponding distributions strongly overlap for all the migration groups considered (Supplementary Note 8).

Our findings demonstrate that people’s geographic mobility, particularly related to urbanisation, is accompanied by changes in the genetic structure of a population. The comparison of Estonia and the UK shows that this phenomenon can manifest in countries with different socio-economic systems as well as population sizes. Such migrations, which are non-random with respect to allele frequencies, generate genotype-environment correlations that should be accounted for in genetic studies.

## Methods

### Participants

The participants of this study were sourced from the Estonian Biobank (EstBB), which is a volunteer-based cohort of the Estonian resident adult population^29,62^. It includes (as of 2022) genetic and diverse phenotype data on 210,438 individuals (72,708 men and 137,730 women) corresponding to ∼20% (∼14% men and ∼24% women) of the contemporary adult population of Estonia^63^. Participants’ age ranges from 18 to 107, determined as of 2022 for alive participants or at the year of death. The EstBB is linked with the Estonian national register, so the information on education level and place of residence is being constantly updated. The participants were recruited over two decades from 2001 to 2021 across the country, covering all the regions and a variety of different settings, providing socio-economic and cultural heterogeneity. Besides genetic and demographic data, participants provided health data, blood samples, and lifestyle information.

### Ethics statement

The activities of the EstBB are regulated by the Human Genes Research Act, which was adopted in 2000 specifically for the operations of the EstBB. Individual level data analysis in the EstBB was carried out under ethical approval “1.1-12/3593” from the Estonian Committee on Bioethics and Human Research (Estonian Ministry of Social Affairs), using data according to release application “4-1.6/GI/79” from the Estonian Biobank.

### Genotypes and quality control

Samples were genotyped on the Infinium Global Screening Array (GSA) of different versions (depending on the time of recruitment) with approximately 550,000 overlapping positions. Samples with <95% call rate or mismatch between genetic and self-reported sex were excluded. Before the imputation step, all non-SNP polymorphisms and strand-ambiguous SNPs were filtered out. The final number of SNPs before the imputation step was 309,258. The genotypes were imputed with Beagle 5.4^64^ using the Estonian Reference panel as a reference set^65^. To create polygenic scores, we extracted a set of 1,075,599 autosomal HapMap 3 SNPs with a minor allele count >5 and info score >0.7. Unrelated individuals were defined as having less than 2nd-degree relationship inferred with KING^66^.

For GREML analysis, the non-imputed genotyping data were used after keeping SNPs with minor allele frequency >0.01, Hardy–Weinberg equilibrium (HWE) p-value >10^−5^ and missingness <0.015. Related individuals with a 2nd-degree relationship and closer were excluded. Relationships were inferred with KING^66^.

### Ancestry and PCA

Genetic ancestry grouping was estimated using imputed genotypes with bigsnpr, following the original workflow^67^. For ancestry inference, genotypes were imputed using 1000 Genomes Project phase 3 samples^3^. Individuals from ‘Europe (East)’, ‘Europe (North West)’, and ‘Finland’ inferred ancestry groups were kept for further analysis. Next, individuals with self-reported ethnicity other than ‘Estonian’ or ‘Russian’ were excluded from the participants who passed the genetic ancestry filter. These steps were implemented to retrieve a relatively genetically homogeneous set of participants. In total, 183,576 individuals (63,753 men and 119,823 women) left after the filtering. Next, the resulting set was subdivided into self-reported Estonians (172,376) and Russians (11,200). The purpose of the latter step was to divide the sample by cultural and historical background rather than by genetic profile^68^.

A principal component analysis (PCA) was conducted separately for self-reported Estonians (182,252 individuals) and Russians (17,954 individuals) to capture genetic structure within the corresponding groups. Before the analysis, genotypes were filtered for minor allele frequency >0.01, Hardy–Weinberg equilibrium (HWE) p-value >10^−5,^ and missingness <0.05. Long-range linkage disequilibrium regions were removed^69^. Genotypes were pruned for linkage disequilibrium with PLINK2^70,71^ with a window size of 50kb, a step 5kb, and r^2^ threshold 0.1. The PCA to construct PCs on self-reported Estonian and Russian individuals was conducted on this SNP set using flashPCA version 2^72^.

### Polygenic score calculations

Polygenic scores were computed for 169 phenotypes using population-based GWAS summary statistics from the UK Biobank (PGSs)^28^. The PGSs were calculated using summary statistics from GWAS in the European ancestry cohort of the UK Biobank conducted by the Pan-UKBB project^32^. The Pan-UKBB project particularly presents an analysis of 7,228 phenotypes, spanning 16,131 studies. The list of traits selected for the analysis included the maximally independent set of 146 phenotypes (with correlation between them <0.1) for which GWAS results passed the quality control. The link to this list is available on the Pan-UKBB project website (https://pan.ukbb.broadinstitute.org/downloads/index.html). Additionally, 23 phenotypes related to education, mental health, fluid intelligence, height, and body mass index (BMI) were added. The complete list of the phenotypes and the numbers of individuals included in the study is presented in Supplementary Table 1. An additional PGS was calculated using summary statistics from the largest GWAS of educational attainment (EA) currently available with Estonian individuals and the 23andMe cohort excluded from the meta-analysis^33^.

Polygenic scores were calculated using SBayesR (gctb_2.02) with default parameters (--pi 0.95,0.02,0.02,0.01 --gamma 0.0,0.01,0.1,1 --chain-length 10000 --burn-in 2000 --out-freq 10), including an LD matrix built using data on 50,000 UK Biobank participants^73^. To reduce the effect of the ancestral genetic structure on polygenic scores, the top 100 principal components (PCs) specific to the Estonian or Russian cohort were regressed out. Sex, age, sex×age, and age^2^ were also regressed out of the PGSs to mitigate the influence of potential sex and age bias reported for population volunteer cohorts^58,74^. In analyses of PGS adjusted for educational attainment, binary or continuous EA (see *Educational attainment phenotypes* in the methods) was also regressed out. PGS for EA (PGS_EA_ or PGS_EA4_) was also regressed out from other PGSs where explicitly mentioned.

### Sources of education and geographic information

Initial information on the highest level of education, place of birth and place of residence was obtained from the questionnaire completed by participants when enrolled in the biobank. The EstBB regularly synchronises its information with the Estonian Population Register on the highest level of education and the municipality of current residence. The data used in this study was last updated in 2022. Participants without information on the counties of birth and residence in Estonia or born outside the country were excluded from the analysis. Participants born or residing in Harju or Tartu Counties and lacking information on the municipality were excluded from the analyses, where it was necessary to distinguish Tallinn/Tartu City from other municipalities of the corresponding counties. After filtering, the analysed sample included 172,376 self-reported Estonians and 11,200 self-reported Russians.

### Educational attainment phenotypes

Continuous and binary traits corresponding to educational attainment were considered. The continuous ‘years of education’ phenotype was derived according to the ISCED 2011 methodology. The link table for the reported level of education, ISCED 2011, and ‘years of education’ is presented in Supplementary Table 3. Alternatively, attainment of a Bachelor’s degree or higher was used as a binary phenotype (0 - not having a Bachelor’s degree; 1 - having a Bachelor’s degree or higher). The quantitative EA phenotype was adjusted to mitigate possible sampling bias in the corresponding analyses: sex, age, sex×age, and age^2^ and 100 genetic PCs were regressed out using linear regression.

### Inter-regional differences in PC values and polygenic scores

To measure the inter-regional differences in a variable of interest, we calculated the proportion of variance explained by county differences:

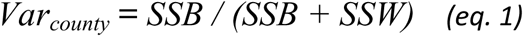

where *SSB* is the sum of squares between counties, and *SSW* is the sum of squares within counties. P-values were calculated from the ANOVA test. The chi-square test was implemented to test whether the difference of variance explained by county of birth and county of residence together is significantly larger than by only one of them. The base model to compare with was a less powerful model with either county of birth or county of residence as an independent variable. Statistical significance was determined using a level of 0.05 after the Bonferroni correction for the number of tests (100 for PCs, 169 for PGSs, or 170 when PGS_EA4_ was included).

### Sibling analyses

The sibling design was applied in the analysis of inter-regional variation of PCs and PGSs, the comparison of migration groups and the association between PGSs and migrations from ORE. Siblings were defined by KING^66^ criteria of first-siblings and an additional threshold of 0.177 < Kinship < 0.354. Only siblings born in the same county were included in the analysis. The total number of siblings after filtering was 41,081 making up 19,407 sibships. PGSs were first adjusted as described above. Then, the sibship mean PGS was subtracted from each individual’s PGS. There is evidence that birth order may affect EA^75^. It could lead to systematic differences in PGS_EA_ between older and younger siblings through participation bias mechanisms. However, we do not observe such significant differences in our data and thus do not adjust PGSs for birth order.

As by design (all the siblings from every family have the same POB), all the county averages and, accordingly, *Var_county_* for POB are zero for all the PGSs and PCs. Because of this characteristic of the model, *Var_county_* does not follow the distribution of F-statistic from the standard ANOVA. To determine the PCs and PGSs significantly exceeding the *Var_county_* expected by chance, we generated 10,140 random variables *v ∼ N(0,1)* and applied the sibling design to them. The estimate of the p-value was obtained as (*r+1)/(n+1)*, where *n* is the number of simulated variables and *r* is the number of these variables that produce *Var_county_* statistic greater than that calculated for the actual PC or PGS^76,77^. The number of variables used was chosen as a compromise between precision and performance as it allowed us to get a p-value = 0.05 after Bonferroni correction for the PGS with *Var_county_* exceeding all but two values of the random variables (3/10,141*169 = 0.05).

### PGS effects on migration from ORE to the cities

Other Regions of Estonia (ORE) are defined as the counties of Estonia, excluding Harju and Tartu Counties, which encompass the largest cities, Tallinn and Tartu, respectively. The migration phenotype was specified for individuals born in ORE with POR in ORE or Tallinn/Tartu City. The phenotype value 0 corresponds to POR also in ORE, the value 1 corresponds to POR Tallinn or Tartu City.

For the estimation of the population effects of the PGSs, we used mixed (*eq. 2*) and fixed (*eq. 3*) effects logistic regression models:

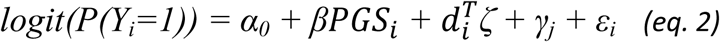

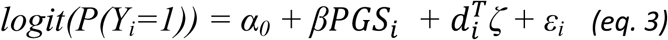

Here *Y_i_* is the phenotypic outcome of an individual *i*. *P(Y_i_=1)* is the probability of the outcome *Y_i_*being 1. is the individual’s polygenic score. *α_0_* is the intercept and *β* is the regression coefficient of the *PGS*. *δ* is the vector of fixed coefficients for the demographic variables s*ex* and *age*, denoted by vector *. γ_j_* is the random intercept with *γ_j_ ∼ N(0,σ^2^_γ_)*, for sibship *j* accounting for the between-family variation in the intercept. The residual is represented as *ε_i_* with *ε_i_ ∼ N(0,σ^2^_ε_)*. The mixed effects model was applied to the sample of siblings. The fixed effects model was applied to the subsample of siblings with a single individual picked up randomly from each of the sibships, as well as to the subsample of unrelated Estonians.

For the estimation of the within-sibship effects of the PGSs, we used mixed effects logistic regression similar to the models introduced by <u>(43)</u> and <u>(26)</u> (*eq. 4*) and fixed effects logistic regression without random effect of sibship (*eq. 5*):

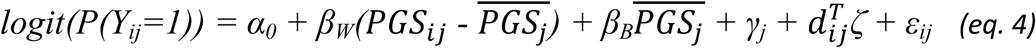

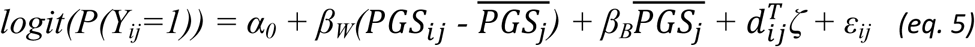

*PGS_ij_* denotes the individual’s polygenic score, and 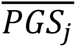 refers to the mean polygenic score of the family *j*. *β_W_* and *β_B_* correspond to within- and between-family PGS effects, respectively. All the other terms correspond to those from the population model.

The joint regression analysis of PGS_EA_ and EA was performed in accordance with the formulas given above with EA (years of education or categories) added as a covariate.

### Heritability and genetic correlation calculations

Bivariate GREML analysis implemented in GCTA software^78,79^ was used to estimate heritabilities and genetic correlations for EA and migration from ORE to Tallinn or Tartu City. For this 43,334 individuals with relatedness more distant than the 2nd-degree were used. Sex, age, age^2^, sex×age, sex×age^2^ and 10 genetic PCs were included as covariates in the models. The heritability estimates were transformed from the observed to the liability scale using the Robertson transformation^80^.

### Geographic data visualisation

Shapefiles used to plot maps of Estonia with county borders were retrieved from the Estonian Land Board website (Administrative and Settlement Division, 2023.02.01)^81^. Geographic data were visualized in R^82^ with the aid of the following packages: ‘sf’^83,84^, ‘geos’^85^ and ‘ggplot2’^86^.

## Data and Code Availability

Access to the Estonian Biobank data (https://genomics.ut.ee/en/content/estonian-biobank) is restricted to approved researchers and can be requested. UK Biobank summary statistics are available through the Pan-UKBB project website (https://pan.ukbb.broadinstitute.org/downloads/index.html). Custom R code used for statistical analyses is available on GitHub (https://github.com/ivkuz/GeneticMigrationStructureEstonia).

## Author contributions

IK, LP, FM, and VP conceived and designed the study. IK performed all the analyses. IK and VP wrote the initial draft of the manuscript. All co-authors contributed to the interpretation of the results, reviewed, and approved the submitted version of the manuscript.

## Competing Interest Statement

The authors declare no competing interests.

## Acknowledgements

We want to acknowledge the participants of the Estonian Biobank. This study was funded by the European Union through the European Regional Development Fund Project No. 2014-2020.4.01.15-0012 GENTRANSMED and by the Ministry of Education and Research of Estonia through the project TK214 Centre of Excellence for Personalised Medicine. Data analysis was carried out in part in the High-Performance Computing Center of the University of Tartu. This project has received funding from the European Union’s Horizon Europe research and innovation programme under grant agreement No 101060011. Views and opinions expressed are however those of the authors only and do not necessarily reflect those of the European Union or European Research Executive Agency. Neither the European Union nor the granting authority can be held responsible for them. IK was a student of and supported by the Center of Life Sciences, Skolkovo Institute of Science and Technology, Moscow, Russia, during the initial phase of the project. VP and MM were supported by the European Union through Horizon 2020 research and innovation programme under grant no 810645 and through the European Regional Development Fund project no. MOBEC008. MM was also supported by the Estonian Research Council grant PUT (PRG1899). UV was supported by the Estonian Research Council grant PUT (PSG759). LP was supported by the Italian Ministry of University and Research (2022B27XYM). FM was supported by Fondazione con il Sud (2018-PDR-01136) and by the Italian Ministry of University and Research (2022P2ZESR). The funders had no role in study design, data collection and analysis, decision to publish or preparation of the manuscript. This work was written at writing retreats organized by the University of Tartu Institute of Genomics and the Estonian Doctoral School for Natural and Agricultural Sciences (2021-2027.4.04.24-0003), co-funded by the European Union.

## Notes

### Competing Interest Statement

The authors have declared no competing interest.

### Summary of Updates

In this revised manuscript, we addressed potential confounding due to population structure and improved the interpretation of the results. One major change is the addition of a within-family analysis of siblings born in the same county, with polygenic scores (PGSs) adjusted for the average PGS of their sibship. Within-family associations between PGSs and migration patterns are consistent with those from the main population-based analysis, reducing concerns about residual population structure and providing evidence for direct genetic effects on migration behavior.

