## Supplementary Material for "Genetic effects on migration behaviour contribute to increasing spatial differentiation at trait-associated loci in Estonia"

Ivan A. Kuznetsov, Estonian Biobank Research Team, Mait Metspalu, Uku Vainik, Luca Pagani, Francesco Montinaro, Vasili Pankratov

### Supplementary Note 1. Estonian Biobank cohort overview

The Estonian Biobank (EstBB) cohort is a volunteer-based sample of the Estonian resident adult population (aged ≥18 years)[1,2]. EstBB participants are genotyped and deeply phenotyped. The phenotype information includes medical records, as well as self-reported medical, lifestyle, demographic, socio-economic, and geographical data. The current number of participants exceeds 210,000 which corresponds to approximately 20% of the contemporary adult population of Estonia. Compared to the UK Biobank[3] it has a more even representation of different age groups and different regions of the country. All of this makes EstBB a good dataset for independent replication of previous findings and novel analyses.

The EstBB Project was initiated in 1999. Since then two main waves of recruitment were conducted (Supplementary Figure 1A). Approximately 52,000 individuals were recruited until 2016[1] and three times more after 2016[2]. These two periods substantially differed in the recruitment strategy. During the first stage, the participating proposals were mainly spread by general practitioners to their patients. The second stage of recruitment was conducted through a wide-scale advertising campaign.

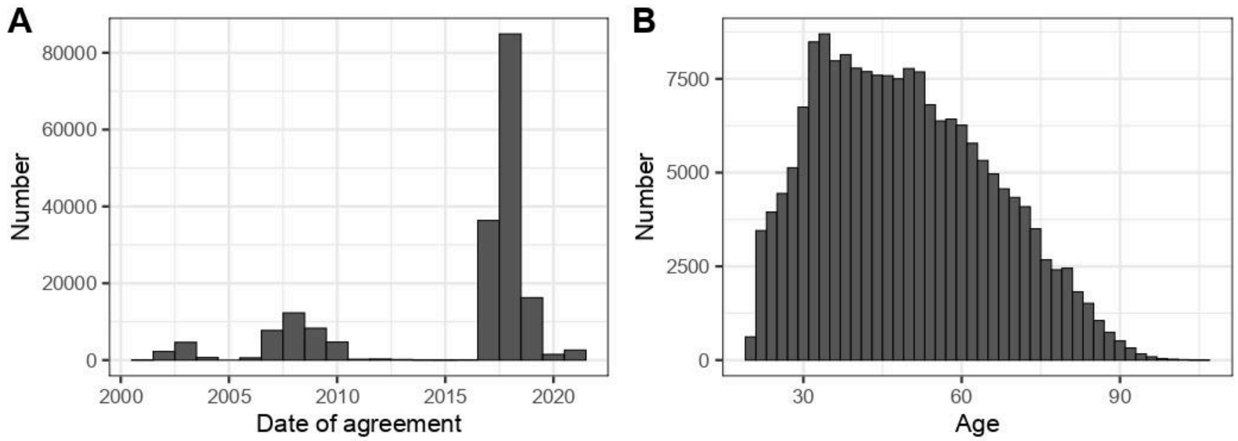

**Supplementary Figure 1. The distribution of EstBB participants after filtering by (A) date of agreement (year of recruitment) and (B) age.**

The data in the biobank is being regularly updated. Information on the level of education and the place of residence is synchronised with the population register database, therefore we used current age or age of death in this study which is almost equal to using a year of birth for separate-generation analysis. The proportion of dead participants is 2.3%. For only 0.2% of the participants, the age group (“18-24”, “25-48”, “49-64”, “65 and older”) mismatches with the corresponding birth year group (“1998-2004”, “1974-1997”, “1958-1973”, “1957 and earlier”), calculated as the difference between 2022 and corresponding boundaries of the age bins. Thus, the effects of such mismatches are expected to be negligible. There are no age restrictions in the recruitment process besides being 18 or older. As a consequence, a full range of ages are covered by the cohort with the oldest participant being 105 (Supplementary Figure 1B).

There are two major ethnic groups (defined based on self-reported ethnicity) currently living in Estonia: Etonians (920,000; 69% of all the people residing in Estonia) and Russians (315,000; 24%), based on the 2021 Population census[4]. The EstBB cohort covers both groups, although Russians are underrepresented in comparison to Estonians. However, the administrative regions are covered relatively uniformly by participants from these groups with 13-31% of Estonians sampled and 1-7% of Russians (Supplementary Figure 2) from each county. Here we use the data only on the participants of self-reported Estonian or Russian ethnicity, excluding individuals who indicated both. Throughout the study, we focused mainly on the Estonian group because of the statistical power available and replicated some analyses in the Russian group.

For the sake of the robustness of the findings as well as for analysis of differences between subgroups, we divided the overall sample into two cohorts based on self-reported ethnicity: Estonians and Russians. The Estonian cohort was further divided into several partially overlapping groups: unrelated individuals (excluding one individual from each pair of closer than 2nd-degree relationship), males, females, Estonians of age 18 - 24, 25 - 48, 48 - 64, 65 and older, and two groups by the period of recruitment (agreement): 2001-2016 and 2017-2021 (Supplementary Figure 3).

We repeated most of our analyses in the above-mentioned subgroups of the Estonian cohort and in the Russian cohort as well as on the entire Estonian cohort but using polygenic scores based on summary statistics from a within-sibship GWAS (sPGS, Supplementary Note 3)[5]. We also conducted analyses of the distribution of EA phenotypes across regions and migration directions for subgroups within the Estonian cohort, as well as for the Russian cohort. The order of these results coincides with the structure of the main text. Detailed descriptions of the analyses can be found in the Results and Methods section of the main text. For each type of analysis where all the subsample categories are presented, we followed the order: “entire Estonian cohort”, “Russian participants”, “unrelated Estonian participants”, “Estonian subgroups by sex”, “Estonian subgroups by age”, “Estonian subgroups by year of joining the biobank”. Where applicable, results of the same analysis with $sPGS_{EA}$ are presented.

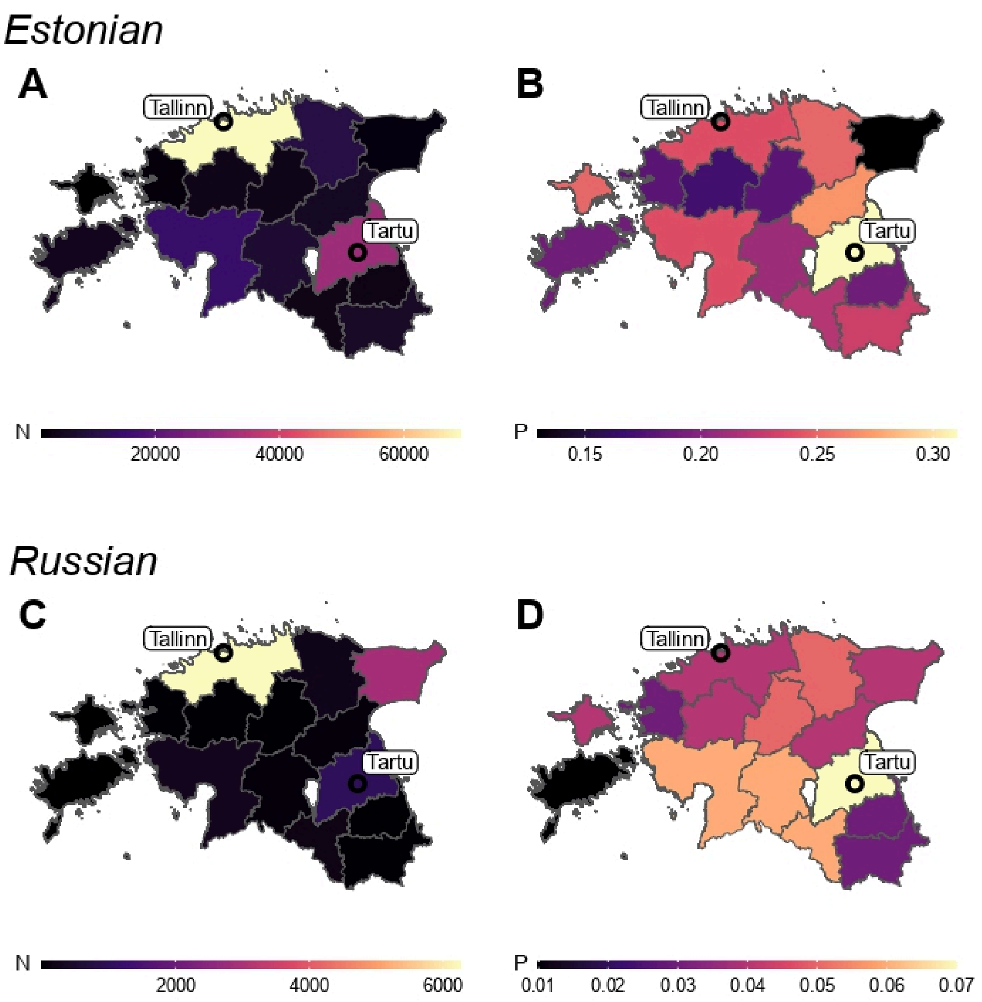

**Supplementary Figure 2. Geographic distribution of EstBB participants by county of residence.** (A) The number of EstBB participants of self-reported Estonian ethnicity ; (B) the fraction of EstBB participants among residents of self-reported Estonian ethnicity; (C) the number of EstBB participants of self-reported Russian ethnicity; (D) the fraction of EstBB participants among residents of self-reported Russian ethnicity. Data on the number of current residents per county was taken from the 2021 Population census[4].

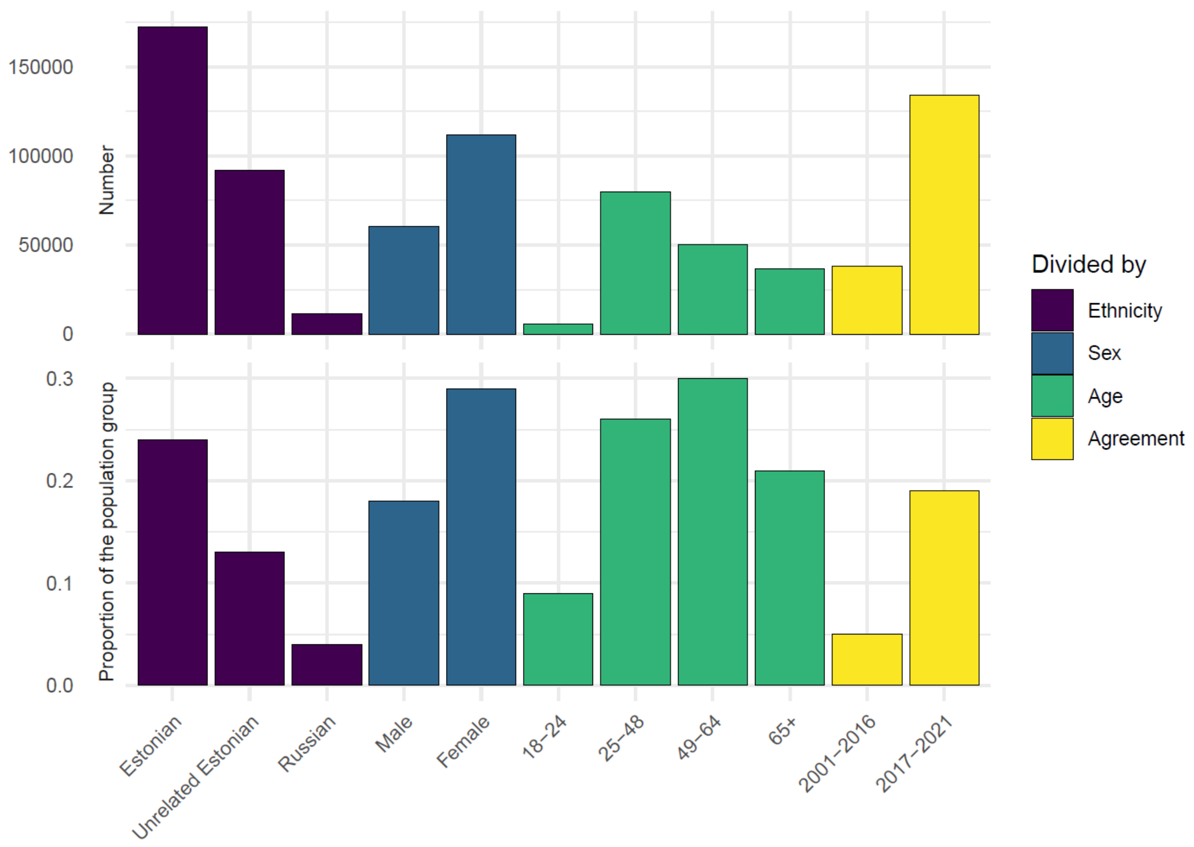

**Supplementary Figure 3. Absolute number of participants in each group (top) and the same, normalized by the size of the corresponding group in the general population (bottom).** In the bottom panel, the proportions of "Estonians", "Unrelated Estonians" and the groups by the year of agreement (yellow) are shown relative to number of adult self-identified Estonians in the population. Census population data was taken from the 2021 Population census[4].

#### Supplementary Note 2. Intensity of the internal migration in Estonia

The migration flows between the counties of Estonia are not in equilibrium (Supplementary Figure 4). The highest net migration is to Tallinn, the capital and largest city of Estonia. The second highest net migration is to Tartu, the major city in Southern Estonia and the largest educational centre in the country.

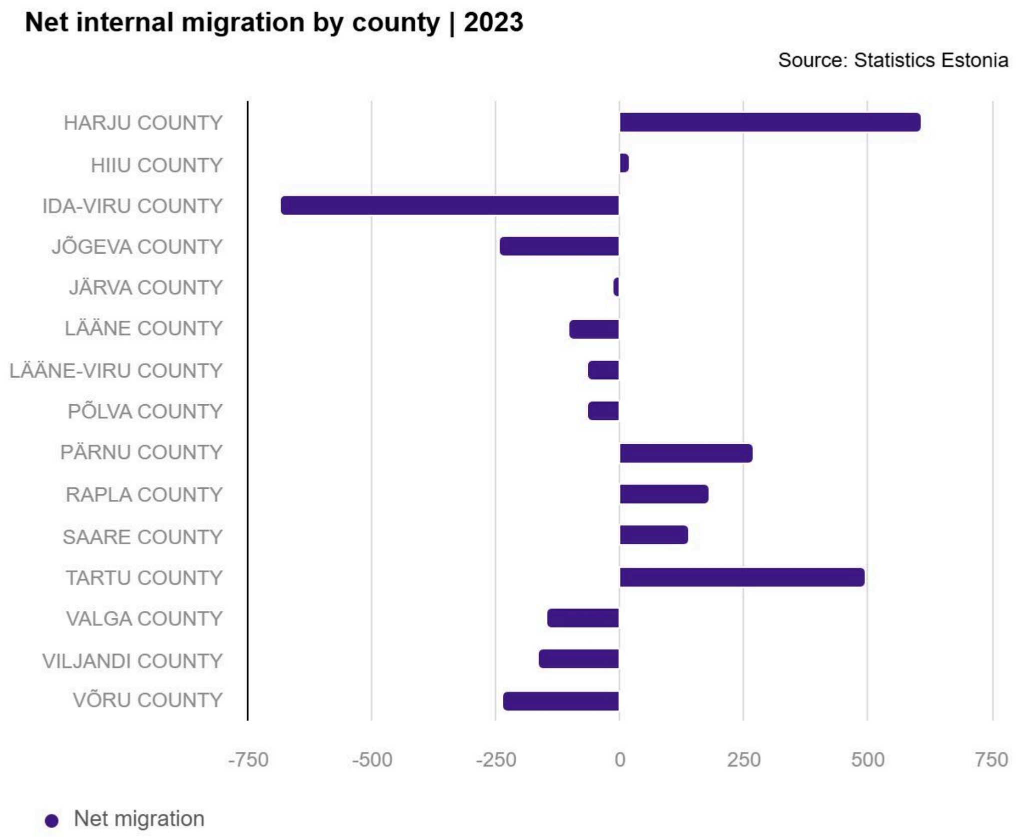

**Supplementary Figure 4.** Net internal migration in Estonia by county in 2023. Source: Statistics Estonia[6].

To access migration flows between counties we utilised the information from both the 2021 Census and the EstBB. Mainly, we used the EstBB as a source of information on the migration flows between the counties. However, the sampling density is not uniform across the country, with Tartu County being the most overrepresented region (Supplementary Figure 2). Without adjustment for the differences in the sampling density, we expect an overestimation of such migration directions as Tartu County. To correct for this effect, we introduced county-specific weights for the study participants:

$$w_i = \frac{n_i}{N} \cdot \frac{N^{EBB}}{n_i^{EBB}},$$

Where $w_i$ is the weight for a study participant residing in a county $i$, $n_i$ is the number of individuals residing in the county $i$ according to the 2021 Census, $N$ is the number of individuals residing in Estonia according to the 2021 Census, $n_i^{EBB}$ is the number of the study participants residing in the county $i$, $N^{EBB}$ is the overall number of the study participants. We calculated these weights separately for self-reported Estonian and Russian adults. The resulting weighted migration matrices show the fractions of individuals born in each of the counties who stayed in the county of birth or moved to any other county (Supplementary Figure 5).

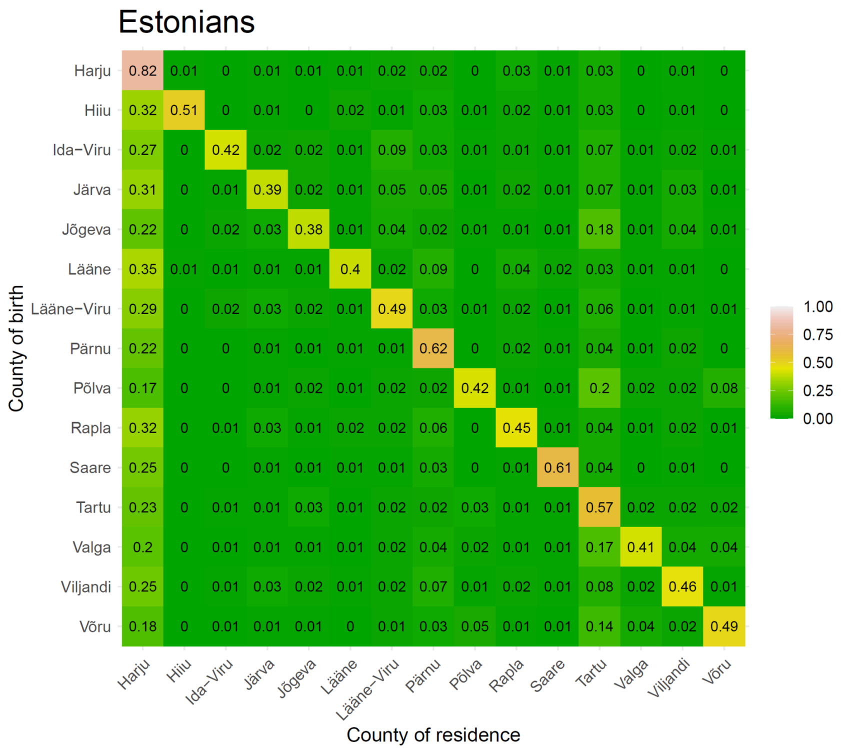

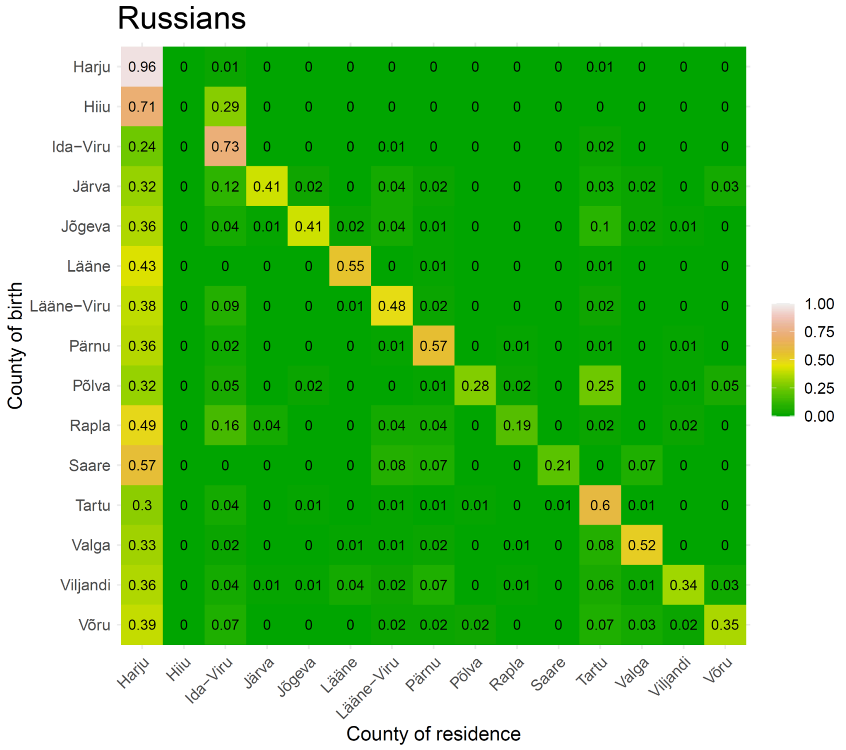

**Supplementary Figure 5. Internal migration in Estonia.** The fractions of individuals of Estonian (top) and Russian (bottom) self-reported ethnicity born in each of the counties who stayed in the county of birth or moved to another county.

#### Supplementary Note 3. Polygenic scores based on within-sibship GWAS summary statistics (sPGS)

Besides the 169 UKB-based PGSs and $PGS_{EA4}$, we calculated polygenic scores for 24 phenotypes using within-sibship GWAS summary statistics (sPGSs)[5]. The sPGSs were calculated for 24 out of 25 phenotypes analysed in the original study presenting a set of within-sibship GWAS results estimating direct genetic effects. The sPGS for C-reactive protein was excluded from the analysis due to an issue with the MCMC procedure in SBayesR when working with the respective summary statistics . Supplementary Table 2 lists 24 traits with corresponding sample sizes. The methodology for calculating sPGS was the same as for the population-based PGSs (see Methods for details on PGS calculation).

sPGSs are less prone to population confounding, however, not necessarily represent direct genetic effects[7,8]. sPGSs are also substantially less powerful than population PGSs. For example, sPGS explains only 0.438% of EA variance in comparison with 5.4% for $PGS_{EA}$ and 7.1% $PGS_{EA4}$ in the set of unrelated Estonians. We conducted the analyses of $Var_{county}$, geographic distribution (for all the subsamples) and comparison of the migration groups (for the main Estonian subsample) with sPGSs. We note again that the sPGSs are not the estimates of the direct genetic effects on the respective traits. Thus, these analyses are meant to be the sensitivity tests as the underlying summary statistics come from a different source, contain more statistical noise but are less confounded.

Within-sibship GWAS summary statistics are available on OpenGWAS (https://gwas.mrcieu.ac.uk/), see the original study[5] for details.

### Supplementary Note 4. Robustness of $Var_{county}$ estimates

The main analysis of the $Var_{county}$ of the PCs in the full Estonian sample shows that a significant fraction of the variance of all the top 100 PCs can be explained by POB and it is always higher than the fraction explained by POR. The same analysis for the PGSs shows that a significant fraction of the variance of most of the tested PGSs can be explained by POR. The fraction of variance explained by POR is higher than the fraction explained by POB for most of the cases when there is a significant difference between them. The highest $Var_{county}$ for POB and POR is demonstrated by $PGS_{EA}$.

In this note, we demonstrate that neither participation bias nor the residual population structure are likely sources of the observed patterns. First, we present the analysis of $Var_{county}$ for PCs, PGSs and sPGSs in the subgroups described in Supplementary Note 1. Second, we explore how the number of PCs used for the $PGS_{EA}$ adjustment affects the results. Third, we analyse the adjustment of $PGS_{EA}$ for the complete genetic relatedness matrix (GRM) in a leave-one-chromosome-out (LOCO) approach.

#### Analysis of principal components and polygenic scores in subgroups

It has been shown that population-based cohorts tend to be biased, which can make generalization to the entire population uncertain. Among the characteristics most prone to bias are sex, age, SES[9,10]. Here, we present the analysis of $Var_{county}$ for PCs, PGSs and sPGSs in the subgroups described in Supplementary Note 1. Briefly, we split the overall sample into subsamples by sex, age, year of recruitment (the corresponding groups differ in their average EA of the participants). We also conduct the analysis for the subcohort of self-reported Russian participants, which can be considered as an independent replication as the geographic distribution as well as historical background of this group is substantially different in comparison with the self-reported Estonian group. The analyses conducted on the subcohort of unrelated Estonians is presented in Supplementary Note 5, Supplementary Figure 17. The results for the subsamples and for the Russian subsample are consistent in general. The comparison of the Estonian subcohorts shows that an increased rate of participation in the EstBB among certain subgroups in the population is unlikely to be the reason of the observed patterns, as this pattern is reproduced in the different groups. Despite the relatively small sample size of the Russian subsample, it also demonstrates patterns in their main aspects replicating the observations in the main Estonian sample. The analysis of sPGS is less stable due to the high proportion of statistical noise in the sPGS which is especially notable in small samples. The $Var_{county}$ values presented in the plots can be found in Supplementary Table 10.

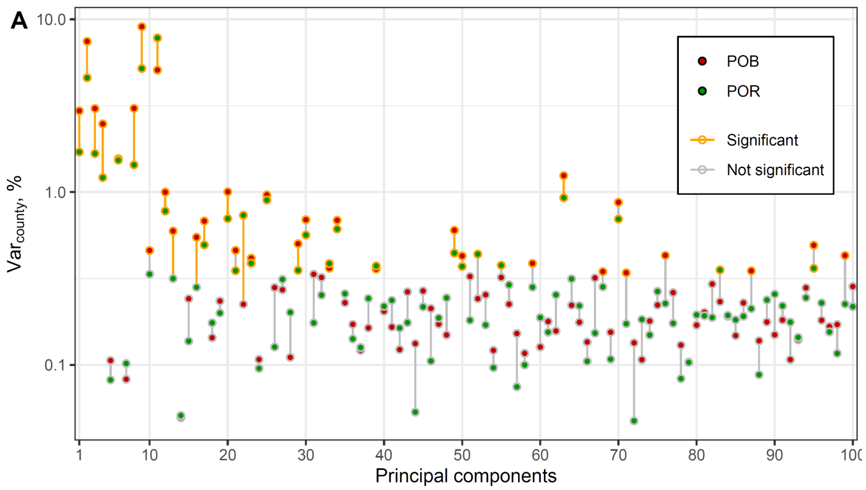

**Supplementary Figure 6. Inter-individual variance of PCs among Russian participants, explained by county of birth (POB) and county of residence (POR)**. The PCs are derived from the Russian subcohort. Red and green dots refer to the POB and POR, correspondingly. Estimates significantly different from zero are outlined in yellow. The line connecting the two points is yellow when the variance explained by POB and POR together is significantly larger than the variance explained by only the weaker predictor. The significance level is 0.05, adjusted for 100 tests with Bonferroni correction.

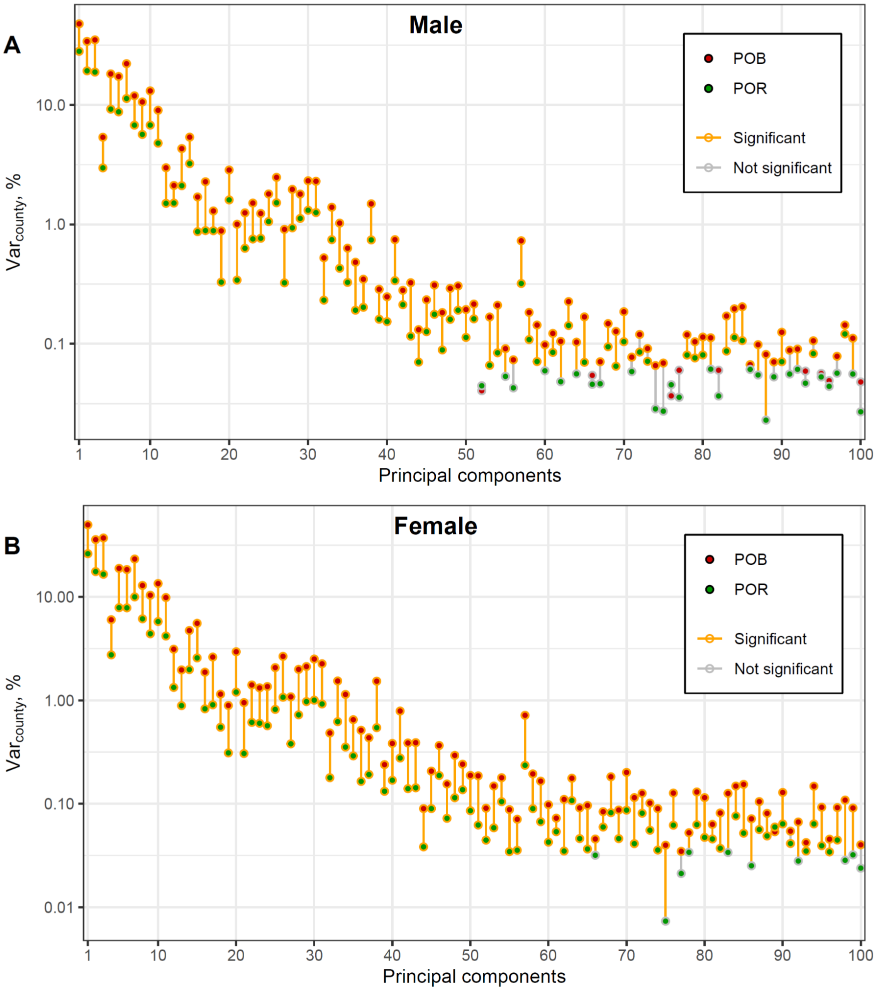

**Supplementary Figure 7. Inter-individual variance of PCs among (A) male and (B) female Estonian participants explained by county of birth (POB) and county of residence (POR)**. Red and green dots refer to the POB and POR, correspondingly. Estimates significantly different from zero are outlined in yellow. The line connecting the two points is yellow when the variance explained by POB and POR together is significantly larger than the variance explained by only the weaker predictor. The significance level is 0.05, adjusted for 100 tests with Bonferroni correction.

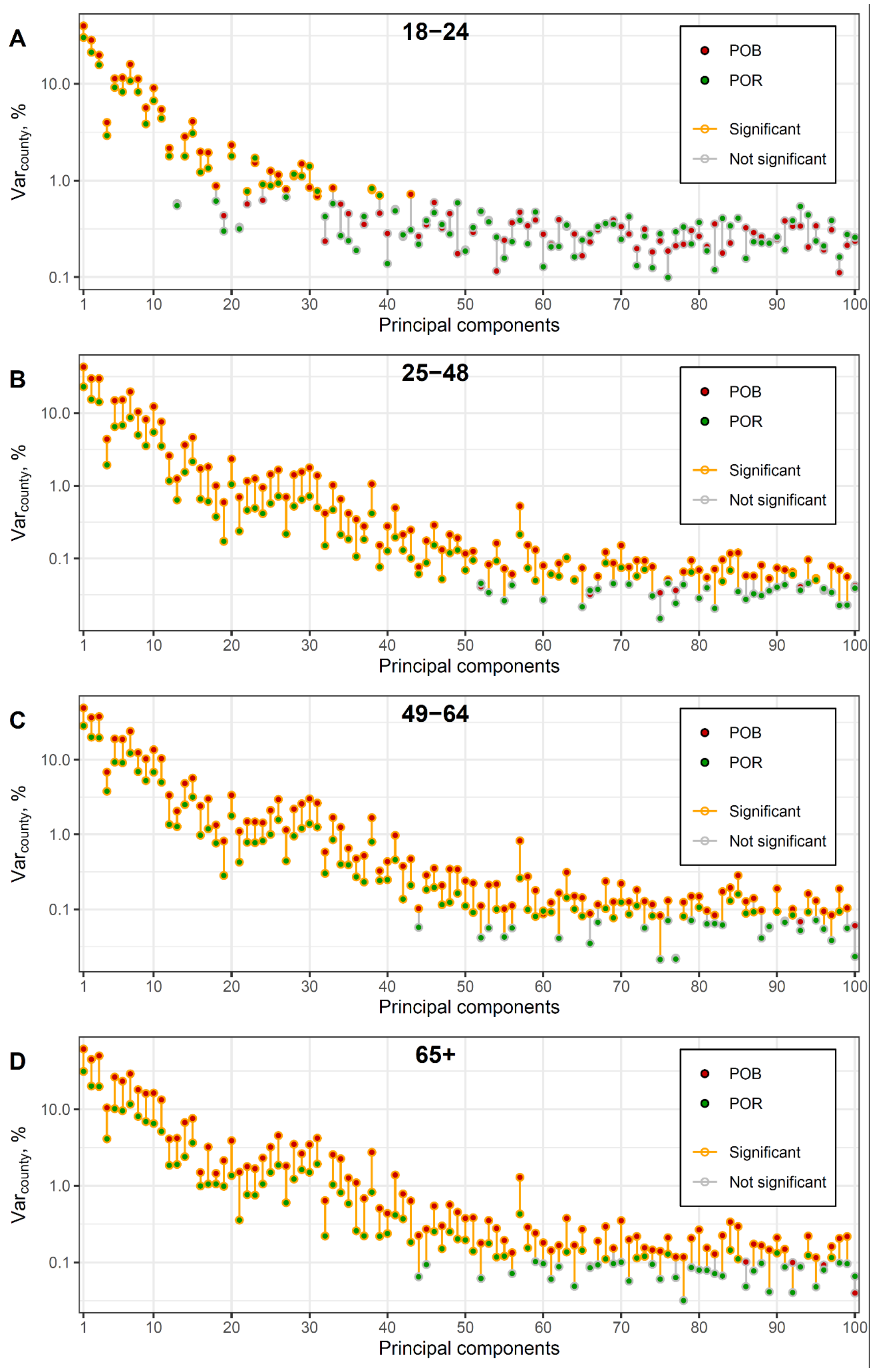

**Supplementary Figure 8. Inter-individual variance of PCs among Estonian participants stratified by age explained by county of birth (POB) and county of residence (POR)**. Age groups were defined as (A) 18-24, (B) 25-48, (C) 49-64, (D) 65+. Red and green dots refer to the POB and POR, correspondingly. Estimates significantly different from zero are outlined in yellow. The line connecting the two points is yellow when the variance explained by POB and POR together is significantly larger than the variance explained by only the weaker predictor. The significance level is 0.05, adjusted for 100 tests with Bonferroni correction.

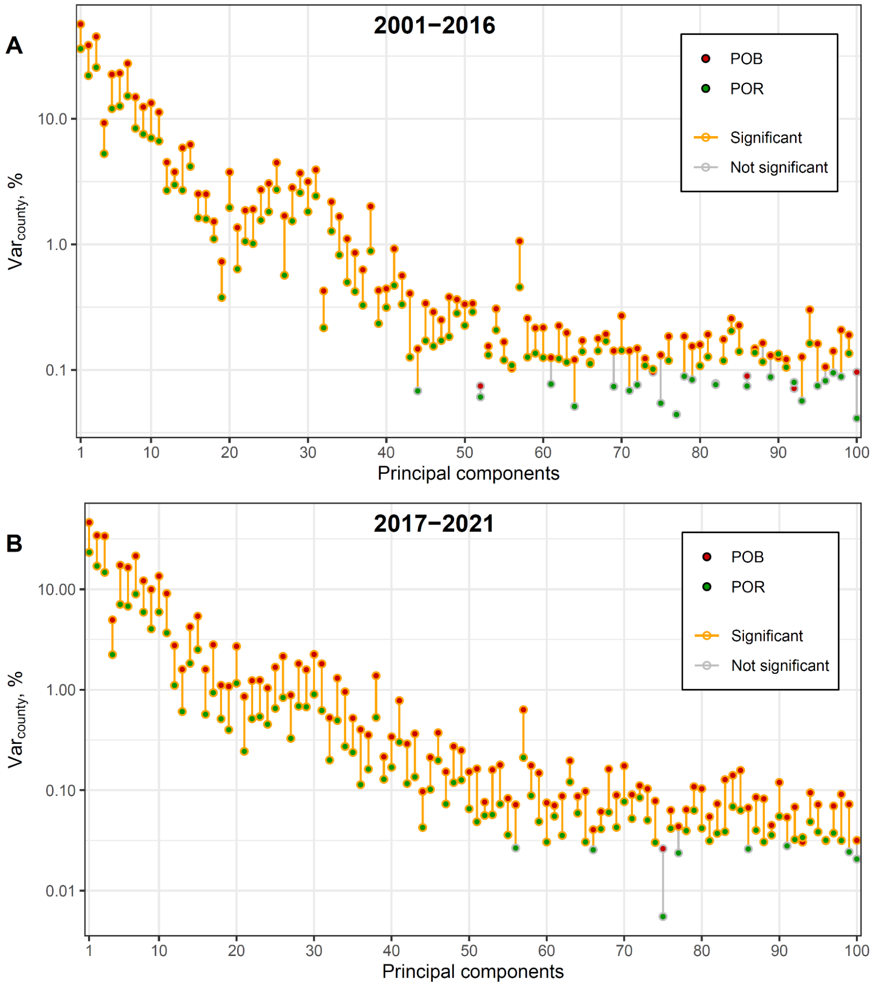

**Supplementary Figure 9. Inter-individual variance of PCs among Estonian participants stratified by year of joining the biobank explained by county of birth (POB) and county of residence (POR)**. The periods of joining are (A) 2001-2016 and (B) 2017-2021. Red and green dots refer to the POB and POR, correspondingly. Estimates significantly different from zero are outlined in yellow. The line connecting the two points is yellow when the variance explained by POB and POR together is significantly larger than the variance explained by only the weaker predictor. The significance level is 0.05, adjusted for 100 tests with Bonferroni correction.

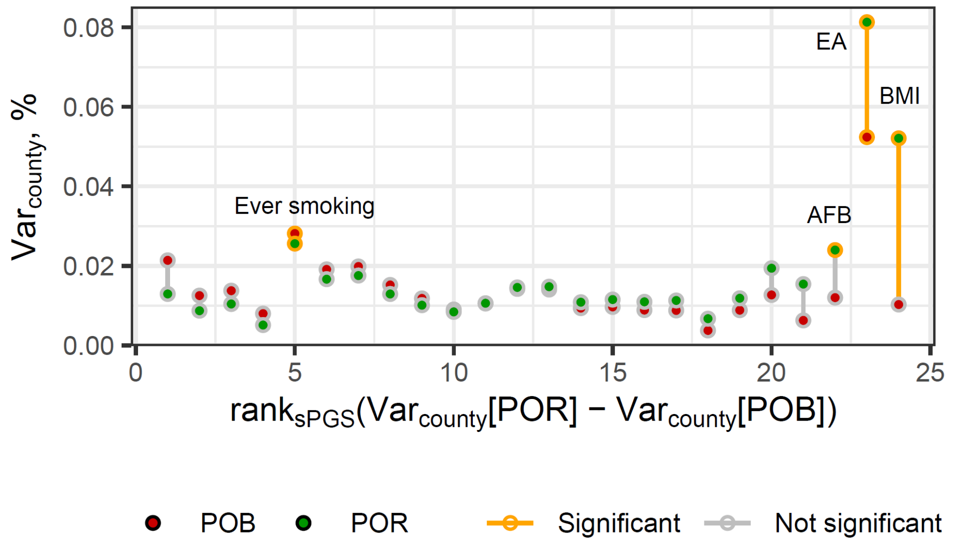

**Supplementary Figure 10. Estimates of the inter-individual variance of sPGSs among Estonian participants explained by POB and POR.** sPGSs are adjusted for demographic and genetic ancestry covariates. sPGSs are ordered according to the rank of difference between $Var_{county}$ for POR and POB. Estimates significantly different from zero are outlined in yellow and labeled (*BMI* - Body Mass Index; *EA* - educational attainment; *AFB* - age at first birth). The line connecting the two points is yellow when the variance explained by POB and POR together is significantly larger than the variance explained by only the weaker predictor. The significance level is 0.05, adjusted for the number of sPGS tested with Bonferroni correction.

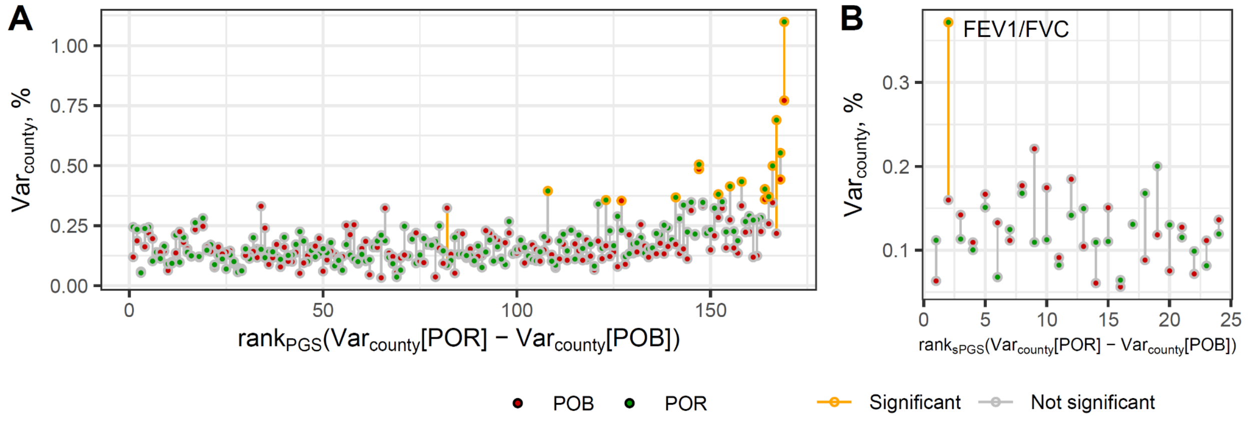

**Supplementary Figure 11. Estimates of the inter-individual variance of (A) PGSs and (B) sPGSs among Russian participants explained by POB and POR.** (s)PGSs are adjusted for demographic and genetic ancestry covariates. PGSs and sPGSs are ordered according to the rank of difference between $Var_{county}$ for POR and POB in the full Estonian subsample (as in Figure 1B and Supplementary Figure 10 accordingly). Estimates significantly different from zero are outlined in yellow. The line connecting the two points is yellow when the variance explained by POB and POR together is significantly larger than the variance explained by only the weaker predictor. The significance level is 0.05, adjusted for the number of (s)PGS tested with Bonferroni correction.

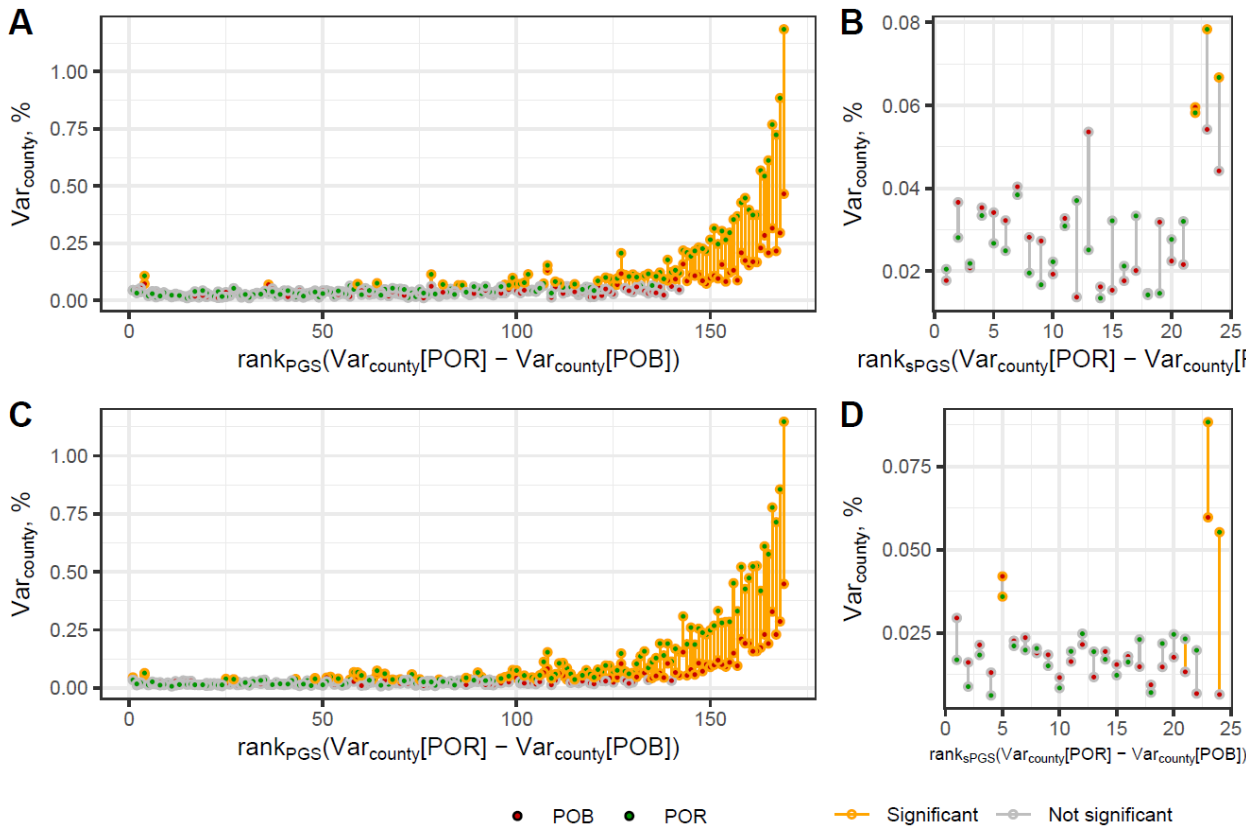

**Supplementary Figure 12. Estimates of the inter-individual variance of (A, C) PGSs and (B, D) sPGSs among (A-B) male and (C-D) female Estonian participants explained by POB and POR.** (s)PGSs are adjusted for demographic and genetic ancestry covariates. PGSs and sPGSs are ordered according to the rank of difference between $Var_{county}$ for POR and POB in the full Estonian subsample. Estimates significantly different from zero are outlined in yellow. The line connecting the two points is yellow when the variance explained by POB and POR together is significantly larger than the variance explained by only the weaker predictor. The significance level is 0.05, adjusted for the number of (s)PGS tested with Bonferroni correction.

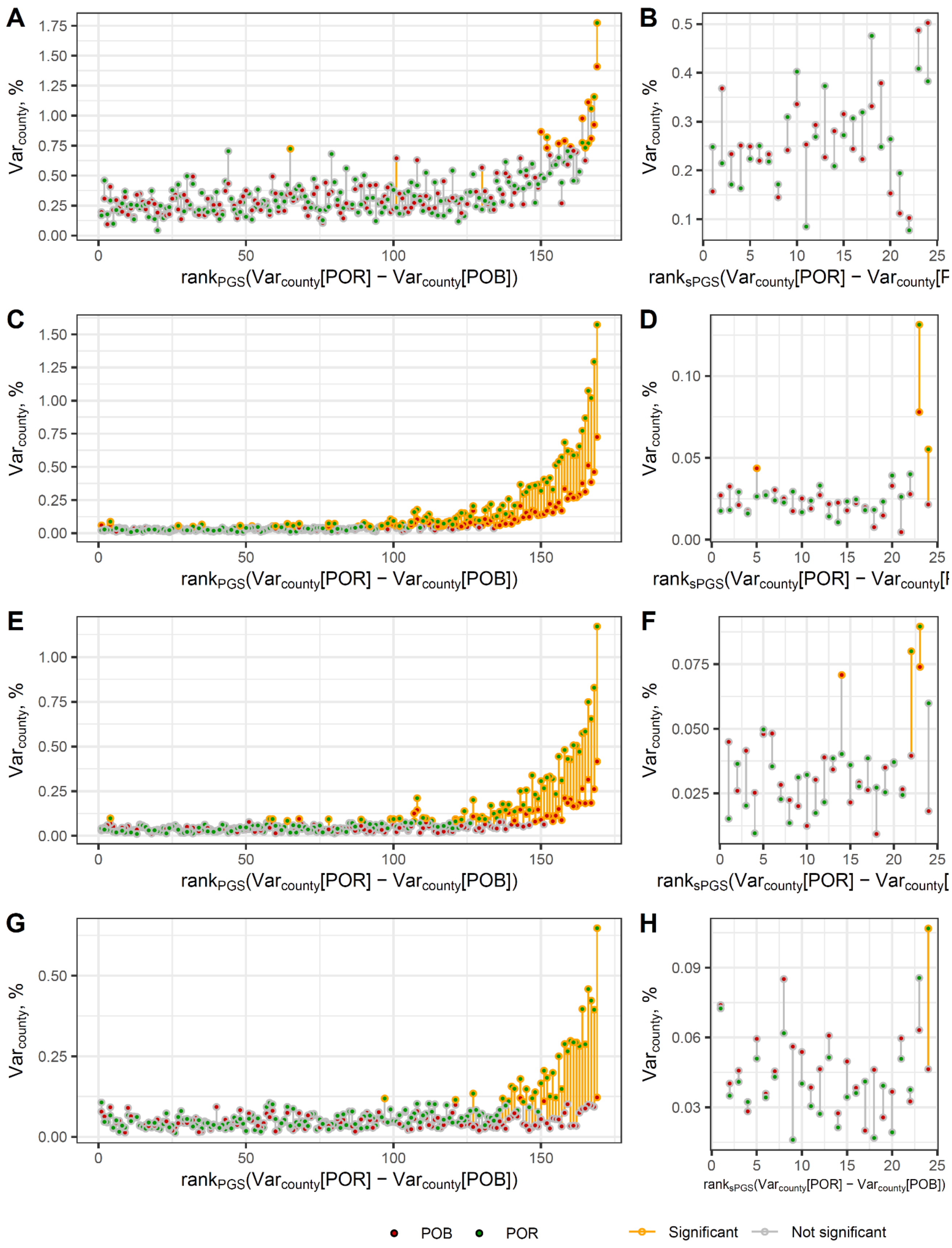
A
B
C
D
E
F
G
H
Var_county, %
rank_PGS(Var_county[POR] − Var_county[POB])
rank_sPGS(Var_county[POR] − Var_county[POB])
POB
POR
Significant
Not significant

**Supplementary Figure 13. Estimates of the inter-individual variance of (A, C, E, G) PGSs and (B, D, F, H) sPGSs among Estonian participants stratified by age explained by POB and POR.** Age groups were defined as (A-B) 18-24, (C-D) 25-48, (E-F) 49-64, (G-H) 65+. (s)PGSs are adjusted for demographic and genetic ancestry covariates. PGSs and sPGSs are ordered according to the rank of difference between $Var_{county}$ for POR and POB in the full Estonian subsample. Estimates significantly different from zero are outlined in yellow. The line connecting the two points is yellow when the variance explained by POB and POR together is significantly larger than the variance explained by only the weaker predictor. The significance level is 0.05, adjusted for the number of (s)PGS tested with Bonferroni correction.

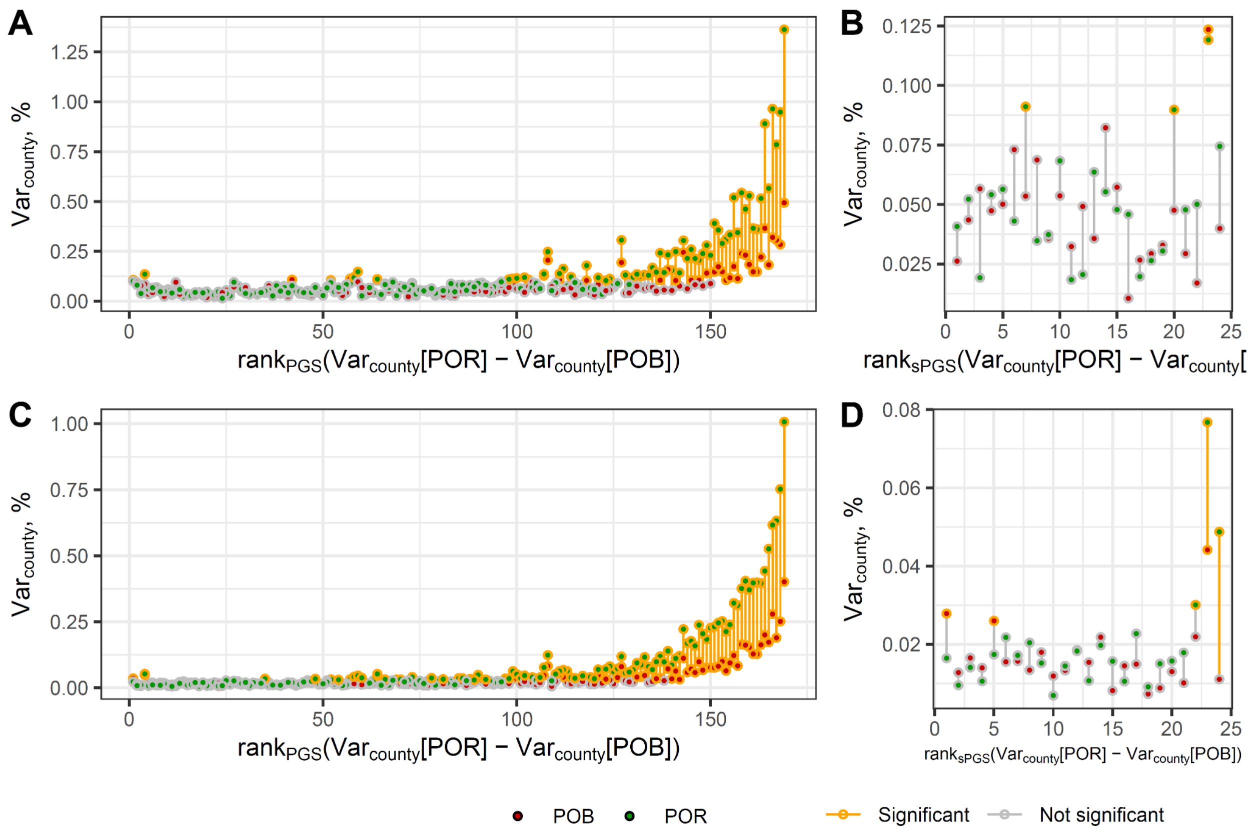

**Supplementary Figure 14. Estimates of the inter-individual variance of (A, C) PGSs and (B, D) sPGSs among Estonian participants stratified by year of joining the biobank explained by POB and POR.** The periods of joining are (A-B) 2001-2016 and (C-D) 2017-2021. (s)PGSs are adjusted for demographic and genetic ancestry covariates. PGSs and sPGSs are ordered according to the rank of difference between $Var_{county}$ for POR and POB in the full Estonian subsample. Estimates significantly different from zero are outlined in yellow. The line connecting the two points is yellow when the variance explained by POB and POR together is significantly larger than the variance explained by only the weaker predictor. The significance level is 0.05, adjusted for the number of (s)PGS tested with Bonferroni correction.

#### Cumulative adjustment of $PGS_{EA}$ for the top 100 PCs

Across the work, we adjust the PGSs for the top 100 PCs of the genetic relatedness matrix built on common SNPs to mitigate the effects of population genetic structure. Adjustment for 100 PCs is substantially more than usually used in the PGS analyses[11–14]. To demonstrate how adjustment for different numbers of PCs affects the $Var_{county}$ of $PGS_{EA}$ we calculated it for POB and POR with the $PGS_{EA}$ adjusted for the top N principal components, where N ranges from 0 to 100 (Supplementary Figure 15). $PGS_{EA}$ was also adjusted for demographic covariates (sex, age, sex×age and $age^2$) in all the models. The set of 92,110 unrelated self-reported Estonian individuals was used. The greatest change in $Var_{county}$ is observed within $N < 5$ while more PCs added have only minor effects, especially when $N > 40$. The difference between $Var_{county}$ for POR and POB increases within $N <= 10$. After a moderate decrease from N = 11 to N = 15, it is rather stable and fluctuates between 0.75 and 0.76% which is more than 0.05% higher than before the adjustment. Note, that with 92,110 PCs (the number equal to the number of individuals), $PGS_{EA}$ can be explained completely. Thus a gradual decline of $Var_{county}$ for POB and POR as well as of the difference between them is expected. To conclude, the adjustment for the PCs decreases the $Var_{county}$ of $PGS_{EA}$ but increases the difference between $PGS_{EA}$ $Var_{county}$ for POR and POB. The most substantial effect is observed with adding roughly the first 10-15 PCs while adding further PCs has a minor effect.

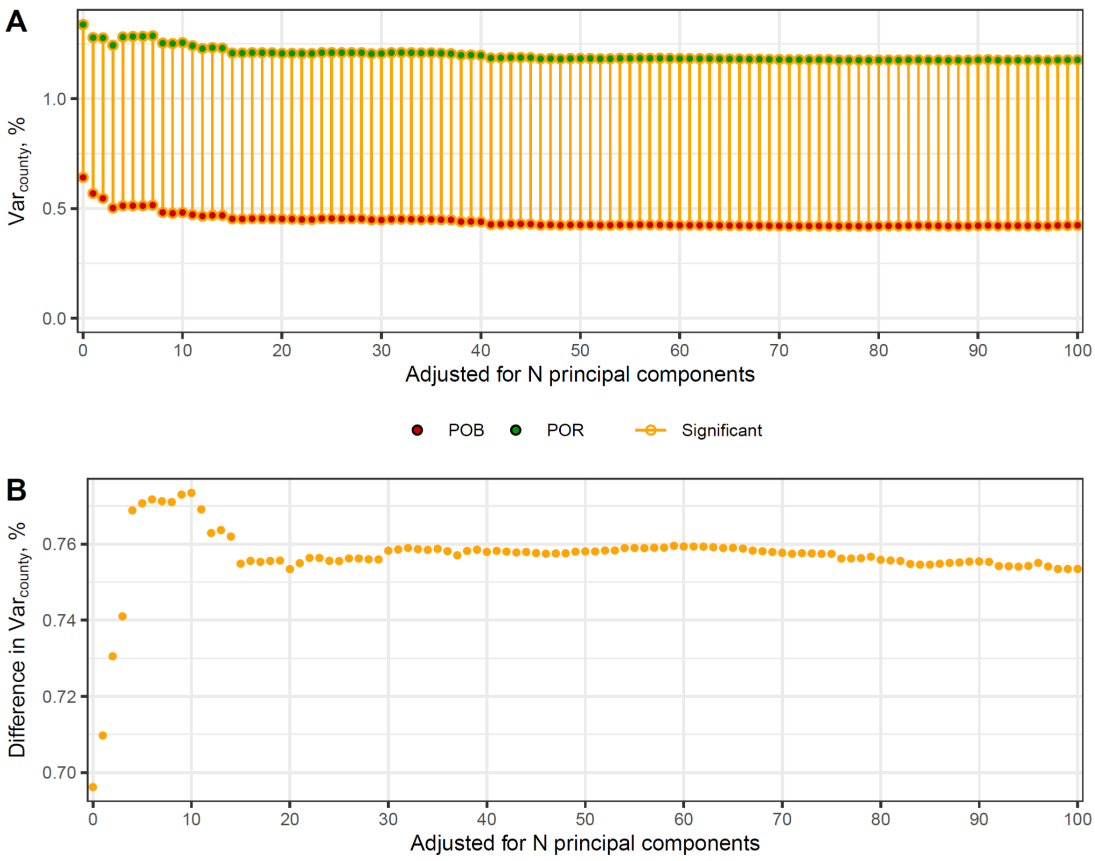

**Supplementary Figure 15. (A) Fraction of the inter-individual variance of $PGS_{EA}$ cumulative adjusted for the top 100 PCs, explained by county of birth (POB) and county of residence (POR) and (B) the difference between $Var_{county}$ for POR and POB.** $PGS_{EA}$ is also adjusted for demographic covariates in all the models. Red and green dots refer to the POB and POR, correspondingly. Estimates significantly different from zero are outlined in yellow. The line connecting the two points (in A) and the points (in B) are yellow when the variance explained by POB and POR together is significantly larger than the variance explained by only the weaker predictor (if significant) or when the stronger predictor is significant. The significance level is 0.05, after Bonferroni correction for 169 tests.

#### Adjustment of $PGS_{EA}$ for the complete genetic relatedness matrix (GRM) in a leave one chromosome out (LOCO) approach

Top 100 principal components (PCs) capture only a fraction of the population structure captured by the Genetic Relationship Matrix (GRM). An alternative approach to correcting a PGS for population structure is to use a Linear Mixed Model (LMM), which incorporates the entire Genetic Relationship Matrix (GRM). This approach has the potential to mitigate the effects of population structure on the PGS to a greater extent than using only the top PCs. However, regressing out too many PCs in a fixed-effect framework can lead to overcorrection[15]. In the limit, when the number of PCs equals the number of individuals, all the variance in the PGS can be explained.

To investigate how adjusting $PGS_{EA}$ for different proxies of population genetic structure influences its $Var_{county}$, we compared the $Var_{county}$ of $PGS_{EA}$ corrected for the top 10 or 100 PCs, the GRM, or a combination of both in the set of unrelated individuals adjusted. To minimize proximal contamination, we employed the leave-one-chromosome-out (LOCO) approach.

GRMs were calculated on imputed genotypes using LDAK software version 5.1[16]. The imputed genotypes (as described in Methods) were filtered using PLINK2 to retain only biallelic single nucleotide polymorphisms (SNPs) with a minor allele frequency (MAF) >0.01 and a Hardy–Weinberg equilibrium (HWE) p-value $>10^{-5}$. Next, SNPs were thinned using a squared correlation ($r^2$) threshold of 0.98 within a 100-kb window. Per-chromosome GRMs were calculated assuming LDAK model, assuming equal weights and scaling parameter (--power) of -0.25. These per-chromosome GRMs were then merged into 22 LOCO-GRMs.

Using LDAK, we fit the Restricted Maximum Likelihood (REML) models for per-chromosome $PGS_{EA}$. In these models, a respective LOCO-GRM was included as a random effect predictor, while fixed-effect covariates included demographic variables (sex, age, sex×age, $age^2$) and 10 genetic PCs. The independent residuals from these models were combined into a single PGS. This final PGS was either used as is ("demography, 10 PCs, GRM" in Supplementary Figure 16) or further adjusted for 100 PCs ("demography, 100 PCs, GRM" in Supplementary Figure 16).

We compared the $Var_{county}$ of $PGS_{EA}$ for POB and POR adjusted only for the demographic factors, adjusted for demographic variables and 10 or 100 PCs, and adjusted for demographic variables, 10 or 100 PCs, and the GRM (Supplementary Figure 16). In both cases, adjustment for genetic PCs makes $Var_{county}$ for POB and POR lower. Additional adjustment of $PGS_{EA}$ for GRM makes $Var_{county}$ even lower, however, this decrease is minor: the difference in $Var_{county}$ with and without adjustment for the GRM ranges from 0.01% to 0.03%. It likely reflects the fact that the top 10 PCs already capture most of the structure affecting the $Var_{county}$ of $PGS_{EA}$. Adjustment for the top 100 PCs in addition to the top 10 PCs in the presence or absence of the GRM has a larger effect on the $Var_{county}$ (0.05-0.08%). It might show that adjustment for PCs is more effective than for GRM in the LOCO approach.

Regardless of the mechanisms, different approaches to adjustment of the $PGS_{EA}$ for the genetic structure lead to only minor changes in $Var_{county}$. The adjustments decrease $Var_{county}$ for both POB and POR and have almost no effect on the difference between them. Thus, neither PC adjustment nor GRM adjustment substantially impacts the signal of non-random geographic distribution of $PGS_{EA}$ or the amplification of this signal resulting from contemporary migrations.

Both the PCA and GRM approach we applied here relied on the information on the common SNPs. It was shown that the recent population structure may be better captured by rare polymorphisms[17]. As the samples we work with here are only genotyped for common SNPs and using imputed genotypes for rare variants can potentially suffer from low accuracy, we abstained from using this approach. Using IBD segments can aid in capturing recent population structure. However, no single correction method fully eliminates residual population structure. Instead, to exclude all the potential effects of the population genetic structure and also of the parental (dynastic) effects, we used a within-sibling design (see the Main text).

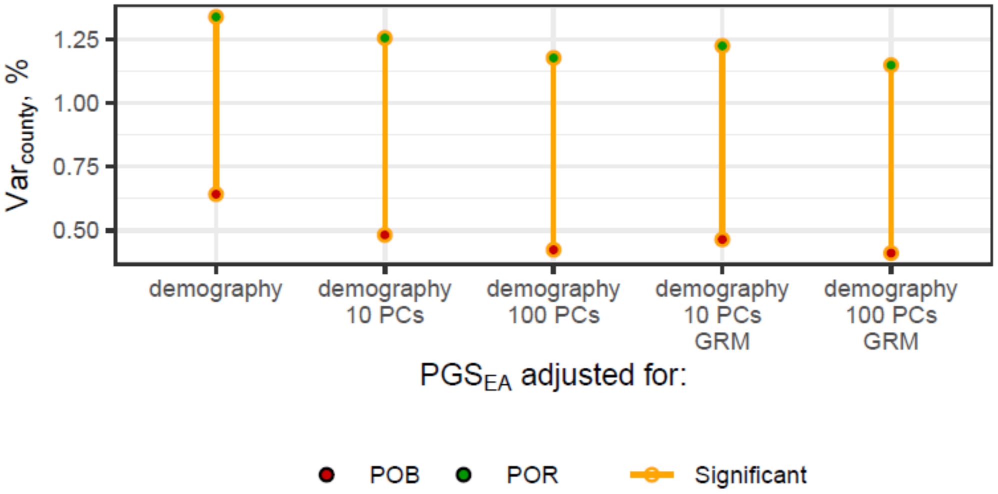

**Supplementary Figure 16. Fraction of the inter-individual variance of $PGS_{EA}$ adjusted for PCs and GRM, explained by county of birth (POB) and county of residence (POR).** $PGS_{EA}$ is also adjusted for demographic covariates in all the models. Red and green dots refer to the POB and POR, correspondingly. Estimates significantly different from zero are outlined in yellow. The line connecting the two points are yellow when the variance explained by POB and POR together is significantly larger than the variance explained by only the weaker predictor (if significant) or when the stronger predictor is significant. The significance level is 0.05, after Bonferroni correction for 169 tests.

### Supplementary Note 5. Replication of the main results in unrelated Estonian individuals

Most of the statistical tests we use in this study require the independence of data points in the sample. In the main analyses, we assume that genetic relatedness in the sample does not affect the results substantially. This assumption is based on the population-based recruitment strategy of the EstBB and a substantial fraction of the overall population presented in the analysed sample (Supplementary Note 1). The former fact makes our estimates close to a description of the population. However, we note that for robust statistical inferences a sample with independent observations should be used. For this, we repeat all the main analyses in the sample of Estonian individuals with relatedness more distant than the 2nd-degree ("unrelated Estonian subsample/individuals"), assuming their independence.

All the results from the analyses with the subset of unrelated Estonians are consistent with the inferences from the overall Estonian sample. The key findings remain statistically significant.

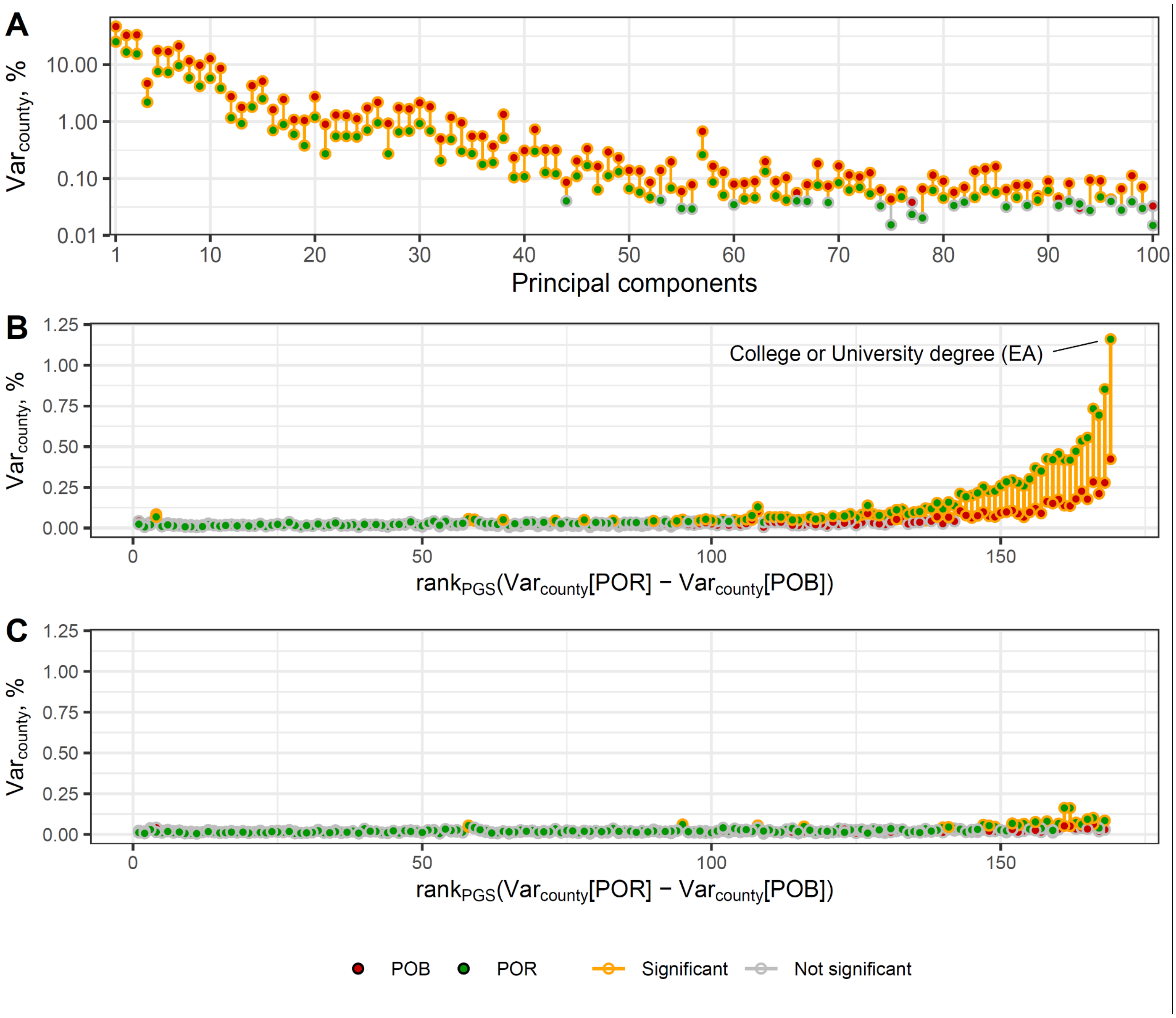

**Supplementary Figure 17. Fraction of the inter-individual variance of (A) PCs, (B) PGSs and (C) PGSs additionally adjusted for $PGS_{EA}$, explained by county of birth (POB) and county of residence (POR) among unrelated Estonian participants**. PGSs are preliminary adjusted for the top 100 PCs and demographic covariates. The y-axis in panel A has a logarithmic scale. The PGSs on the x-axis in panels B and C are ordered according to the rank of difference between $Var_{county}$ for POR and POB in the full Estonian subsample (as in Figure 1B). Red and green dots refer to the POB and POR, correspondingly. Estimates significantly different from zero are outlined in yellow. The line connecting the two points is yellow when the variance explained by POB and POR together is significantly larger than the variance explained by only the weaker predictor (if significant) or when the stronger predictor is significant. The significance level is 0.05, after Bonferroni correction.

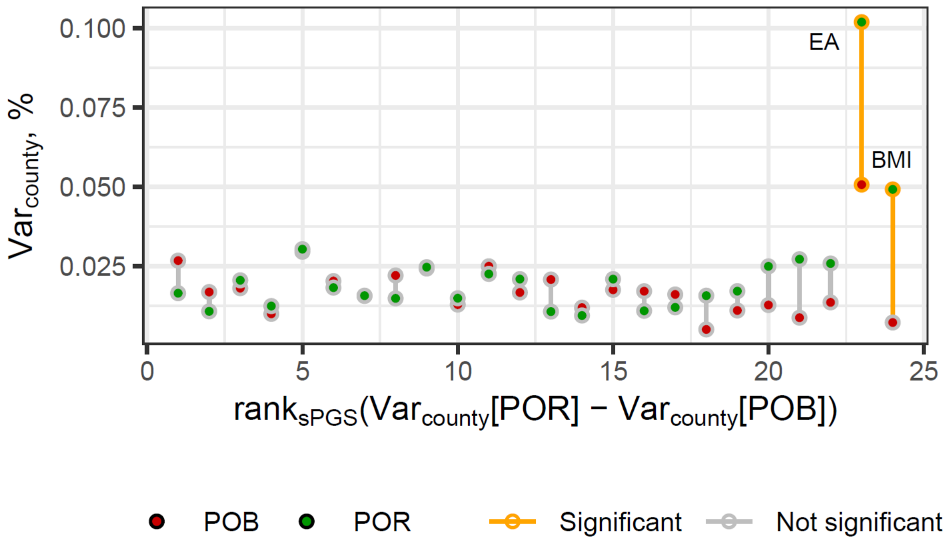

**Supplementary Figure 18. Estimates of the inter-individual variance of sPGSs among unrelated Estonian participants explained by POB and POR.** sPGSs are adjusted for demographic and genetic ancestry covariates. PGSs and sPGSs are ordered according to the rank of difference between $Var_{county}$ for POR and POB in the full Estonian subsample. Estimates significantly different from zero are outlined in yellow. The line connecting the two points is yellow when the variance explained by POB and POR together is significantly larger than the variance explained by only the weaker predictor. The significance level is 0.05, adjusted for the number of sPGS tested with Bonferroni correction.

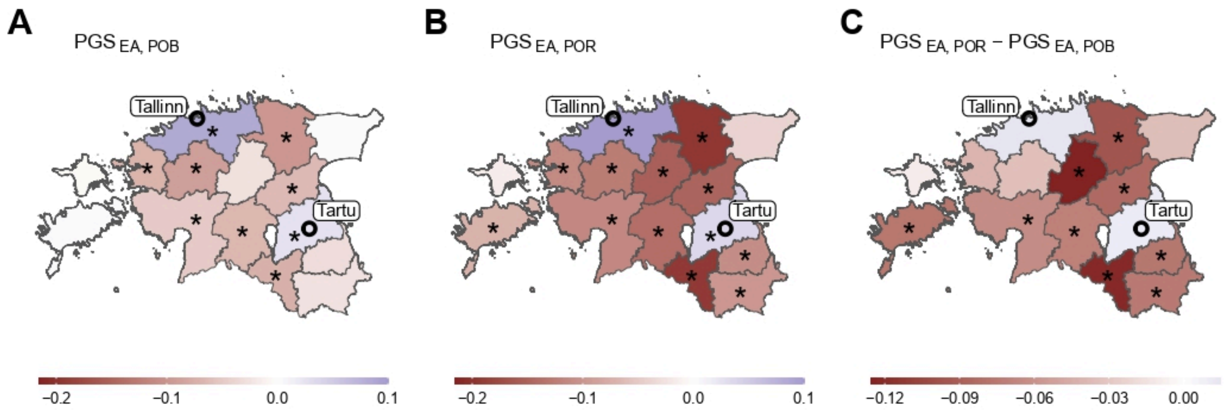

**Supplementary Figure 19. $PGS_{EA}$ landscape in Estonia among unrelated Estonian participants.** Mean $PGS_{EA}$ of individuals (A) born or (B) residing in each county. (C) Differences between values in "B" and "A" panels. $PGS_{EA}$ is adjusted for demographic and genetic ancestry covariates. Counties with sample mean values significantly different from zero after FDR correction at the 0.05 level are marked with an asterisk (*).

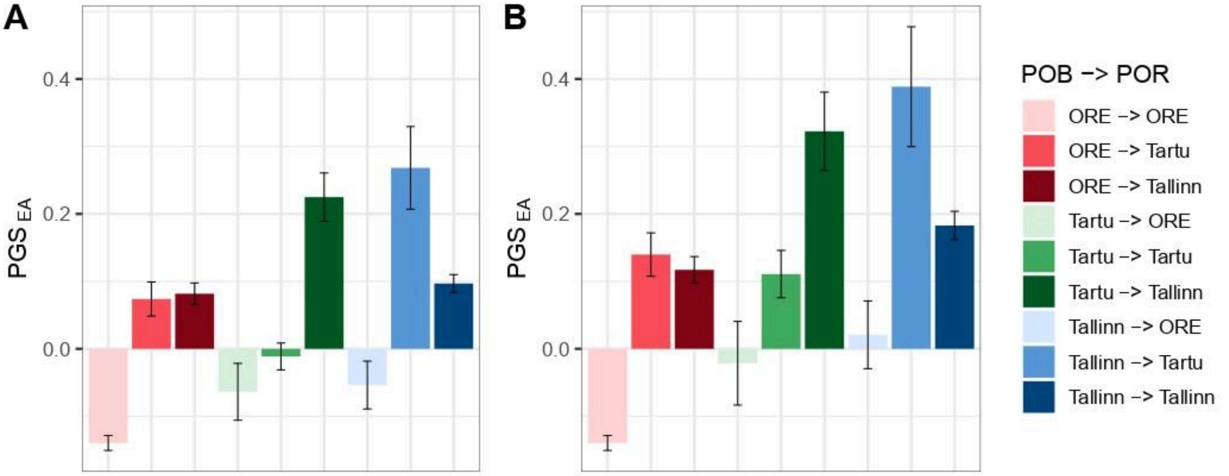

**Supplementary Figure 20. $PGS_{EA}$ in migration groups among unrelated Estonian participants by region of birth (POB) and residence (POR).** (A) County-based analysis where POB and POR refer to Tartu County ("Tartu"), Harju County ("Tallinn") and other counties ("ORE"). (B) City-based analysis, where POB and POR refer to Tartu City ("Tartu"), Tallinn ("Tallinn") and other counties ("ORE"). $PGS_{EA}$ is adjusted for demographic and genetic ancestry covariates. Error bars correspond to 95% confidence intervals.

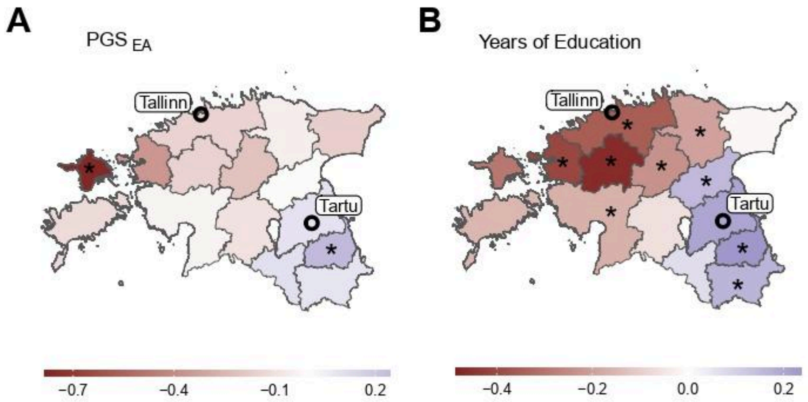

**Supplementary Figure 21. The contrast in mean $PGS_{EA}$ and EA (years of education) between residents of Tallinn and Tartu City among unrelated Estonian participants by county of birth.** (A) The value for each county corresponds to the mean $PGS_{EA}$ of individuals born in that county and living in Tartu City subtracted from the mean $PGS_{EA}$ of individuals born in the same county and living in Tallinn. Individuals born in Tallinn or Tartu City are excluded from the analysis. (B) The same but for the "years of education" phenotype. Counties with significant differences between the migrant groups after FDR correction at level 0.05 are marked with an asterisk (*).

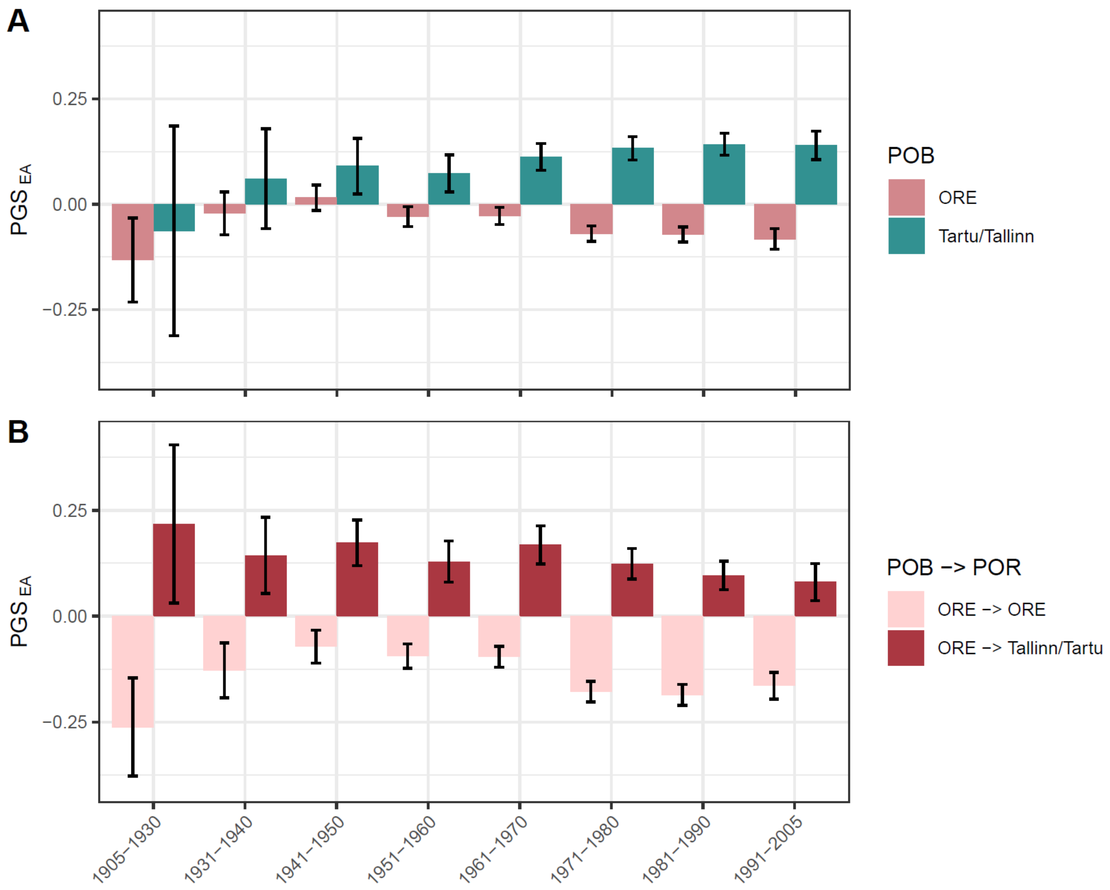

**Supplementary Figure 22. Difference in average $PGS_{EA}$ between cities (Tallinn and Tartu combined) and ORE across birth year bins in the sample of unrelated Estonians.** (A) Mean $PGS_{EA}$ of individuals born in either ORE or Tallinn/Tartu; (B) mean $PGS_{EA}$ of individuals born in ORE and residing in either ORE or Tallinn/Tartu. $PGS_{EA}$ is adjusted for the top 100 PCs and demographic covariates. Error bars correspond to 95% confidence intervals.

#### Supplementary Note 6. Replication of the main results with $PGS_{EA4}$

The analysis of $Var_{county}$ showed that SES-related PGSs demonstrate the highest increase in inter-regional variance with $PGS_{EA}$ being the top signal. Most of the inter-regional variance of other PGSs can be explained through their correlation with $PGS_{EA}$. Thus, in the main text, we provide results of the population-level analyses with $PGS_{EA}$.

However, if the $Var_{county}$ patterns for the tested PGSs are mostly linked to the EA-associated loci, a more powerful PGS for EA is expected to demonstrate an even stronger increase in $Var_{county}$ due to migration. Also, correcting for a more powerful PGS for EA is expected to reduce the signal for other PGSs even further. To test this, we used the summary statistics from the EA4 GWAS[13] which were the most powerful summary statistics for EA available to us. Indeed, $PGS_{EA4}$ demonstrates the highest $Var_{county}$ for both POB and POR with a particular increase in $Var_{county}$ for POR in comparison with that of $PGS_{EA}$ (Supplementary Figure 23A). In comparison with the adjustment for $PGS_{EA}$, the adjustment for $PGS_{EA4}$ stronger decreases $Var_{county}$ for other PGSs following the expectation (Supplementary Figure 23B, Main Text).

As the EA4 GWAS is a meta-analysis of a large number of cohorts with various sample sizes, it is more prone to confounding due to residual population stratification. $PGS_{EA}$ is based on the UK Biobank cohort GWAS and thus is less vulnerable to this effect[18,19]. Sibship-based analyses are robust towards residual confounding in PGSs thus we use $PGS_{EA4}$ there for more statistical power. Other analyses that are described for $PGS_{EA}$ are conducted also for $PGS_{EA4}$. We present their results in this supplementary note.

Briefly, all the original results are reproduced with $PGS_{EA4}$. Moreover, absolute differences between groups by POB, POR, or migration profile for $PGS_{EA4}$ are in general larger than for $PGS_{EA}$ (Supplementary Figures 24 and 27). These differences also reach the threshold of significance more often (Supplementary Figures 24 and 26).

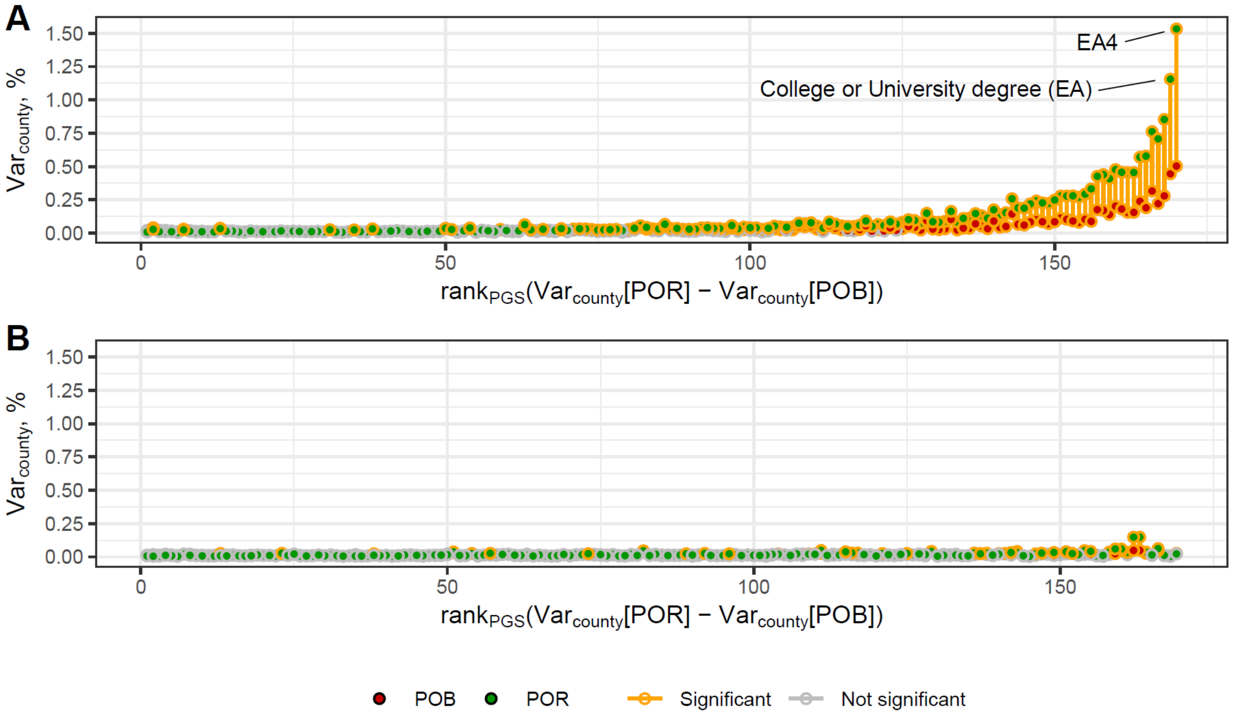

**Supplementary Figure 23. Fraction of the inter-individual variance of (A) PGSs and (B) PGSs additionally adjusted for $PGS_{EA4}$, explained by county of birth (POB) and county of residence (POR)**. PGSs are preliminary adjusted for the top 100 PCs and demographic covariates. In panel A the PGSs on the x-axis are ordered according to the difference between POR and POB which is the same order as in Figure 1B with the addition of $PGS_{EA4}$. In panel B the order is the same as in panel A but without $PGS_{EA4}$. $Var_{county}$ for PGSs adjusted for the top 100 PCs and demographic covariates. Red and green dots refer to the POB and POR, correspondingly. Estimates significantly different from zero are outlined in yellow. The line connecting the two points is yellow when the variance explained jointly by POB and POR is significantly larger than the variance explained by only the weaker predictor (if significant) or when the stronger predictor is significant. The significance level is 0.05, after Bonferroni correction.

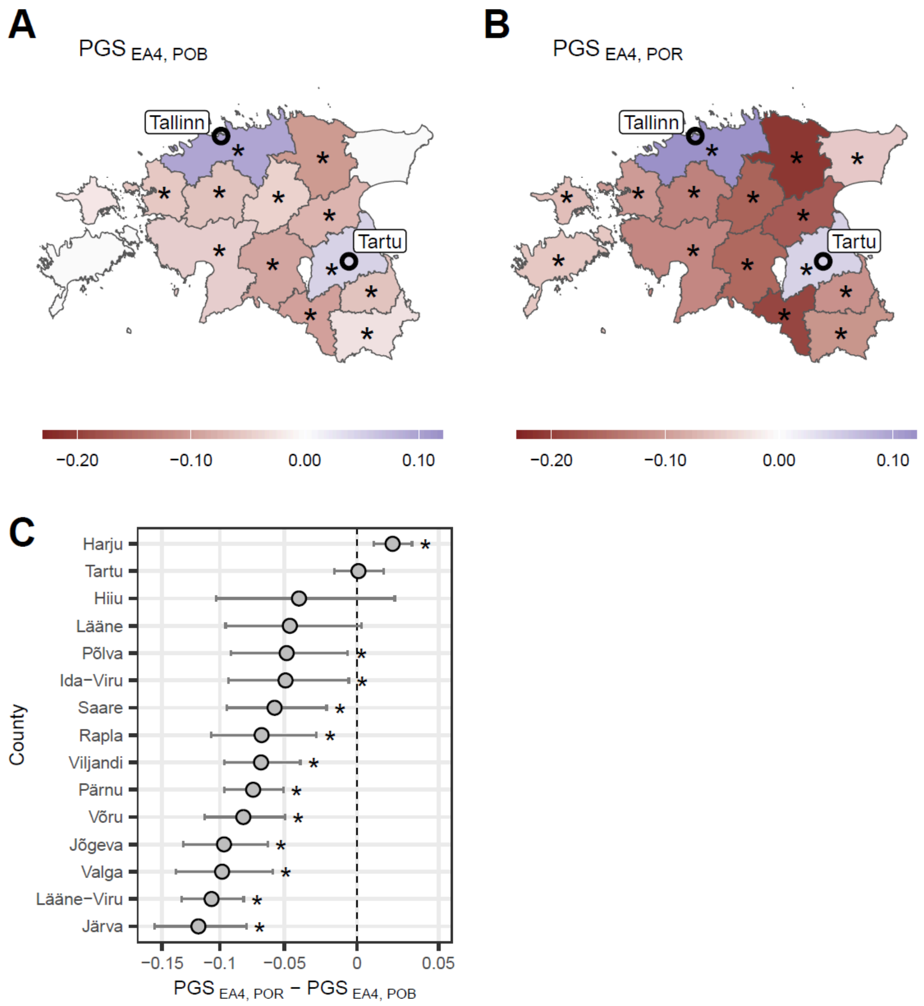

**Supplementary Figure 24. $PGS_{EA4}$ landscape in Estonia.** Mean $PGS_{EA4}$ of individuals (A) born or (B) residing in each county. (C) Difference between values in panels B and A. $PGS_{EA4}$ is adjusted for the top 100 PCs and demographic covariates. Counties with the corresponding value being significantly different from zero after FDR correction at the 0.05 significance level are marked with an asterisk (*). $PGS_{EA4}$ is measured in standard deviations.

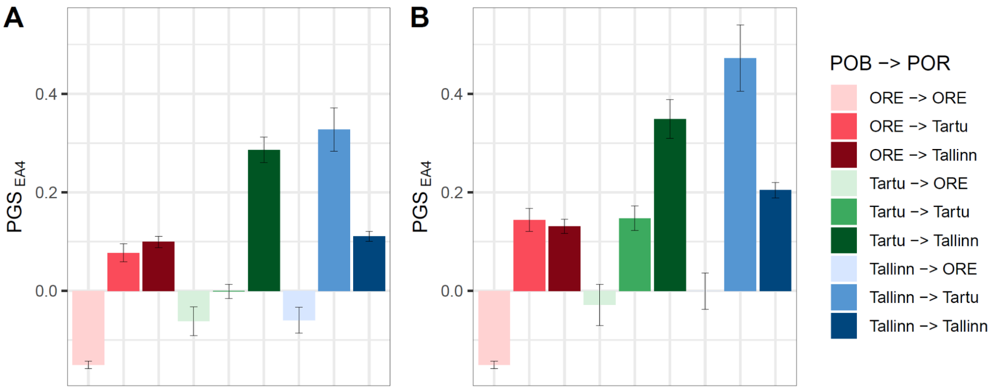

**Supplementary Figure 25. $PGS_{EA4}$ in migration groups defined by combination of place of birth (POB) and residence (POR).** (A) County-based analysis where “Tartu” and “Tallinn” refer to Tartu County and Harju County respectively while “ORE” refers to other counties. (B) City-based analysis, where “Tartu” and “Tallinn” refer to the respective cities while “ORE” refers to other counties as in A. $PGS_{EA}$ is adjusted for the top 100 PCs and demographic covariates. In all panels error bars correspond to 95% confidence intervals.

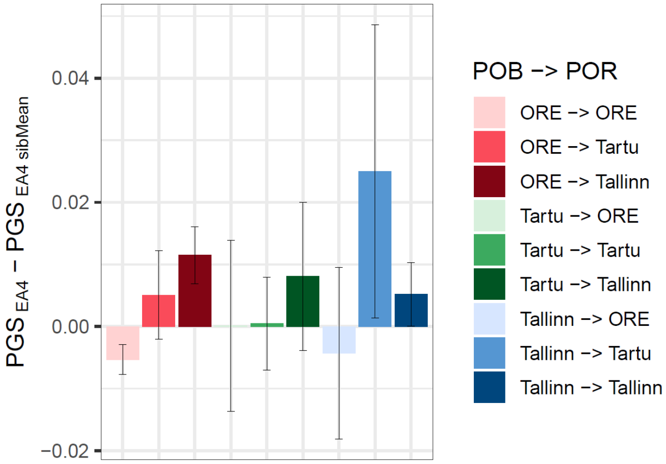

**Supplementary Figure 26. $PGS_{EA4}$ adjusted for sibship-average in migration groups defined by combination of place of birth (POB) and residence (POR).** “Tartu” and “Tallinn” refer to Tartu County and Harju County respectively while “ORE” refers to other counties. Error bars correspond to 95% confidence intervals.

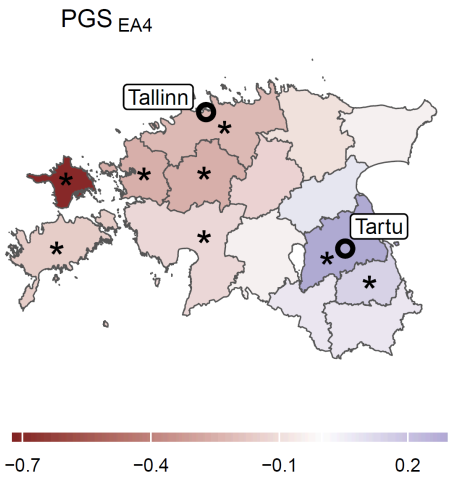

**Supplementary Figure 27. The difference in mean PGS$_{EA4}$ between residents of Tallinn and Tartu City by county of birth.** The value for each county corresponds to the mean PGS$_{EA4}$ of individuals born in that county and residing in Tartu City subtracted from the mean PGS$_{EA}$ of individuals born in the same county and residing in Tallinn. Individuals born in Tallinn or Tartu City are excluded from the analysis. PGS$_{EA}$ is adjusted for the top 100 PCs and demographic covariates. Counties with the difference being significant after FDR correction at level 0.05 are marked with an asterisk (*).

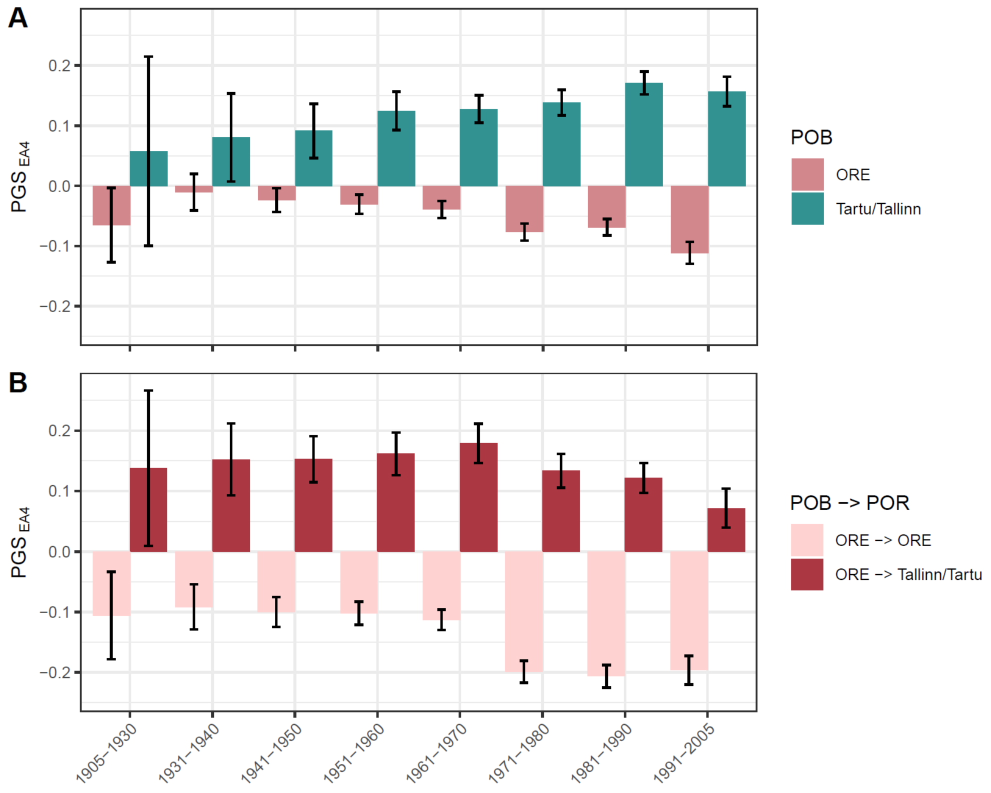

**Supplementary Figure 28. Difference in average $PGS_{EA4}$ between cities (Tallinn and Tartu combined) and ORE across birth year bins.** (A) Mean $PGS_{EA}$ of individuals born in either ORE or Tallinn/Tartu; (B) mean $PGS_{EA4}$ of individuals born in ORE and residing in either ORE or Tallinn/Tartu. $PGS_{EA4}$ is adjusted for the top 100 PCs and demographic covariates. Error bars correspond to 95% confidence intervals.

#### Supplementary Note 7. Selective migration and correlations between mate-pair PGSs

Assortative mating (AM) refers to genetic similarity between partners resulting from mate choice based on phenotype. It has been shown that for educational attainment (EA), the correlation between genetic predictors in partners is higher than expected based on phenotypic similarity[13,20]. Potential explanations for this include shared genetic ancestry between partners or assortment based on traits genetically correlated with EA.

Another factor contributing to genetic similarity between partners may be geographic clustering due to selective migration, as documented in this study and by Abdellaoui et al.[21]. Selective migration can lead to individuals with lower or higher PGSs for certain traits (particularly EA) being more prevalent in specific regions. Since individuals typically find partners relatively close to their place of residence, we expect a correlation between partners' PGSs even in the absence of AM, driven by mating by proximity. However, the consequences of this process should be the same as those of AM: genetic similarity between partners at loci associated with specific traits.

Since the strongest observed differentiation between geographic regions in Estonia occurs in the PGSs for EA ($PGS_{EA}$ and $PGS_{EA4}$), we test our hypothesis using these PGS. Height and BMI are also well-known traits for which AM is observed in humans[20]. However, the observed differentiation due to contemporary migration is weaker for their corresponding PGSs ($PGS_{height}$ and $PGS_{BMI}$). Therefore, we do not expect a strong effect of mating by proximity on partner similarity for these PGSs. For each of the four tested PGSs, we calculated correlations between randomly selected pairs of individuals from: (1) the overall Estonian population, (2) individuals born in the same county (based on place of birth, POB), and (3) individuals residing in the same county (based on place of residence, POR). We then compared these correlations with those of actual spouses, defined as pairs of individuals sharing a child in the EstBB (Supplementary Figure 29). In both randomly selected pairs and spouses, the sex of individuals was not taken into account.

Correlations between pairs matched by POB as well as completely random pairs are not significantly different from zero for all the PGSs. However, the mean correlations for $PGS_{EA}$ and $PGS_{EA4}$ in pairs matched by POB are shifted towards positive values which is not observed for $PGS_{height}$ and $PGS_{BMI}$. While correlations for $PGS_{height}$ and $PGS_{BMI}$ in pairs matched by POR remain statistically indistinguishable from zero, correlations for $PGS_{EA}$ and $PGS_{EA4}$ are significantly higher than zero. The absence of correlations among pairs matched by POB suggests that the correlations for pairs matched by POR are not the consequence of residual population structure. At the same time, the slight positive shift in average estimates for $PGS_{EA}$ and $PGS_{EA4}$ are likely caused by clustering due to migration in previous generations.

These results suggest that mating by proximity is a likely factor inflating estimates of the strength of AM for EA and potentially for other traits with a high genetic correlation with EA. Here, we selected pairs of individuals based on county-level geographic information. However, mating by proximity likely has a stronger influence on partner similarity at a finer scale, for example, due to residential segregation in cities.

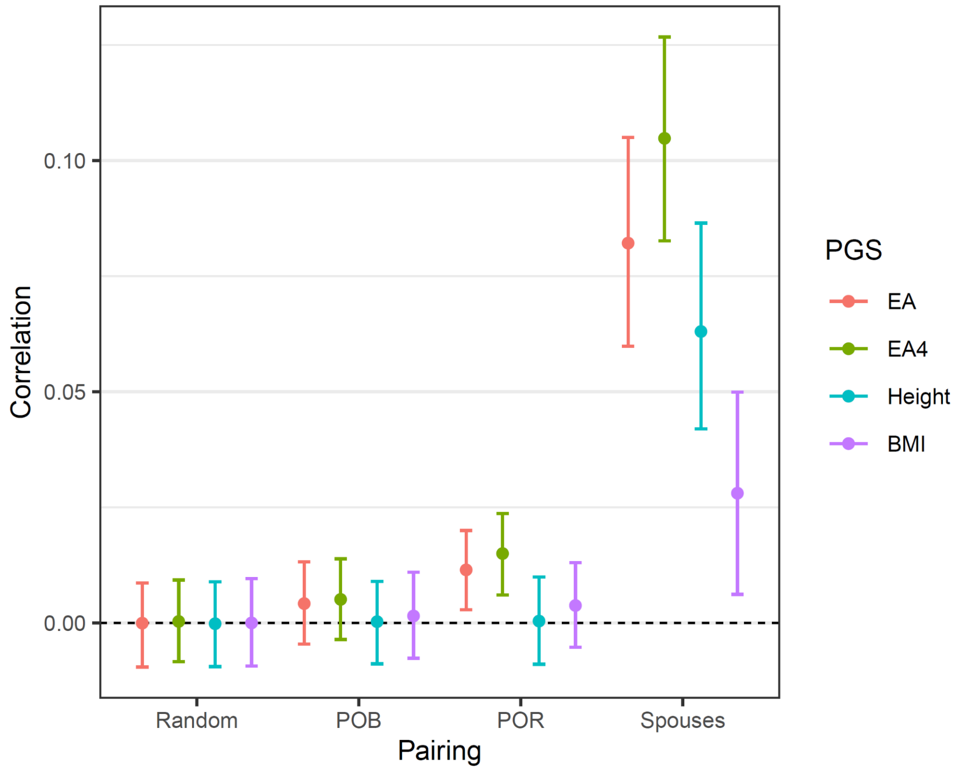

**Supplementary Figure 29. Mating by proximity and assortative mating in Estonia.** Correlation between (1) random pairs of individuals from the entire EstBB; (2) random paris of individulas with the same county-level place of birth (POB); (3) random paris of individulas with the same county-level place of residence (POR) and (4) spouses defined as pairs of individuals sharing a child in the EstBB. Error bars correspond to 95% confidence intervals (CIs). The estimates and CIs for the points 1-3 were derived from 1000 random sets of pairs picked according to the described rules. The CIs for spouses were calculated using bootstrap procedure with 1000 replicates.

### Supplementary Note 8. How large are the regional differences in $PGS_{EA}$?

We can see that the regional differences in polygenic scores are statistically significant and are increasing due to migrations. However, if we compare polygenic scores for education attainment among people born and living in Tallinn and Tartu City versus those born and living in other regions (ORE) we see only a subtle difference in their distributions (Supplementary Figure 30). Although those distributions are shifted, they overlap vastly. Practically, it means that a random individual born and residing in the city will have a polygenic score lower than that of a random individual born and residing outside the cities in close to 50% of the cases.

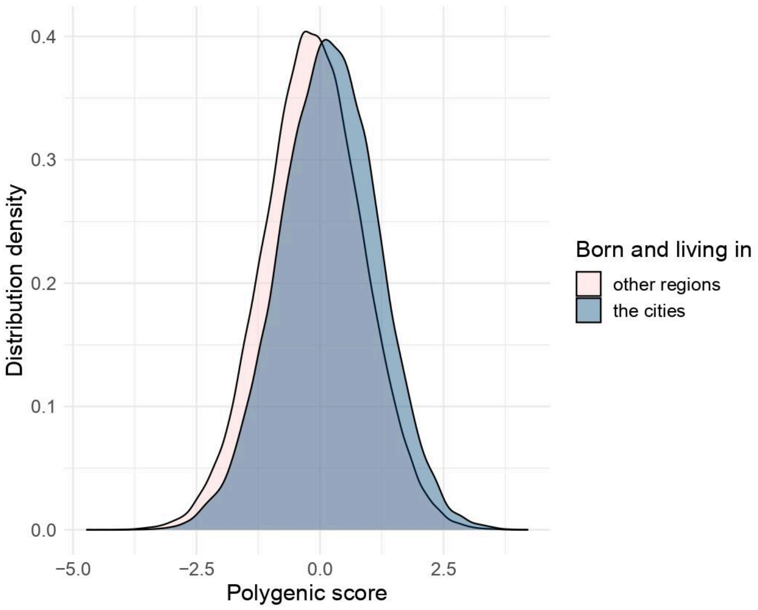

**Supplementary Figure 30. Comparison of the distribution of the polygenic score for educational attainment for individuals born and living in Tallinn and Tartu City versus those born and living in other regions of Estonia (ORE).**

We would also like to stress that differences in $PGS_{EA}$ between EstBB cohorts cannot be directly interpreted as regional differences in genetic predisposition to educational attainment in the general Estonian populations. First, although the Estonian Biobank includes approximately 20% of the country's adult population from a wide range of socioeconomic backgrounds and localities, the data set is not entirely representative of the Estonian population. Second, a polygenic score is a correlate of a trait and its genetic basis, not an exact genetic value. For any trait but especially for behavioural characteristics like educational attainment it accumulates a complex combination of direct and indirect effects as well as non-causal correlates. However, the polygenic scores based on the population-based genetic association study used in our analyses capture from 5 to 7% of individual differences in educational attainment (depending on underlying GWAS and analysed cohort). This is only a fraction of the total estimated genetic effect on educational attainment. Thus, $PGS_{EA}$ from a population-based GWAS should be seen as a relatively weak, noisy and confounded proxy for the genetic predisposition of an individual that affects their EA.

### Supplementary figures

#### Genetic predictors of ORE-to-cities migration

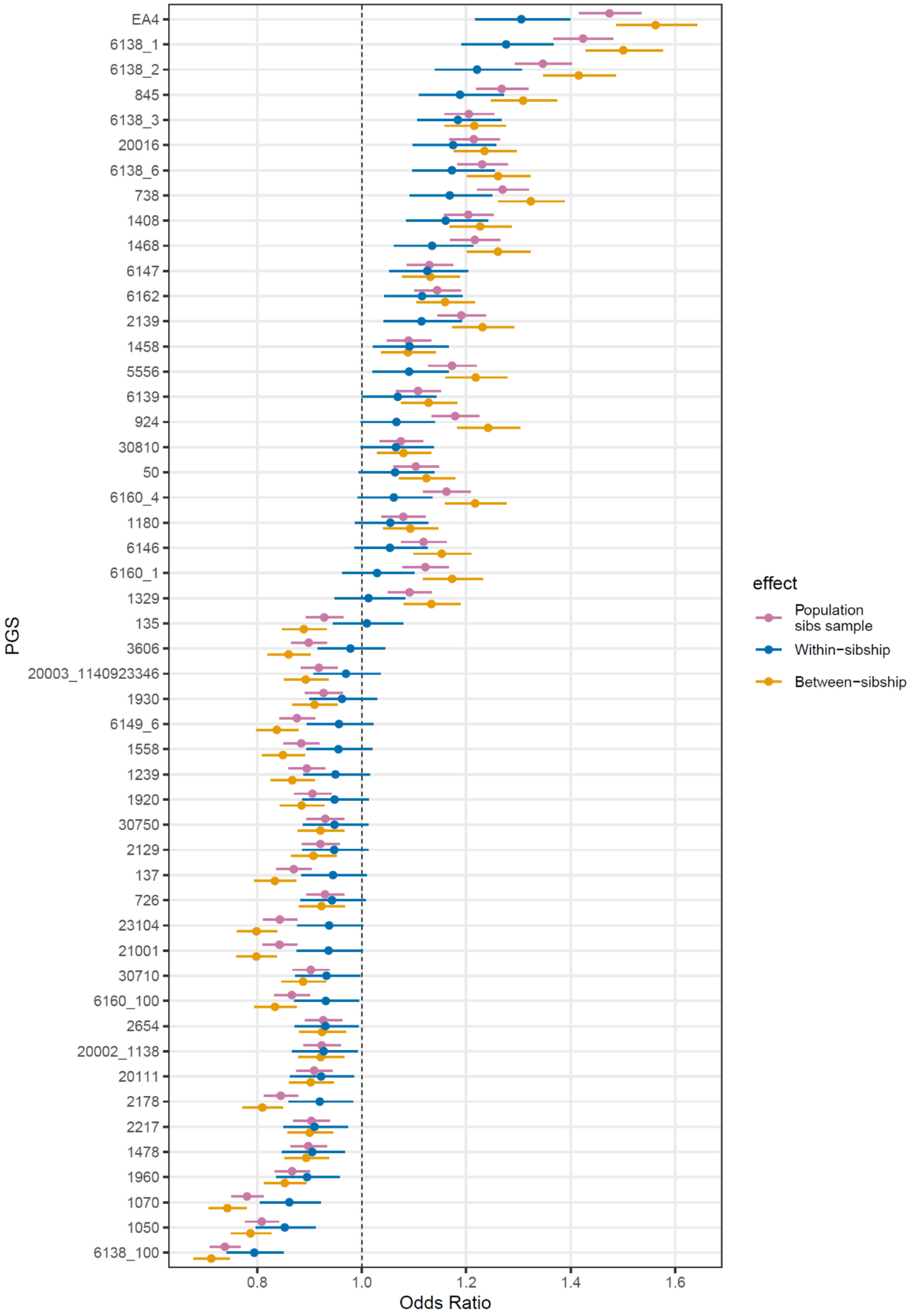

**Supplementary Figure 31. Mixed effects logistic regression results for all the PGSs tested as predictors of migration from ORE to the major cities (Tallinn or Tartu).** All the effects are estimated in a sample of siblings with only siblings born in the same county being included. The estimates are obtained using mixed effects logistic regression with a random intercept for sibship. All PGSs are preliminary adjusted for the top 100 PCs and the demographic covariates. Results are shown for PGSs with a significant population effect after the Bonferroni correction. Vertical dashed line indicates Odds Ratio equal to 1. Error bars correspond to 95% confidence intervals.

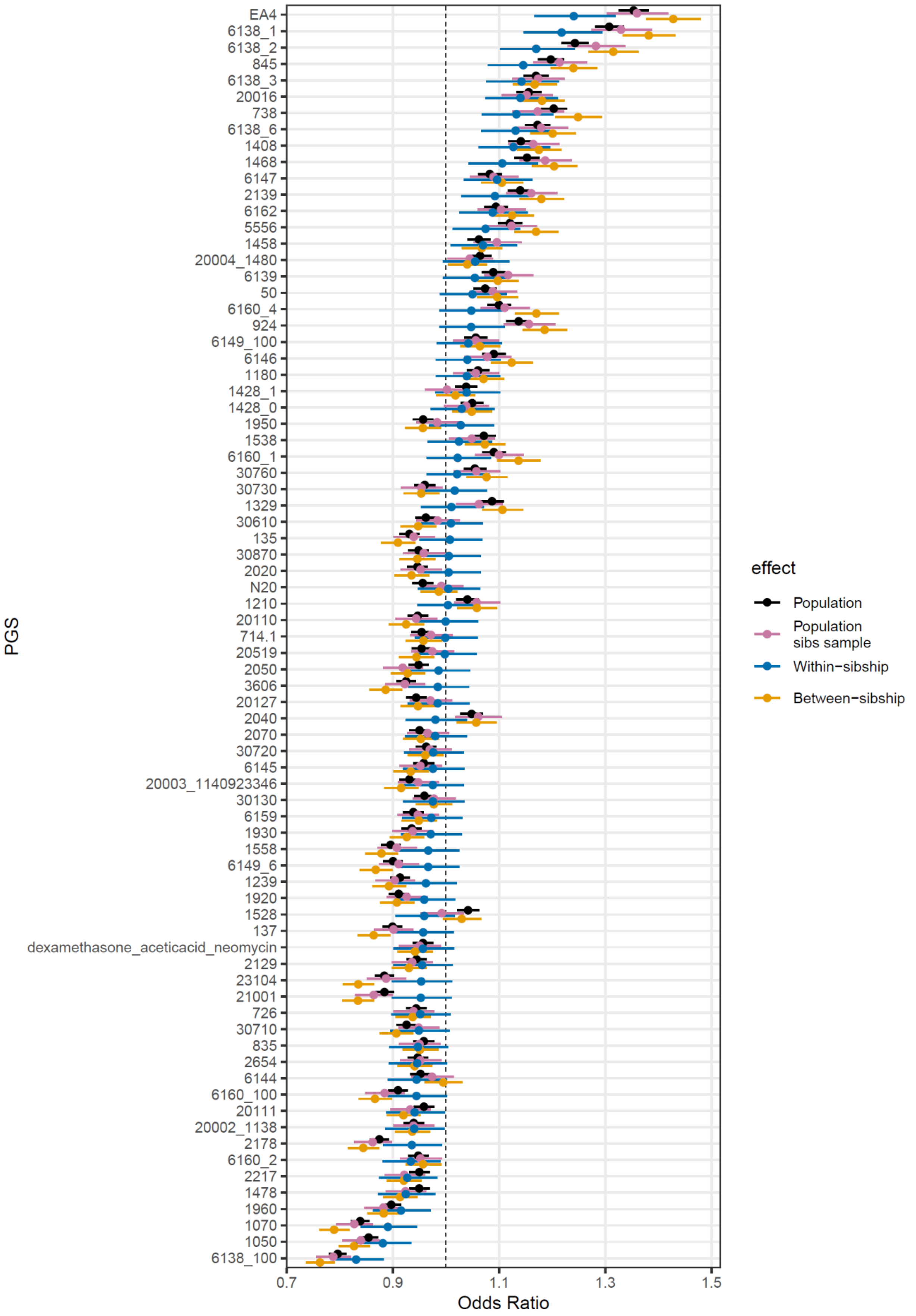
PGS
EA4
6138_1
6138_2
845
6138_3
20016
738
6138_6
1408
1468
6147
2139
6162
5556
1458
20004_1480
6139
50
6160_4
924
6149_100
6146
1180
1428_1
1428_0
1950
1538
6160_1
30760
30730
1329
30610
135
30870
2020
N20
1210
20110
714.1
20519
2050
3606
20127
2040
2070
30720
6145
20003_1140923346
30130
6159
1930
1558
6149_6
1239
1920
1528
137
dexamethasone_aceticacid_neomycin
2129
23104
21001
726
30710
835
2654
6144
6160_100
20111
20002_1138
2178
6160_2
2217
1478
1960
1070
1050
6138_100
0.7
0.9
1.1
1.3
1.5
Odds Ratio
effect
Population
Population sibs sample
Within−sibship
Between−sibship

**Supplementary Figure 32. A fixed effects logistic regression results for all the PGSs tested as predictors of migration from ORE to the major cities (Tallinn or Tartu).** Population effects are estimated in the subsample of unrelated Estonians. The other effects are estimated in a sample of siblings with only siblings born in the same county being included. The estimates are obtained using fixed effects logistic regression. All PGSs are preliminary adjusted for the top 100 PCs and the demographic covariates. Results are shown for PGSs with a significant population effect estimated in the subsample of unrelated Estonians after the Bonferroni correction. Vertical dashed line indicates Odds Ratio equal to 1. Error bars correspond to 95% confidence intervals.

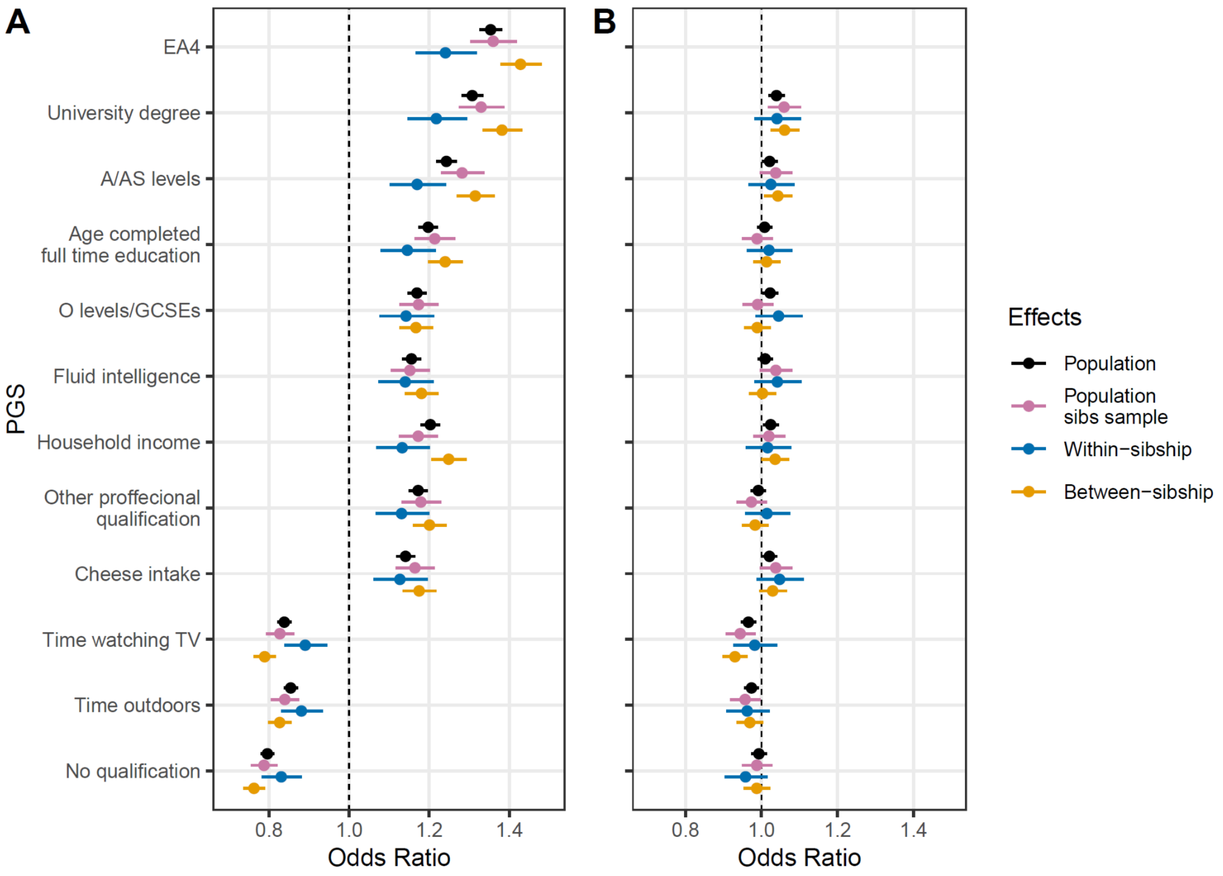

**Supplementary Figure 33. Fixed effects logistic regression results for PGSs with a significant within-sibship effect after the Bonferroni correction in Supplementary Figure 32, as predictors of migration from ORE to the major cities (Tallinn or Tartu).** Population effects are estimated in the subsample of unrelated Estonians. The other effects are estimated in a sample of siblings with only siblings born in the same county being included. (A) Effect sizes for PGSs; (B) Effect sizes of PGSs additionally adjusted for $PGS_{EA4}$. All PGSs are preliminary adjusted for top 100 PCs and the demographic covariates. Vertical dashed line indicates Odds Ratio equal to 1. Error bars correspond to 95% confidence intervals.

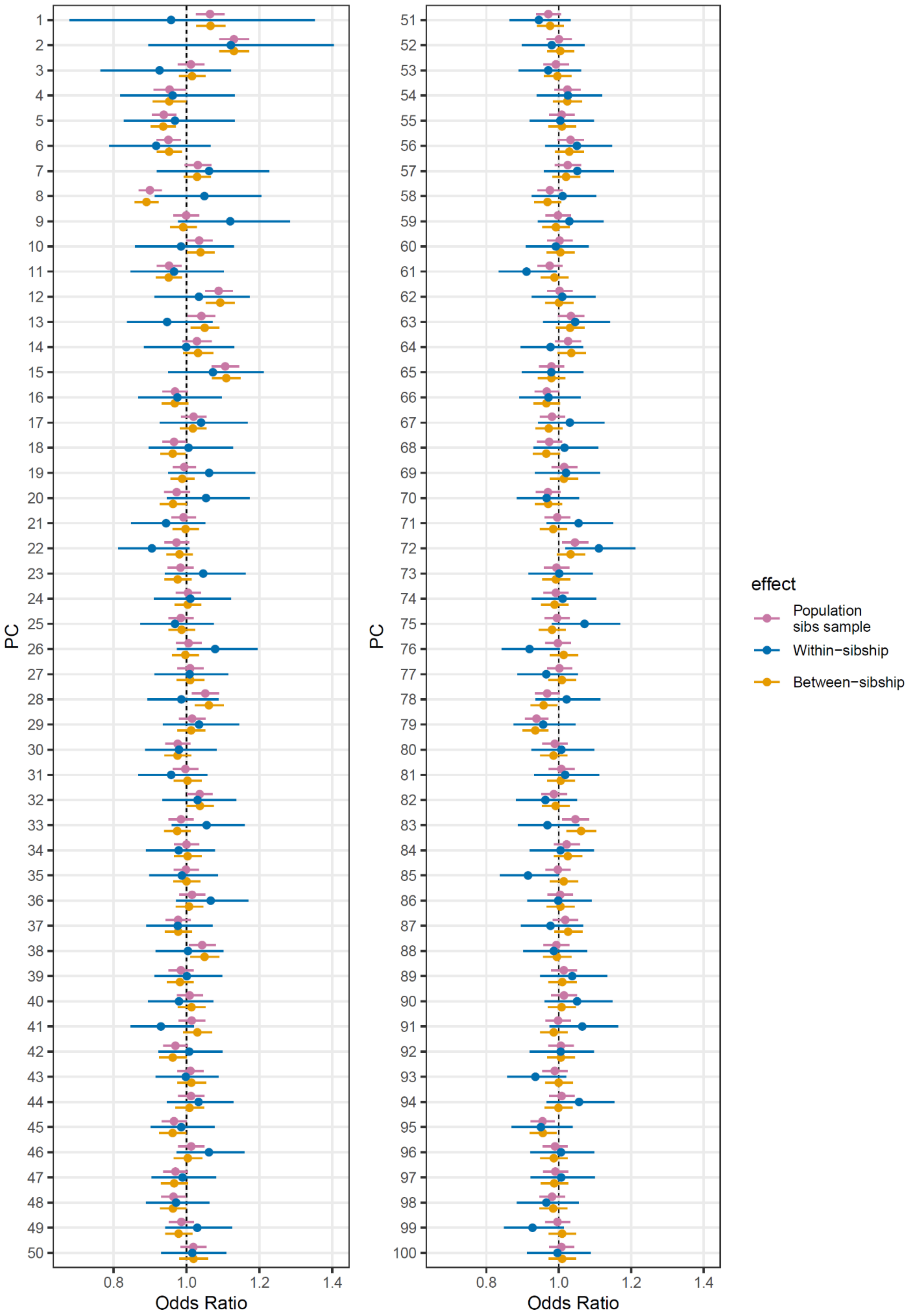
PC
Odds Ratio
0.8
1.0
1.2
1.4
effect
Population sibs sample
Within−sibship
Between−sibship

**Supplementary Figure 34. Mixed effects logistic regression results for the top 100 PCs as predictors of migration from ORE to the major cities (Tallinn or Tartu).** All the effects are estimated in a sample of siblings with only siblings born in the same county being included. The estimates are obtained using mixed effects logistic regression with a random intercept for sibship. Vertical dashed line indicates Odds Ratio equal to 1. Error bars correspond to 95% confidence intervals.

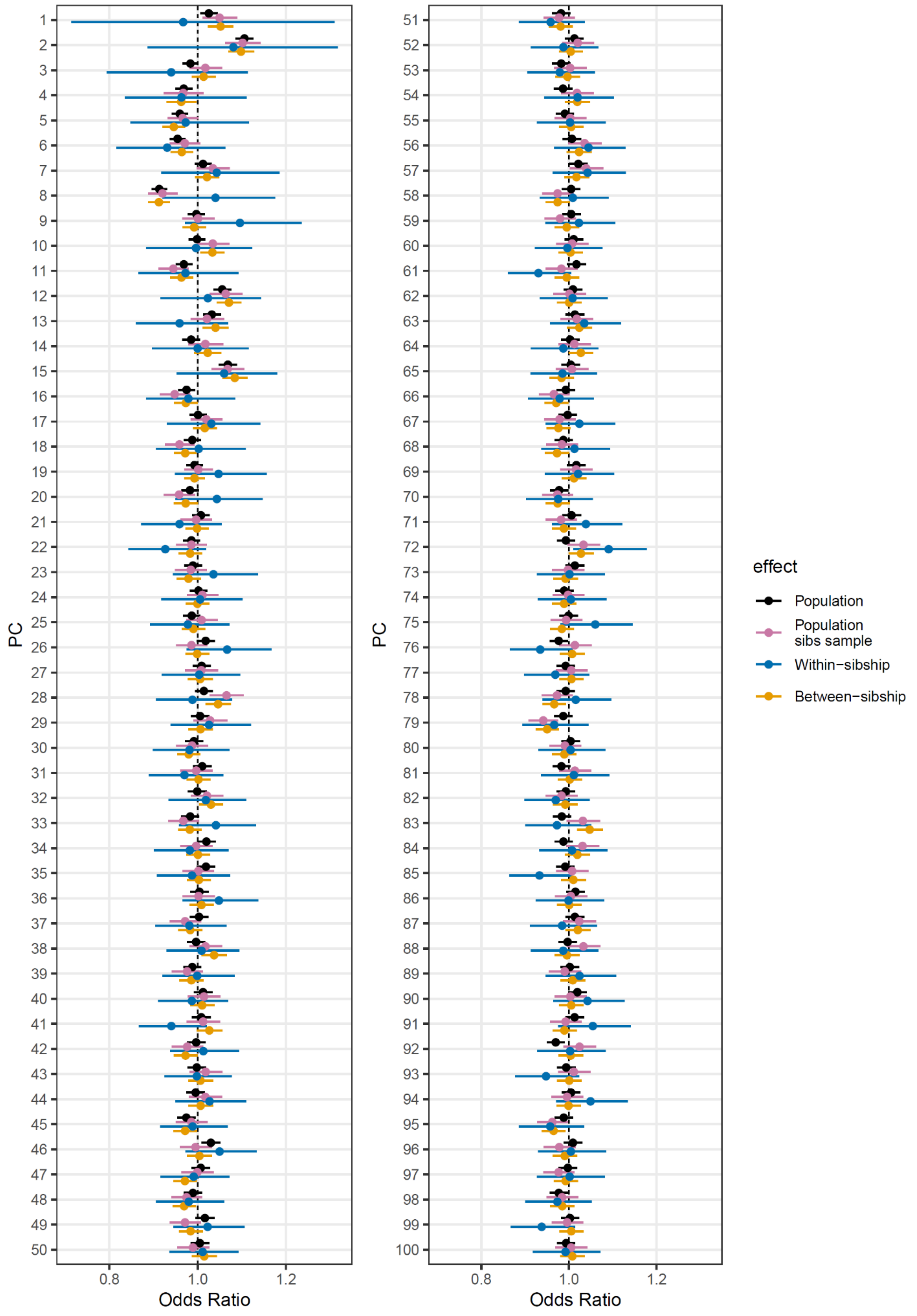

PC
Odds Ratio
0.8
1.0
1.2
effect
Population
Population sibs sample
Within−sibship
Between−sibship

**Supplementary Figure 35. Fixed effects logistic regression results for the top 100 PCs as predictors of migration from ORE to the major cities (Tallinn or Tartu).** Population effects are estimated in the subsample of unrelated Estonians. The other effects are estimated in a sample of siblings with only siblings born in the same county being included. The estimates are obtained using fixed effects logistic regression. Vertical dashed line indicates Odds Ratio equal to 1. Error bars correspond to 95% confidence intervals.

#### Geographical distribution of (s)PGS$_{EA}$

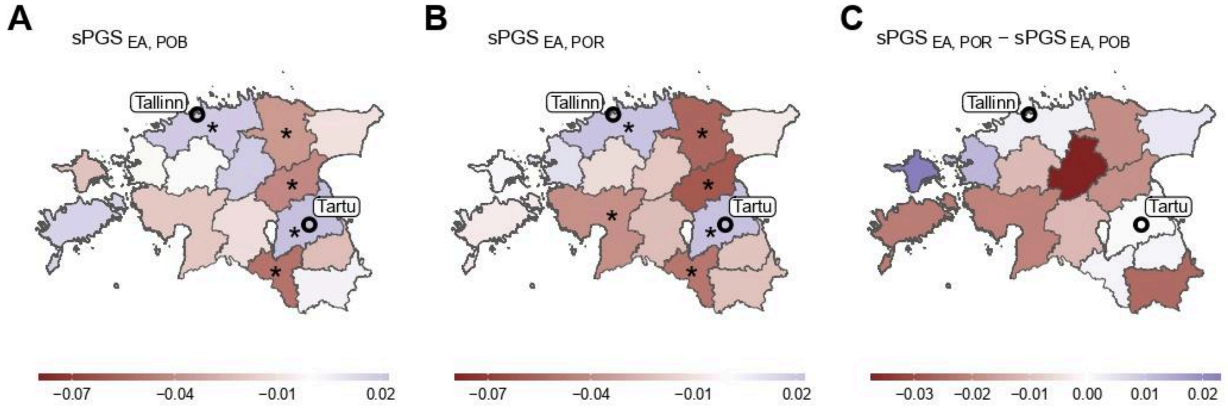

**Supplementary Figure 36. sPGS$_{EA}$ landscape in Estonia among Estonian participants.** Mean sPGS$_{EA}$ of individuals (A) born or (B) residing in each county. (C) Differences between values in "B" and "A" panels. PGS$_{EA}$ is adjusted for demographic and genetic ancestry covariates. Counties with sample mean values significantly different from zero after FDR correction at the 0.05 level are marked with an asterisk (*).

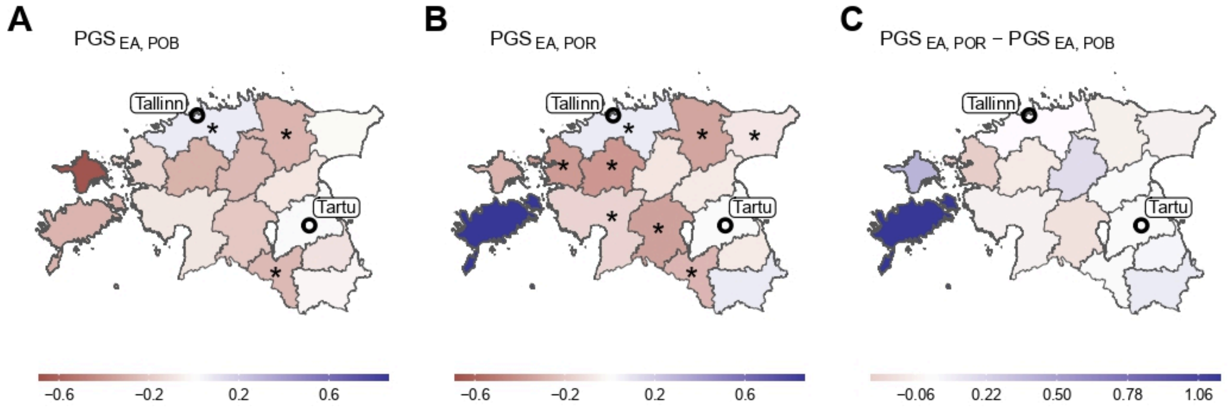

**Supplementary Figure 37. PGS$_{EA}$ landscape in Estonia among Russian participants.** Mean PGS$_{EA}$ of individuals (A) born or (B) residing in each county. (C) Differences between values in "B" and "A" panels. PGS$_{EA}$ is adjusted for demographic and genetic ancestry covariates. Counties with sample mean values significantly different from zero after FDR correction at the 0.05 level are marked with an asterisk (*).

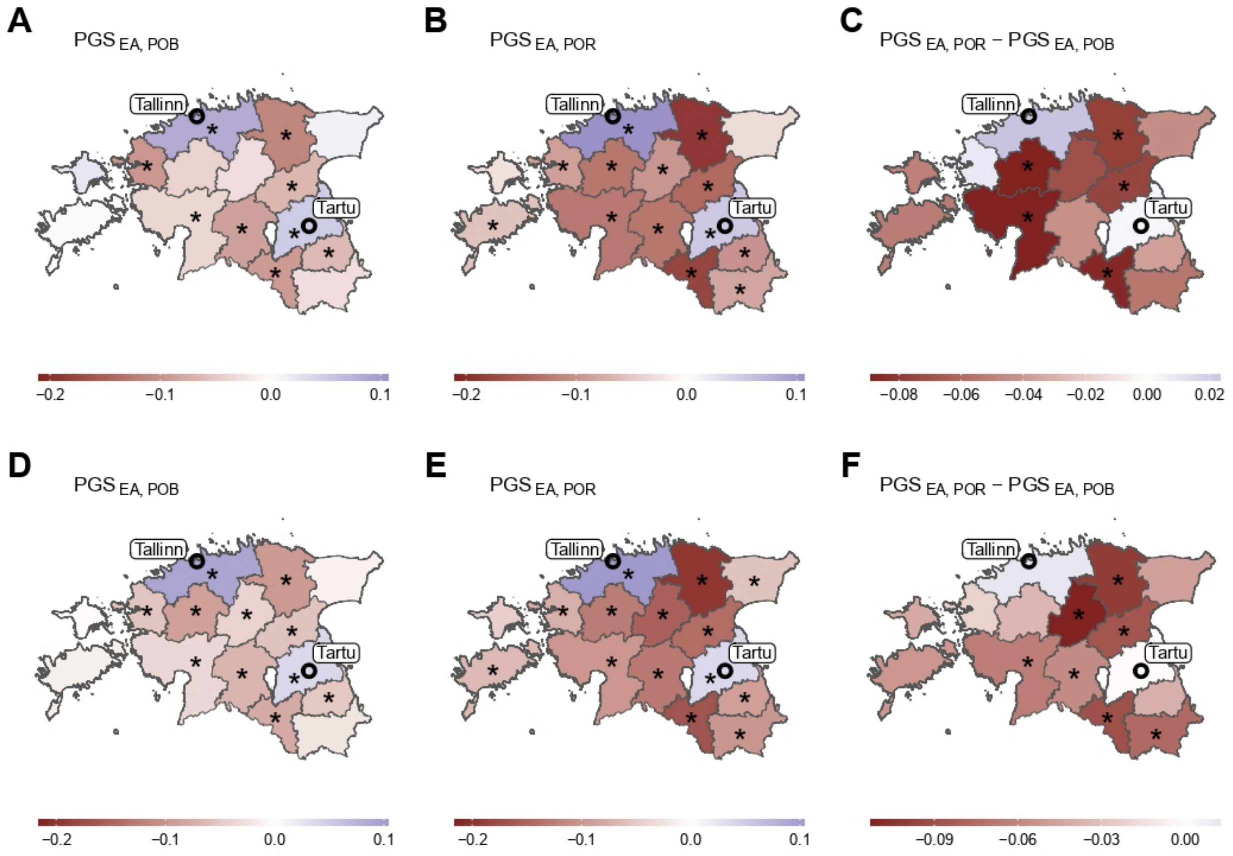

**Supplementary Figure 38. $PGS_{EA}$ landscape in Estonia among (A-C) male and (D-F) female Estonian participants.** Mean $PGS_{EA}$ of individuals (A, D) born or (B, E) residing in each county. (C, F) Differences between values in "B" and "A" panels ("E" and "D", correspondingly). $PGS_{EA}$ is adjusted for demographic and genetic ancestry covariates. Counties with sample mean values significantly different from zero after FDR correction at the 0.05 level are marked with an asterisk (*).

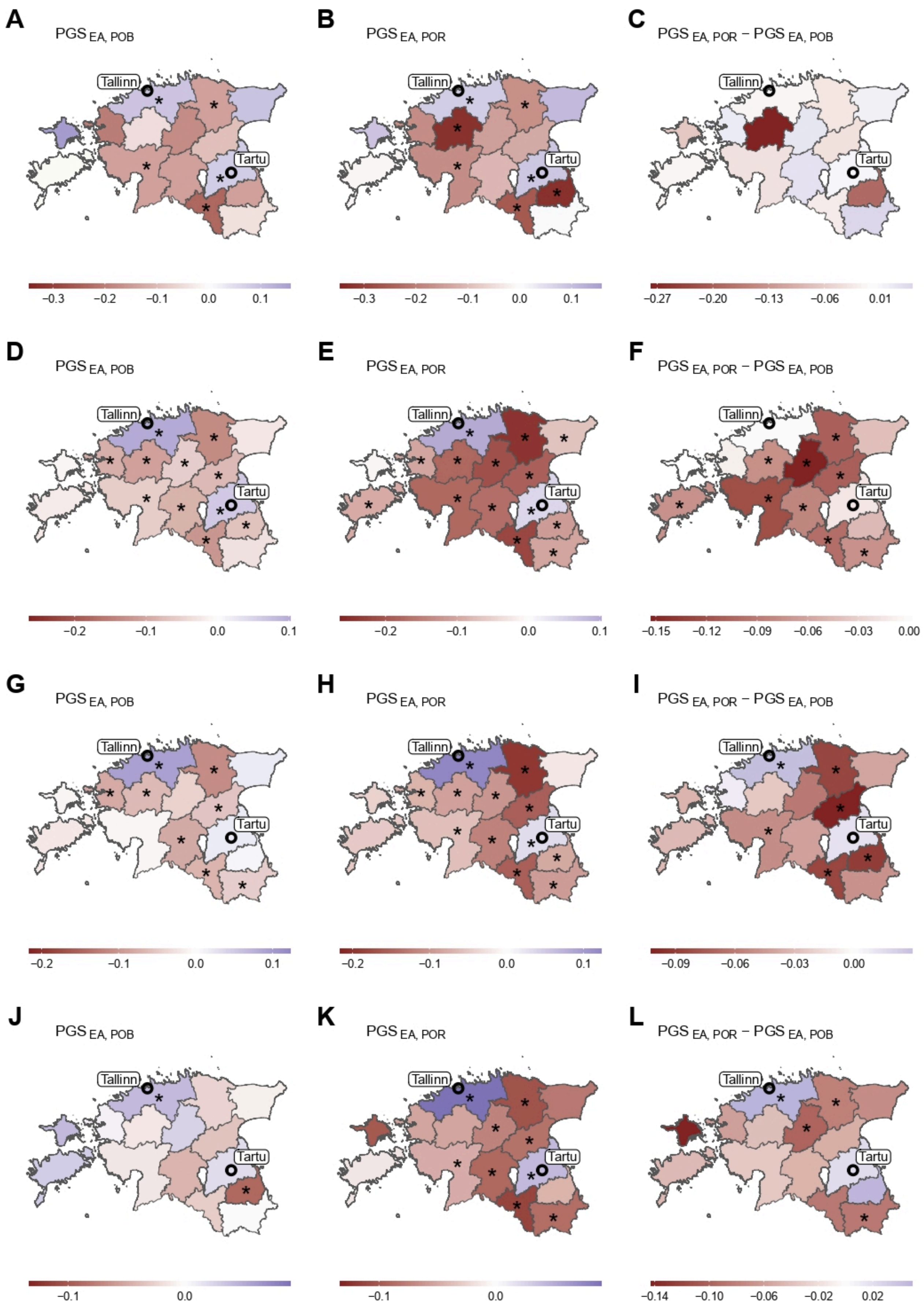
A
PGS EA, POB
B
PGS EA, POR
C
PGS EA, POR − PGS EA, POB
D
PGS EA, POB
E
PGS EA, POR
F
PGS EA, POR − PGS EA, POB
G
PGS EA, POB
H
PGS EA, POR
I
PGS EA, POR − PGS EA, POB
J
PGS EA, POB
K
PGS EA, POR
L
PGS EA, POR − PGS EA, POB
Tallinn
Tartu
−0.3 −0.2 −0.1 0.0 0.1
−0.27 −0.20 −0.13 −0.06 0.01
−0.2 −0.1 0.0 0.1
−0.15 −0.12 −0.09 −0.06 −0.03 0.00
−0.09 −0.06 −0.03 0.00
−0.1 0.0
−0.14 −0.10 −0.06 −0.02 0.02

**Supplementary Figure 39. $PGS_{EA}$ landscape in Estonia among Estonian participants stratified by age.** Age groups were defined as (A-C) 18-24, (D-F) 25-48, (G-I) 49-64, (J-L) 65+. Mean $PGS_{EA}$ of individuals (A, D, G, J) born or (B, E, H, K) residing in each county. (C, F, I, L) Differences between values in "B" and "A" panels ("E"-"D", "H"-"G", "K"-"J" correspondingly). $PGS_{EA}$ is adjusted for demographic and genetic ancestry covariates. Counties with sample mean values significantly different from zero after FDR correction at the 0.05 level are marked with an asterisk (*).

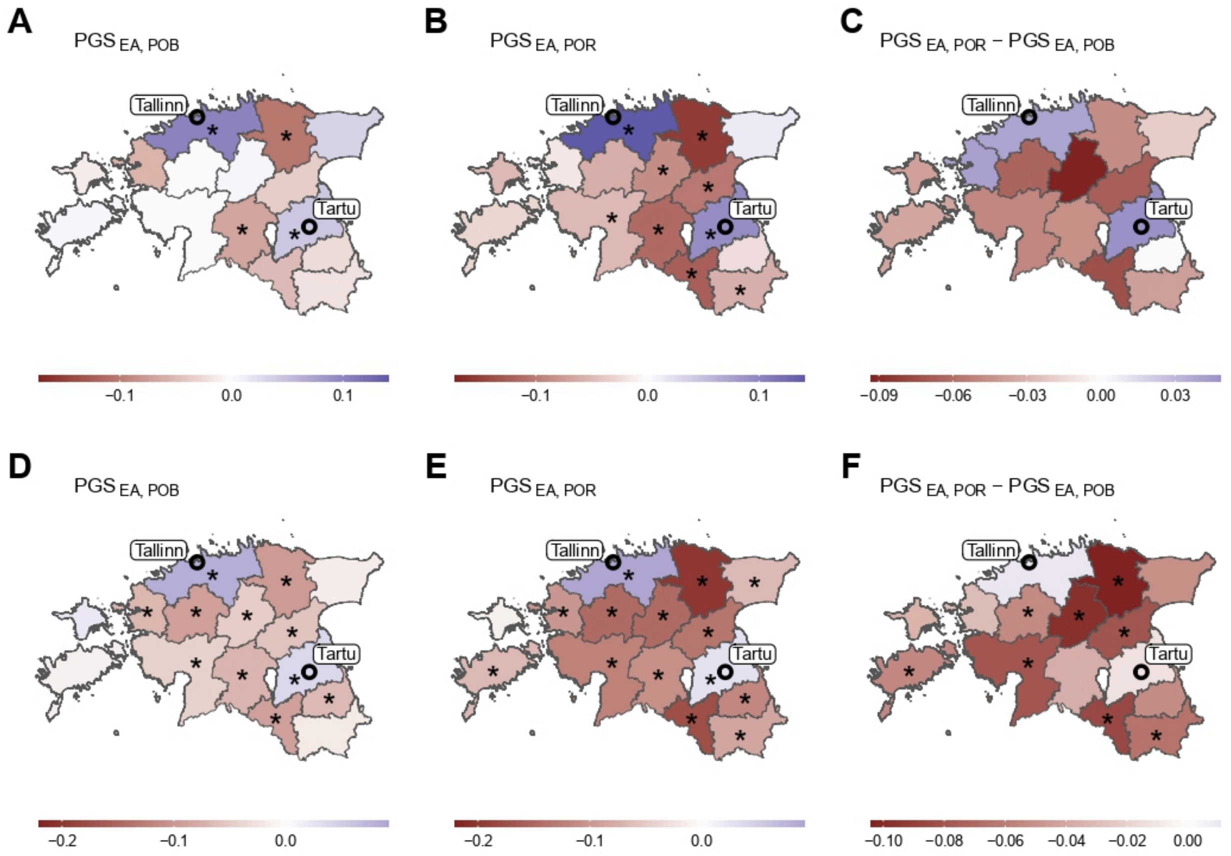

**Supplementary Figure 40. $PGS_{EA}$ landscape in Estonia among Estonian participants stratified by year of joining the biobank.** The periods of joining are (A) 2001-2016 and (B) 2017-2021. Mean $PGS_{EA}$ of individuals (A, D) born or (B, E) residing in each county. (C, F) Differences between values in "B" and "A" panels ("E" and "D", correspondingly). $PGS_{EA}$ is adjusted for demographic and genetic ancestry covariates. Counties with sample mean values significantly different from zero after FDR correction at the 0.05 level are marked with an asterisk (*).

#### Geographical distribution of educational attainment phenotype

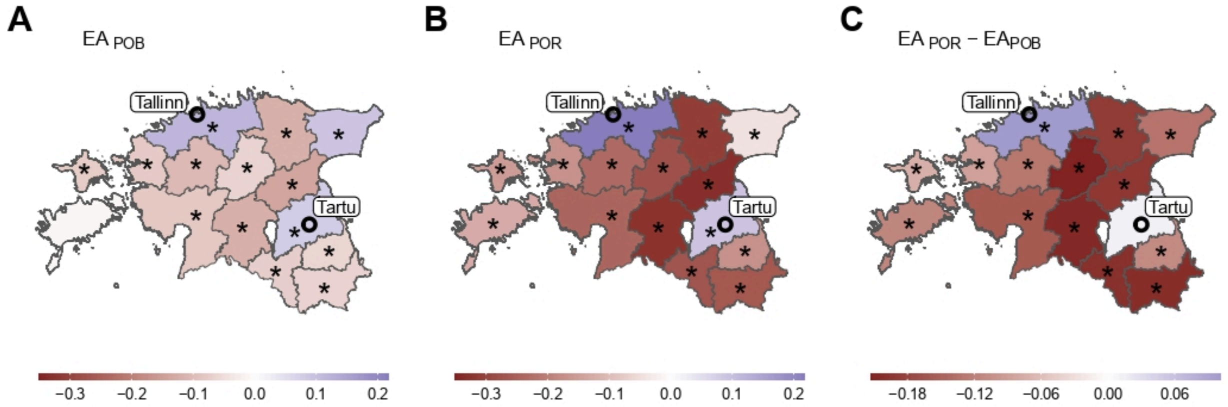

**Supplementary Figure 41. EA (years of education) landscape in Estonia among Estonian participants.** Mean EA of individuals (A) born or (B) residing in each county. (C) Differences between values in "B" and "A" panels. EA is adjusted for demographic and genetic ancestry covariates. Counties with sample mean values significantly different from zero after FDR correction at the 0.05 level are marked with an asterisk (*).

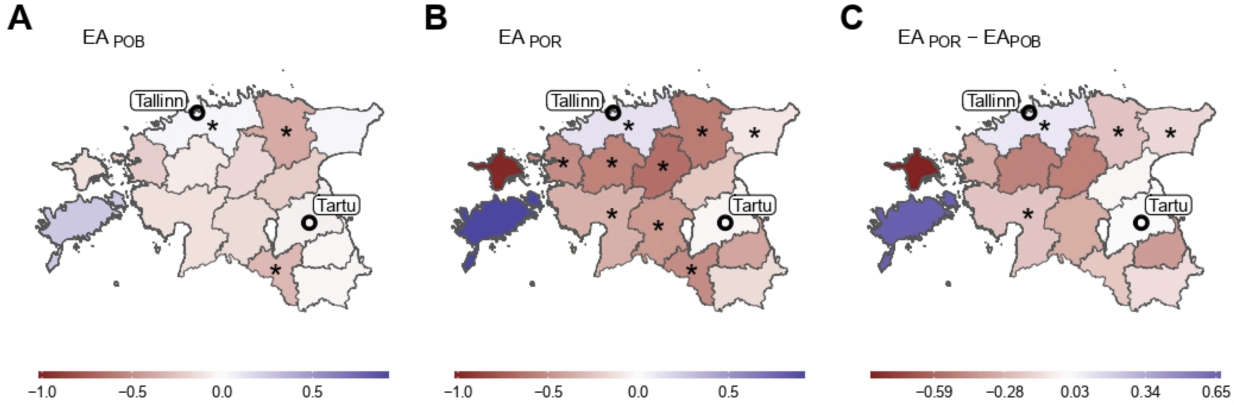

**Supplementary Figure 42. EA (years of education) landscape in Estonia among Russian participants.** Mean EA of individuals (A) born or (B) residing in each county. (C) Differences between values in "B" and "A" panels. EA is adjusted for demographic and genetic ancestry covariates. Counties with sample mean values significantly different from zero after FDR correction at the 0.05 level are marked with an asterisk (*).

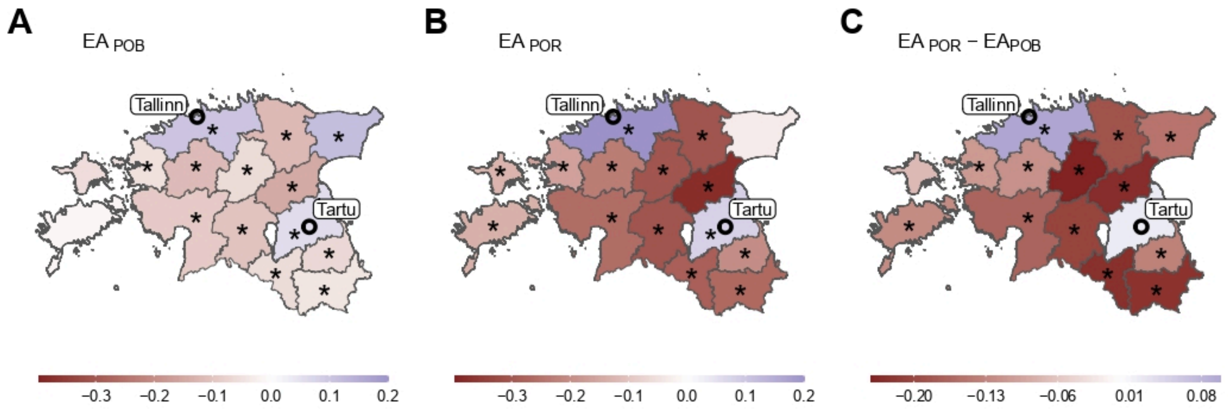

**Supplementary Figure 43. EA (years of education) landscape in Estonia among unrelated Estonian participants.** Mean EA of individuals (A) born or (B) residing in each county. (C) Differences between values in "B" and "A" panels. EA is adjusted for demographic and genetic ancestry covariates. Counties with sample mean values significantly different from zero after FDR correction at the 0.05 level are marked with an asterisk (*).

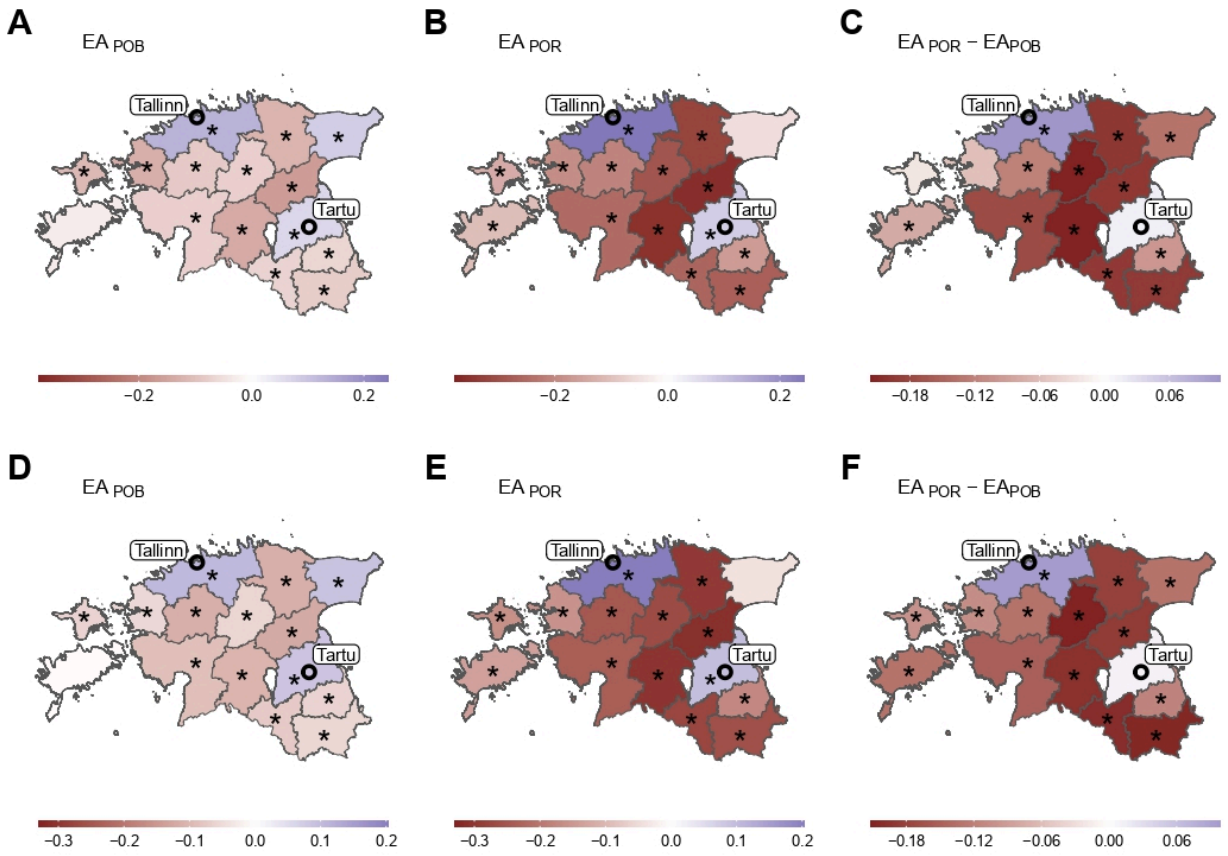

**Supplementary Figure 44. EA (years of education) landscape in Estonia among (A-C) male and (D-F) female Estonian participants.** Mean EA of individuals (A, D) born or (B, E) residing in each county. (C, F) Differences between values in "B" and "A" panels ("E" and "D", correspondingly). EA is adjusted for demographic and genetic ancestry covariates. Counties with sample mean values significantly different from zero after FDR correction at the 0.05 level are marked with an asterisk (*).

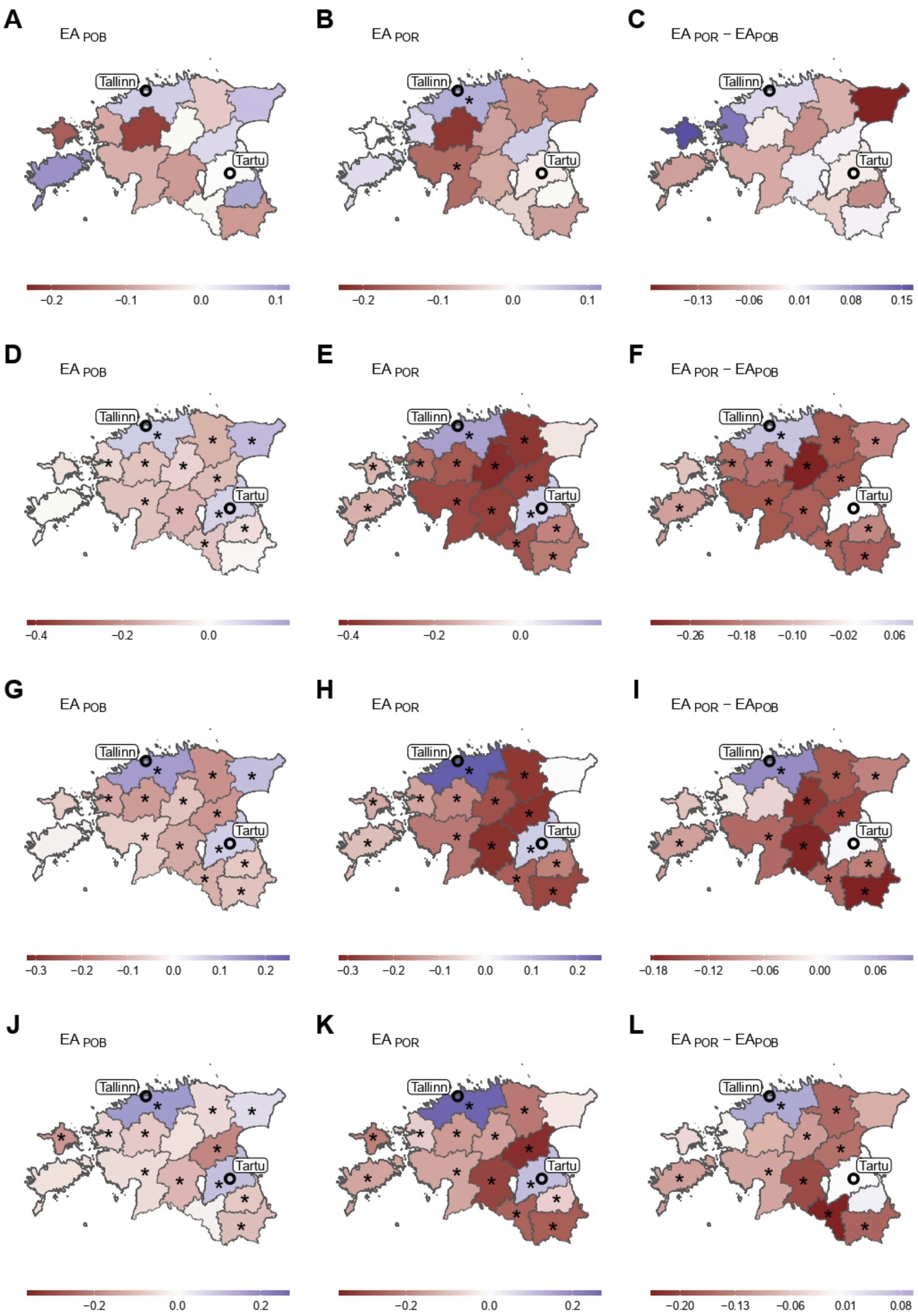
A
EA POB
Tallinn
Tartu
−0.2 −0.1 0.0 0.1
B
EA POR
Tallinn
Tartu
−0.2 −0.1 0.0 0.1
C
EA POR − EAPOB
Tallinn
Tartu
−0.13 −0.06 0.01 0.08 0.15
D
EA POB
Tallinn
Tartu
−0.4 −0.2 0.0
E
EA POR
Tallinn
Tartu
−0.4 −0.2 0.0
F
EA POR − EAPOB
Tallinn
Tartu
−0.26 −0.18 −0.10 −0.02 0.06
G
EA POB
Tallinn
Tartu
−0.3 −0.2 −0.1 0.0 0.1 0.2
H
EA POR
Tallinn
Tartu
−0.3 −0.2 −0.1 0.0 0.1 0.2
I
EA POR − EAPOB
Tallinn
Tartu
−0.18 −0.12 −0.06 0.00 0.06
J
EA POB
Tallinn
Tartu
−0.2 0.0 0.2
K
EA POR
Tallinn
Tartu
−0.2 0.0 0.2
L
EA POR − EAPOB
Tallinn
Tartu
−0.20 −0.13 −0.06 0.01 0.08

**Supplementary Figure 45. EA (years of education) landscape in Estonia among Estonian participants stratified by age.** Age groups were defined as (A-C) 18-24, (D-F) 25-48, (G-I) 49-64, (J-L) 65+. Mean EA of individuals (A, D, G, J) born or (B, E, H, K) residing in each county. (C, F, I, L) Differences between values in “B” and “A” panels (“E”-“D”, “H”-“G”, “K”-“J” correspondingly). EA is adjusted for demographic and genetic ancestry covariates. Counties with sample mean values significantly different from zero after FDR correction at the 0.05 level are marked with an asterisk (*).

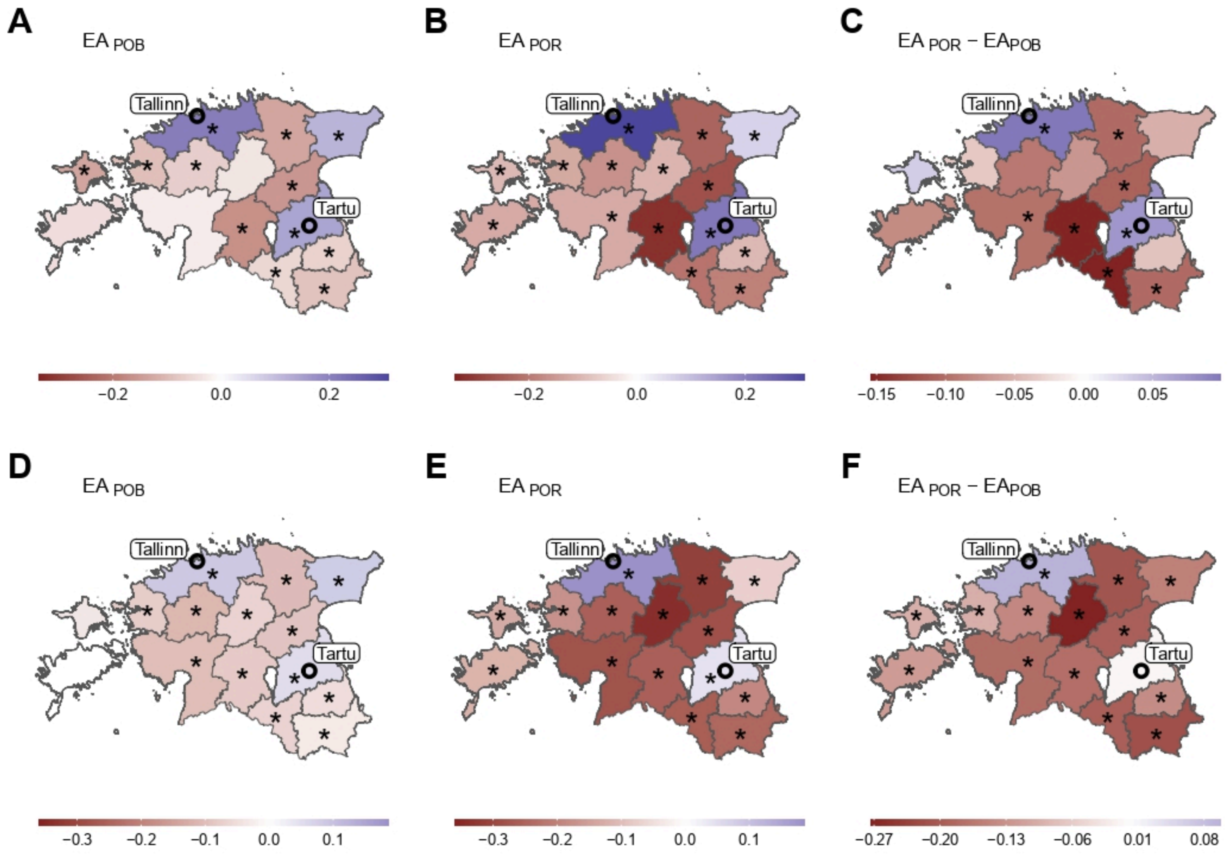

**Supplementary Figure 46. EA (years of education) landscape in Estonia among Estonian participants stratified by year of joining the biobank.** The periods of joining are (A-C) 2001-2016 and (D-F) 2017-2021. Mean EA of individuals (A, D) born or (B, E) residing in each county. (C, F) Differences between values in “B” and “A” panels (“E” and “D”, correspondingly). EA is adjusted for demographic and genetic ancestry covariates. Counties with sample mean values significantly different from zero after FDR correction at the 0.05 level are marked with an asterisk (*).

#### (s)$PGS_{EA}$ values in groups with different migration profiles

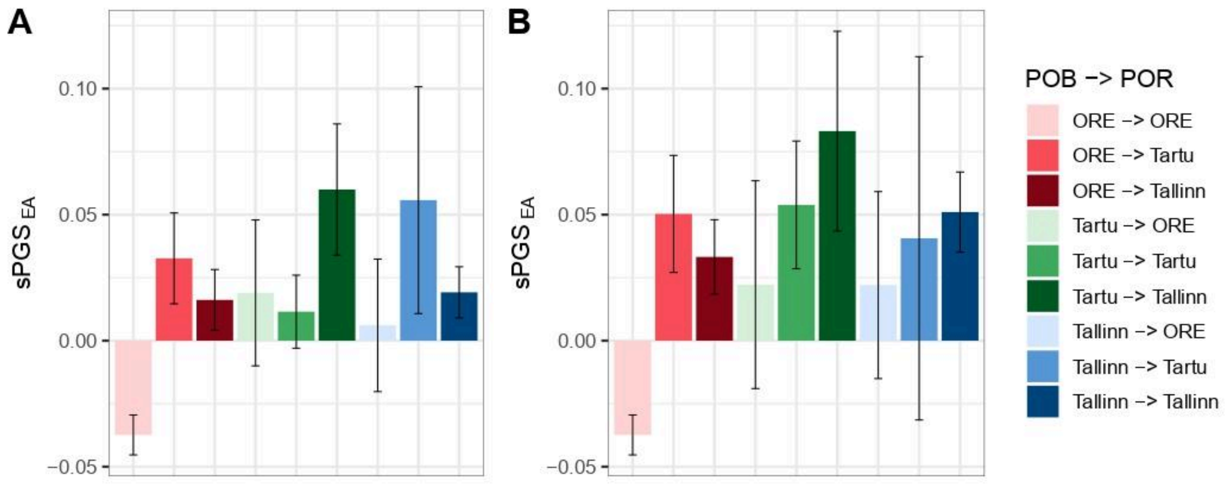

**Supplementary Figure 47. $sPGS_{EA}$ in migration groups among Estonian participants by region of birth (POB) and residence (POR).** (A) County-based analysis where POB and POR refer to Tartu County ("Tartu"), Harju County ("Tallinn") and other counties ("ORE"). (B) City-based analysis, where POB and POR refer to Tartu City ("Tartu"), Tallinn ("Tallinn") and other counties ("ORE"). $sPGS_{EA}$ is adjusted for demographic and genetic ancestry covariates. Error bars correspond to 95% confidence intervals.

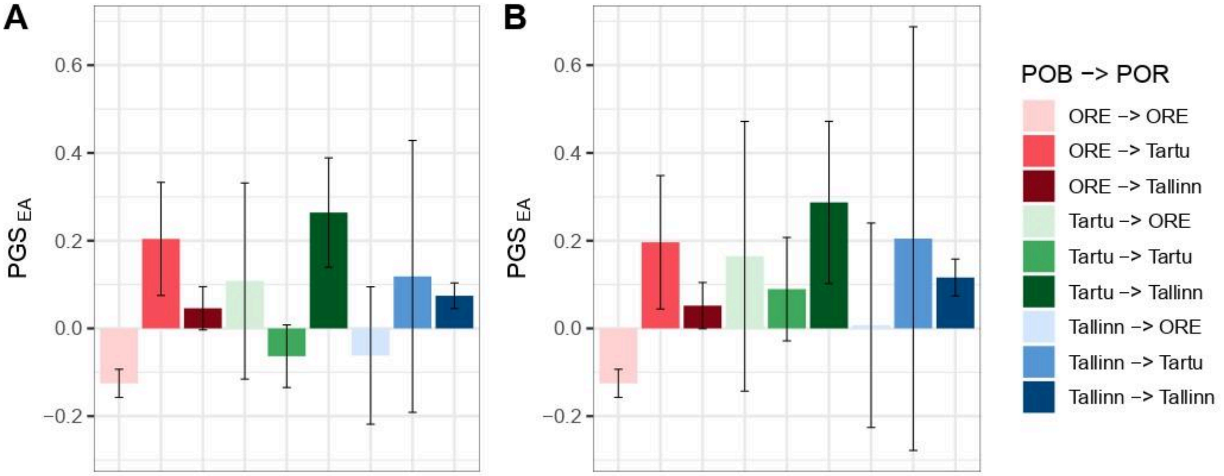

**Supplementary Figure 48. $PGS_{EA}$ in migration groups among Russian participants by region of birth (POB) and residence (POR).** (A) County-based analysis where POB and POR refer to Tartu County ("Tartu"), Harju County ("Tallinn") and other counties ("ORE"). (B) City-based analysis, where POB and POR refer to Tartu City ("Tartu"), Tallinn ("Tallinn") and other counties ("ORE"). $PGS_{EA}$ is adjusted for demographic and genetic ancestry covariates. Error bars correspond to 95% confidence intervals.

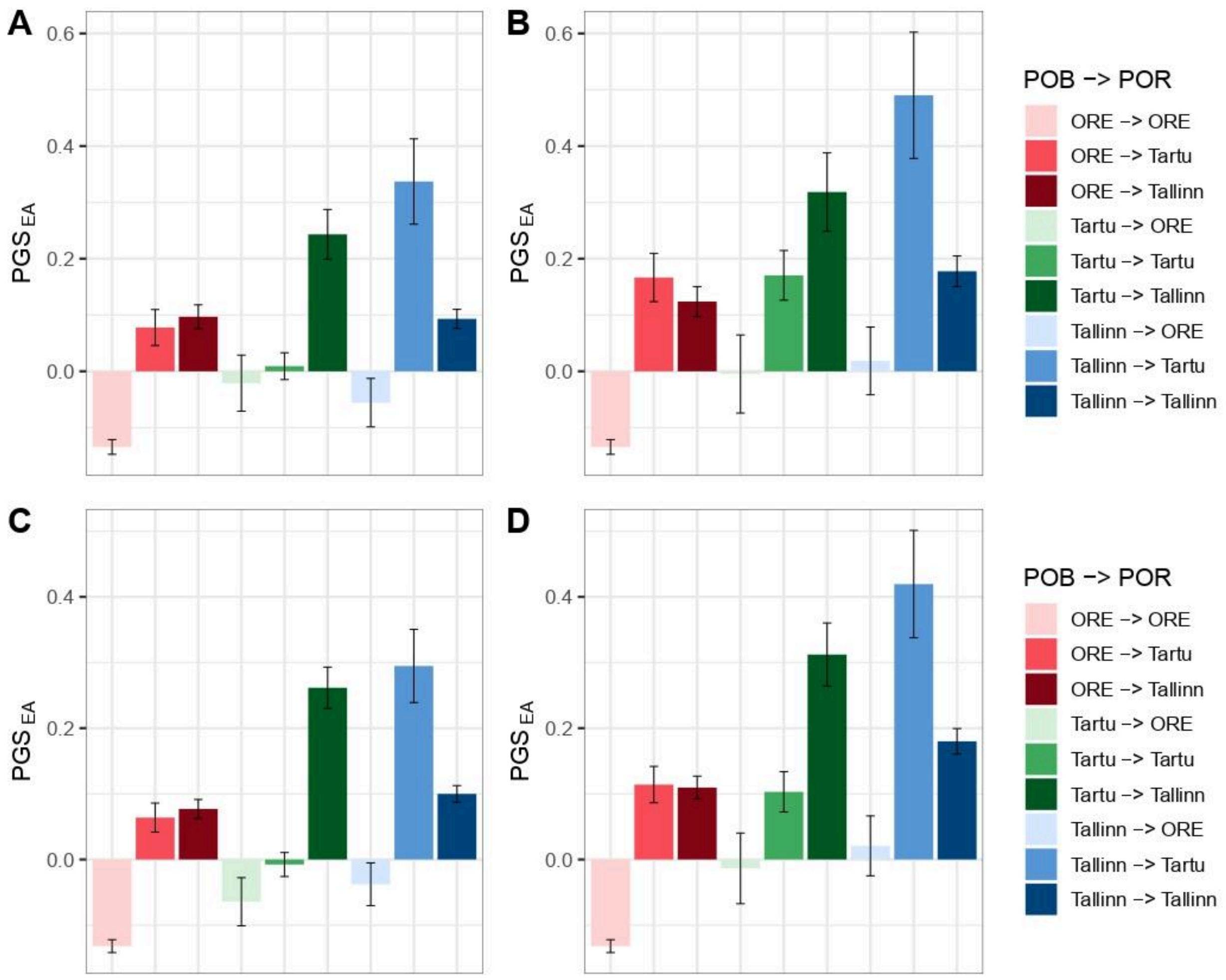

**Supplementary Figure 49. PGS$_{EA}$ in migration groups among (A-B) male and (C-D) female Estonian participants by region of birth (POB) and residence (POR).** (A, C) County-based analysis where POB and POR refer to Tartu County ("Tartu"), Harju County ("Tallinn") and other counties ("ORE"). (B, D) City-based analysis, where POB and POR refer to Tartu City ("Tartu"), Tallinn ("Tallinn") and other counties ("ORE"). PGS$_{EA}$ is adjusted for demographic and genetic ancestry covariates. Error bars correspond to 95% confidence intervals.

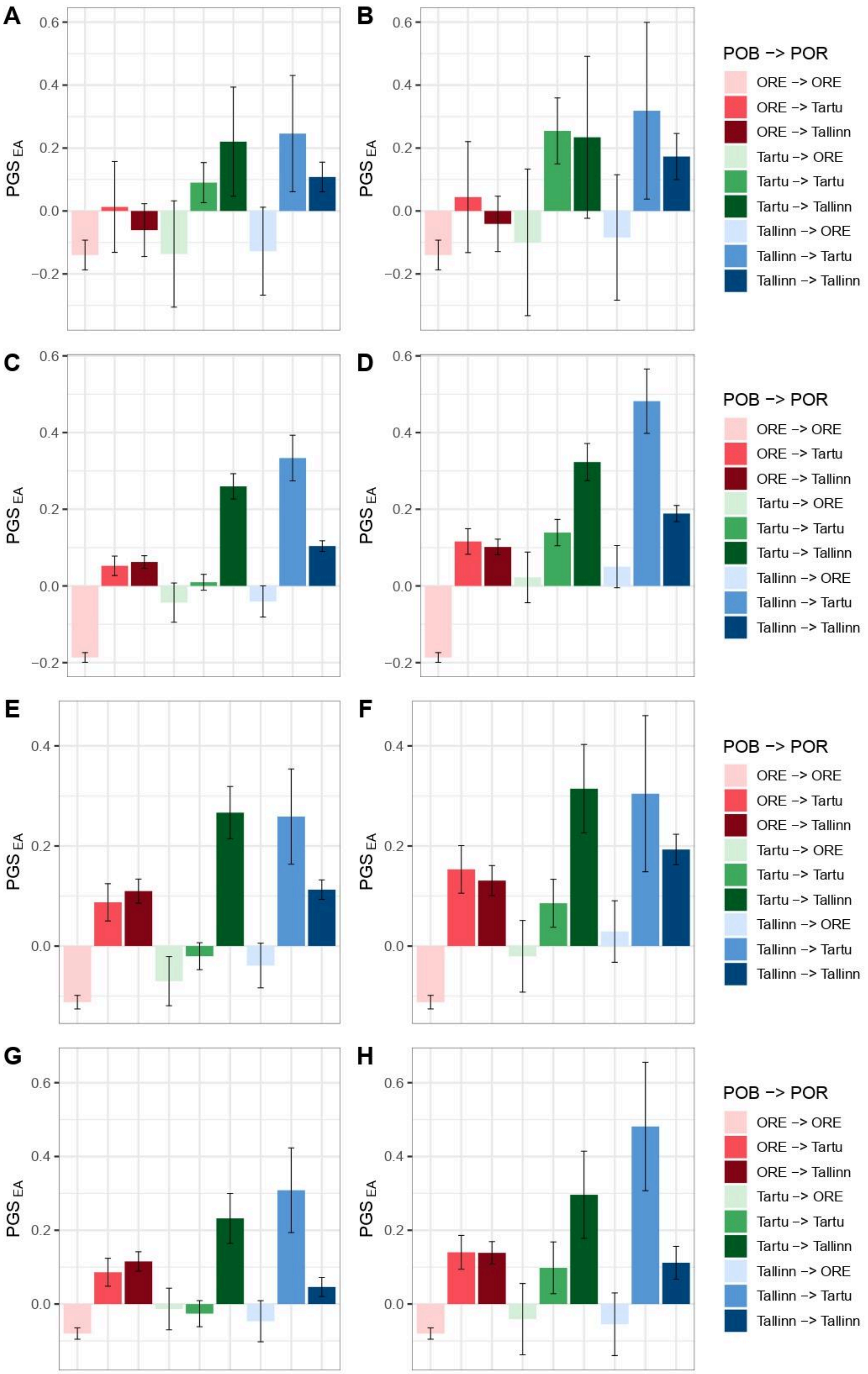
A
B
C
D
E
F
G
H
PGS EA
POB -> POR
ORE -> ORE
ORE -> Tartu
ORE -> Tallinn
Tartu -> ORE
Tartu -> Tartu
Tartu -> Tallinn
Tallinn -> ORE
Tallinn -> Tartu
Tallinn -> Tallinn

**Supplementary Figure 50. $PGS_{EA}$ in migration groups among Estonian participants stratified by age by region of birth (POB) and residence (POR).** Age groups were defined as (A-B) 18-24, (C-D) 25-48, (E-F) 49-64, (G-H) 65+. (A, C, E, G) County-based analysis where POB and POR refer to Tartu County ("Tartu"), Harju County ("Tallinn") and other counties ("ORE"). (B, D, F, H) City-based analysis, where POB and POR refer to Tartu City ("Tartu"), Tallinn ("Tallinn") and other counties ("ORE"). $PGS_{EA}$ is adjusted for demographic and genetic ancestry covariates. Error bars correspond to 95% confidence intervals.

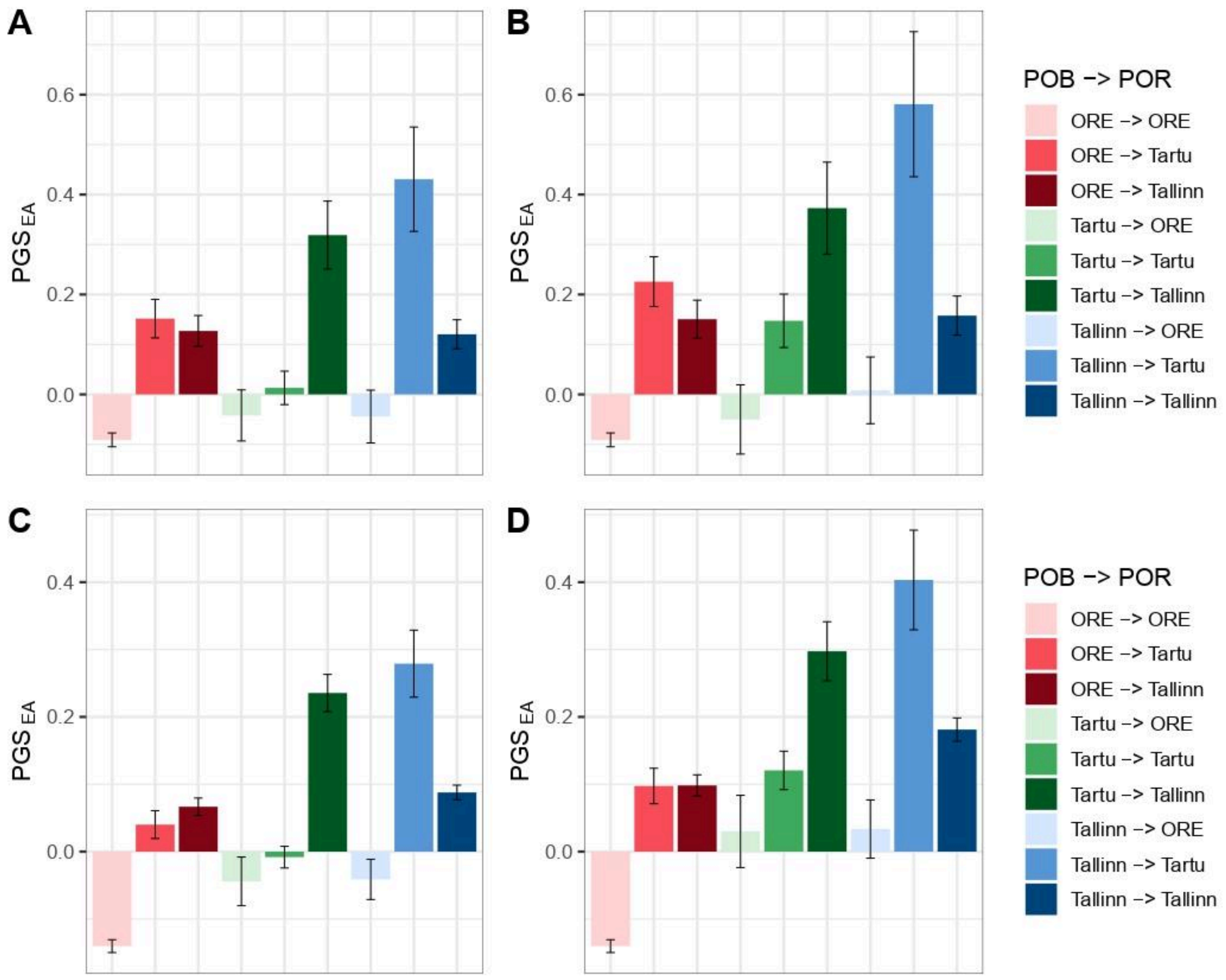

**Supplementary Figure 51. $PGS_{EA}$ in migration groups among Estonian participants stratified by year of joining the biobank by region of birth (POB) and residence (POR).** The periods of joining are (A-B) 2001-2016 and (C-D) 2017-2021. (A, C) County-based analysis where POB and POR refer to Tartu County ("Tartu"), Harju County ("Tallinn") and other counties ("ORE"). (B, D) City-based analysis, where POB and POR refer to Tartu City ("Tartu"), Tallinn ("Tallinn") and other counties ("ORE"). $PGS_{EA}$ is adjusted for demographic and genetic ancestry covariates. Error bars correspond to 95% confidence intervals.

#### Migration direction and (s)$PGS_{EA}$

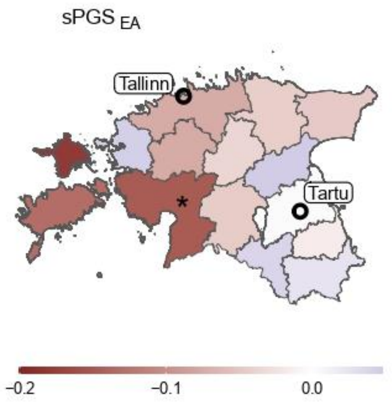

**Supplementary Figure 52. The contrast in mean $sPGS_{EA}$ between residents of Tallinn and Tartu City among Estonian participants by county of birth.** The value for each county corresponds to the mean $sPGS_{EA}$ of individuals born in that county and living in Tartu City subtracted from the mean $sPGS_{EA}$ of individuals born in the same county and living in Tallinn. Individuals born in Tallinn or Tartu City are excluded from the analysis. Counties with significant differences between the migrant groups after FDR correction at level 0.05 are marked with an asterisk (*).

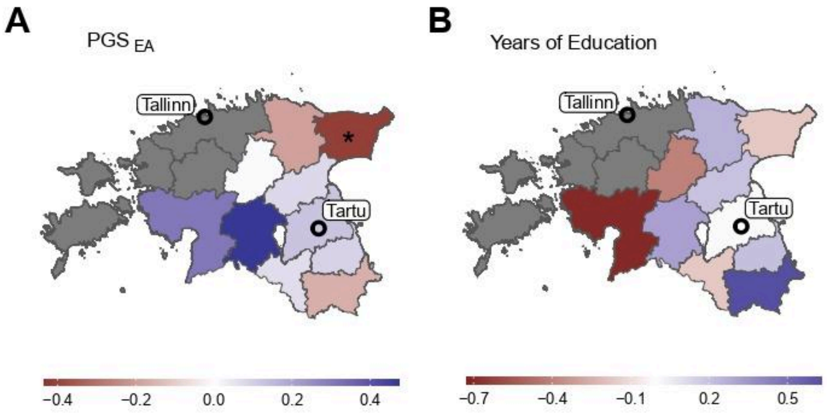

**Supplementary Figure 53. The contrast in mean $PGS_{EA}$ and EA (years of education) between residents of Tallinn and Tartu City among Russian participants by county of birth.** (A) The value for each county corresponds to the mean $PGS_{EA}$ of individuals born in that county and living in Tartu City subtracted from the mean $PGS_{EA}$ of individuals born in the same county and living in Tallinn. Individuals born in Tallinn or Tartu City are excluded from the analysis. (B) The same but for the "years of education" phenotype. Counties with significant differences between the migrant groups after FDR correction at level 0.05 are marked with an asterisk (*). The grey colour means there are fewer than two migrants to Tallinn or Tartu City from the corresponding region in the dataset.

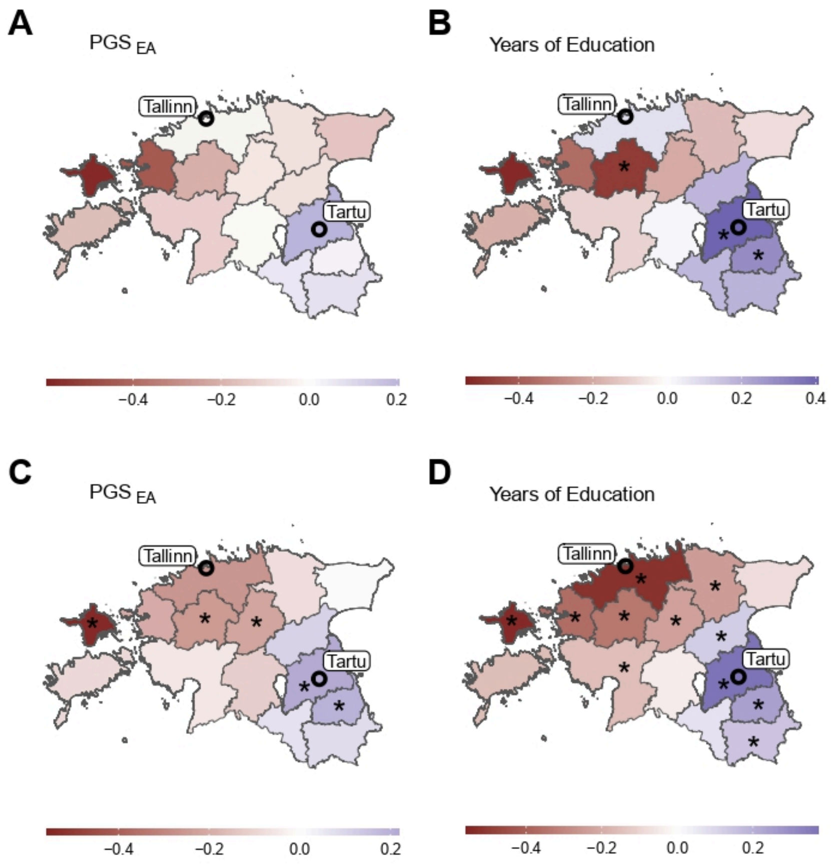

**Supplementary Figure 54. The contrast in mean $PGS_{EA}$ and EA (years of education) between residents of Tallinn and Tartu City among (A-B) male and (C-D) female Estonian participants by county of birth.** (A, C) The value for each county corresponds to the mean $PGS_{EA}$ of individuals born in that county and living in Tartu City subtracted from the mean $PGS_{EA}$ of individuals born in the same county and living in Tallinn. Individuals born in Tallinn or Tartu City are excluded from the analysis. (B, D) The same but for the "years of education" phenotype. Counties with significant differences between the migrant groups after FDR correction at level 0.05 are marked with an asterisk (*).

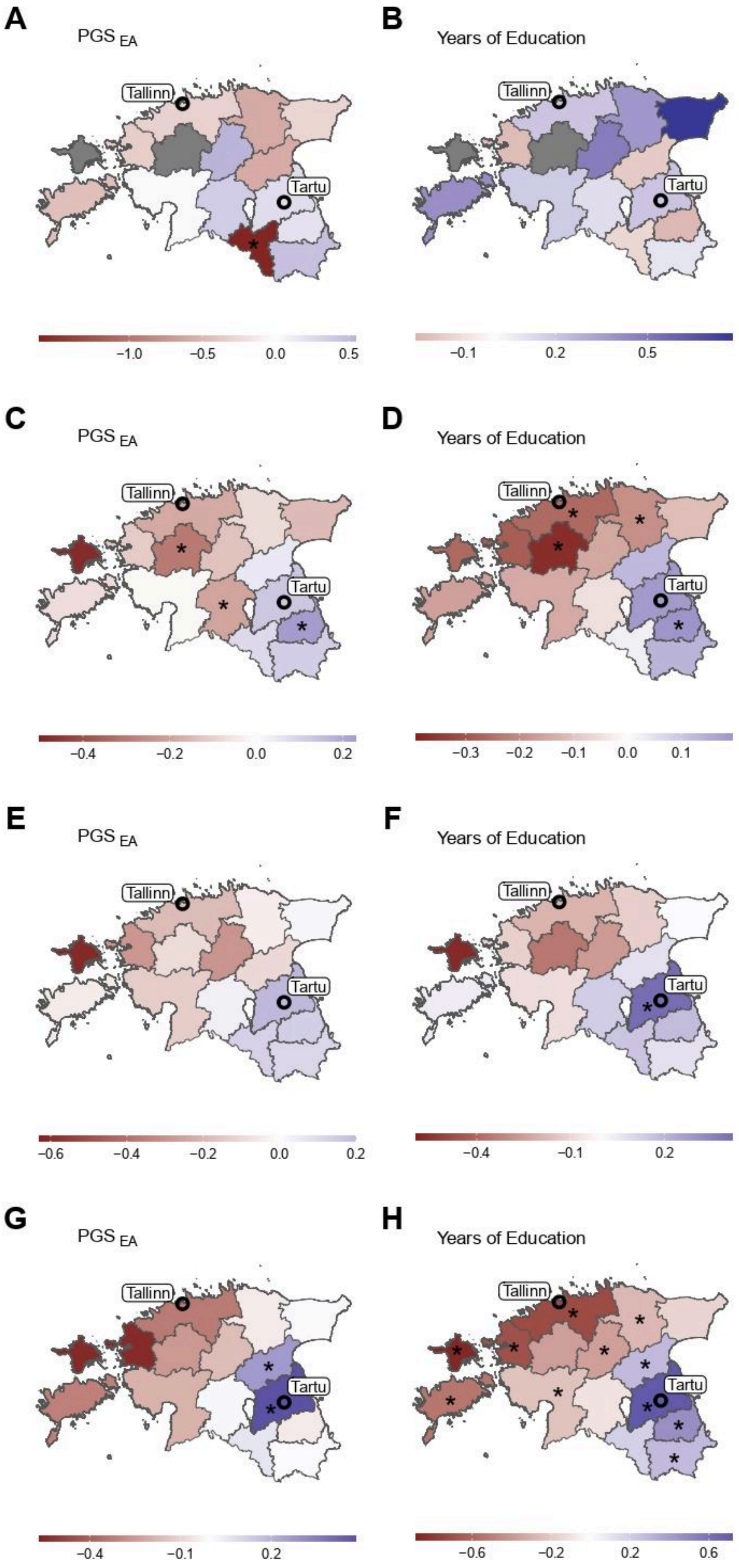
A
PGS EA
Tallinn
Tartu
−1.0 −0.5 0.0 0.5
B
Years of Education
Tallinn
Tartu
−0.1 0.2 0.5
C
PGS EA
Tallinn
Tartu
−0.4 −0.2 0.0 0.2
D
Years of Education
Tallinn
Tartu
−0.3 −0.2 −0.1 0.0 0.1
E
PGS EA
Tallinn
Tartu
−0.6 −0.4 −0.2 0.0 0.2
F
Years of Education
Tallinn
Tartu
−0.4 −0.1 0.2
G
PGS EA
Tallinn
Tartu
−0.4 −0.1 0.2
H
Years of Education
Tallinn
Tartu
−0.6 −0.2 0.2 0.6

**Supplementary Figure 55. The contrast in mean $PGS_{EA}$ and EA (years of education) between residents of Tallinn and Tartu City among Estonian participants stratified by age by county of birth.** Age groups were defined as (A-B) 18-24, (C-D) 25-48, (E-F) 49-64, (G-H) 65+. (A, C, E, G) The value for each county corresponds to the mean $PGS_{EA}$ of individuals born in that county and living in Tartu City subtracted from the mean $PGS_{EA}$ of individuals born in the same county and living in Tallinn. Individuals born in Tallinn or Tartu City are excluded from the analysis. (B, D, F, H) The same but for the "years of education" phenotype. Counties with significant differences between the migrant groups after FDR correction at level 0.05 are marked with an asterisk (*). The grey colour means there are fewer than two migrants to Tartu City or Tallinn from the corresponding region in the dataset.

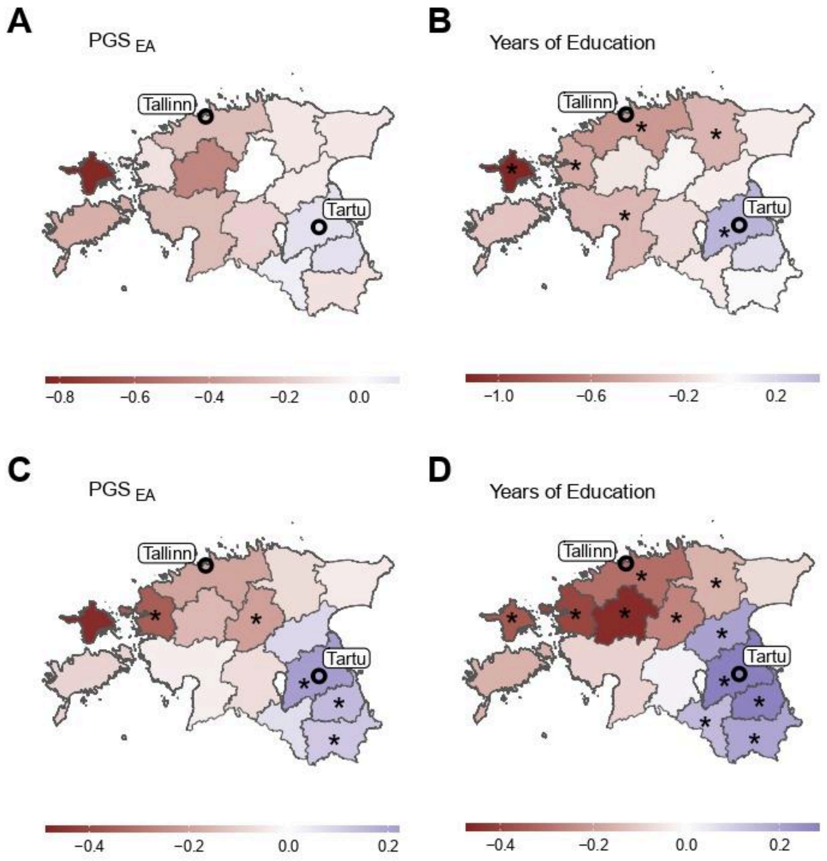

**Supplementary Figure 56. The contrast in mean $PGS_{EA}$ and EA (years of education) between residents of Tallinn and Tartu among Estonian participants stratified by year of joining the biobank by county of birth.** The periods of joining are (A-B) 2001-2016 and (C-D) 2017-2021. (A, C) The value for each county corresponds to the mean $PGS_{EA}$ of individuals born in that county and living in Tartu City subtracted from the mean $PGS_{EA}$ of individuals born in the same county and living in Tallinn. Individuals born in Tallinn or Tartu City are excluded from the analysis. (B, D) The same but for the "years of education" phenotype. Counties with significant differences between the migrant groups after FDR correction at level 0.05 are marked with an asterisk (*).

#### Educational attainment phenotype in groups with different migration profiles

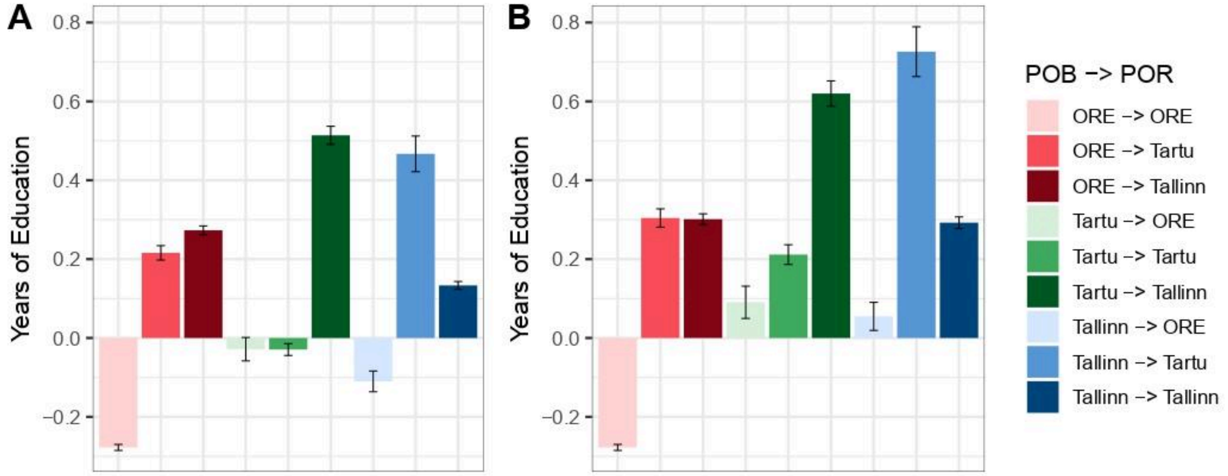

**Supplementary Figure 57. EA (years of education) in migration groups among Estonian participants by region of birth (POB) and residence (POR).** (A) County-based analysis where POB and POR refer to Tartu County ("Tartu"), Harju County ("Tallinn") and other counties ("ORE"). (B) City-based analysis, where POB and POR refer to Tartu City ("Tartu"), Tallinn ("Tallinn") and other counties ("ORE"). EA is adjusted for demographic and genetic ancestry covariates. Error bars correspond to 95% confidence intervals.

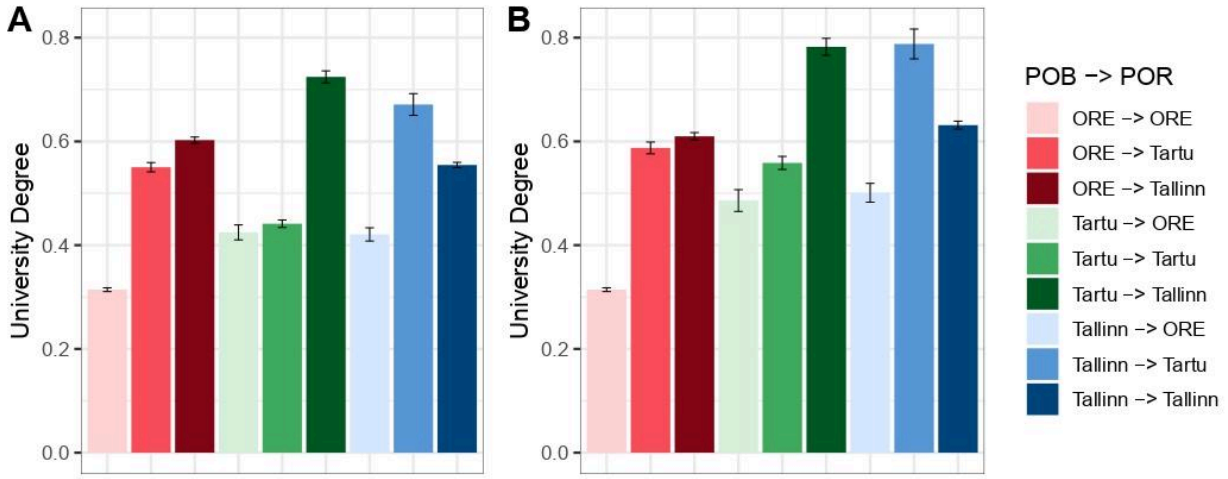

**Supplementary Figure 58. EA (university degree) in migration groups among Estonian participants by region of birth (POB) and residence (POR).** (A) County-based analysis where POB and POR refer to Tartu County ("Tartu"), Harju County ("Tallinn") and other counties ("ORE"). (B) City-based analysis, where POB and POR refer to Tartu City ("Tartu"), Tallinn ("Tallinn") and other counties ("ORE"). Error bars correspond to 95% confidence intervals.

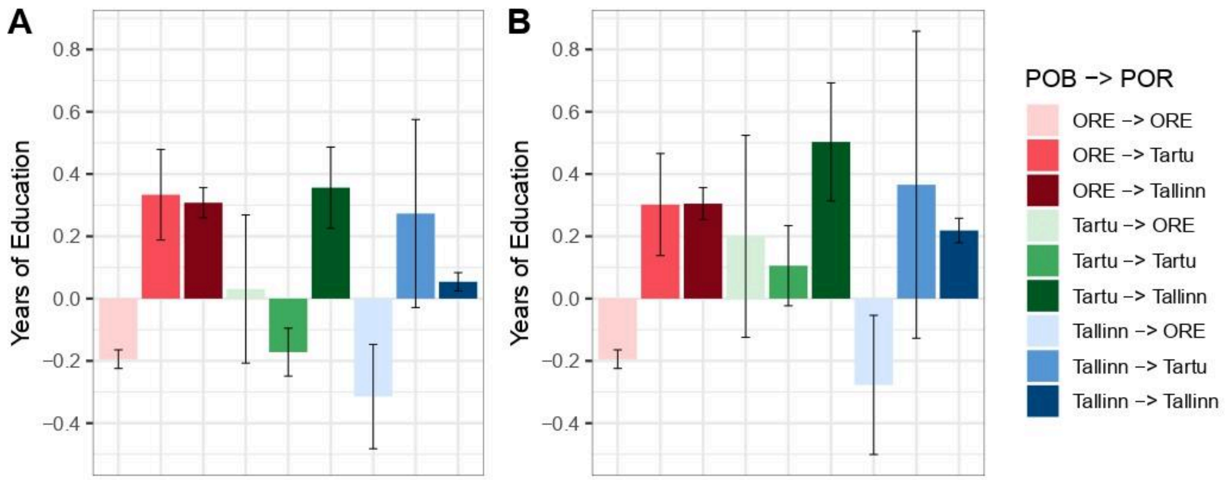

**Supplementary Figure 59. EA (years of education) in migration groups among Russian participants by region of birth (POB) and residence (POR).** (A) County-based analysis where POB and POR refer to Tartu County ("Tartu"), Harju County ("Tallinn") and other counties ("ORE"). (B) City-based analysis, where POB and POR refer to Tartu City ("Tartu"), Tallinn ("Tallinn") and other counties ("ORE"). EA is adjusted for demographic and genetic ancestry covariates. Error bars correspond to 95% confidence intervals.

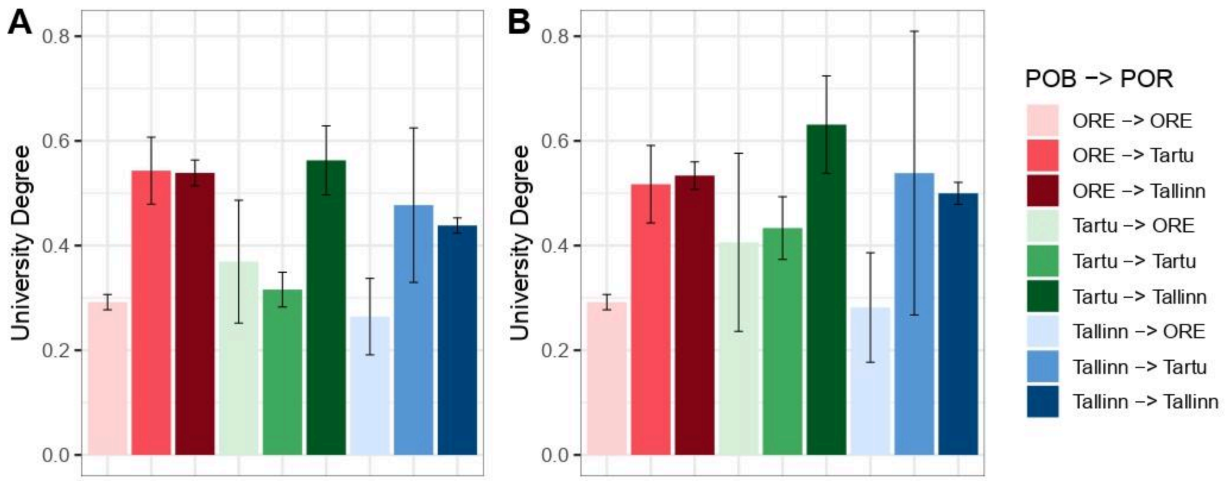

**Supplementary Figure 60. EA (university degree) in migration groups among Russian participants by region of birth (POB) and residence (POR).** (A) County-based analysis where POB and POR refer to Tartu County ("Tartu"), Harju County ("Tallinn") and other counties ("ORE"). (B) City-based analysis, where POB and POR refer to Tartu City ("Tartu"), Tallinn ("Tallinn") and other counties ("ORE"). Error bars correspond to 95% confidence intervals.

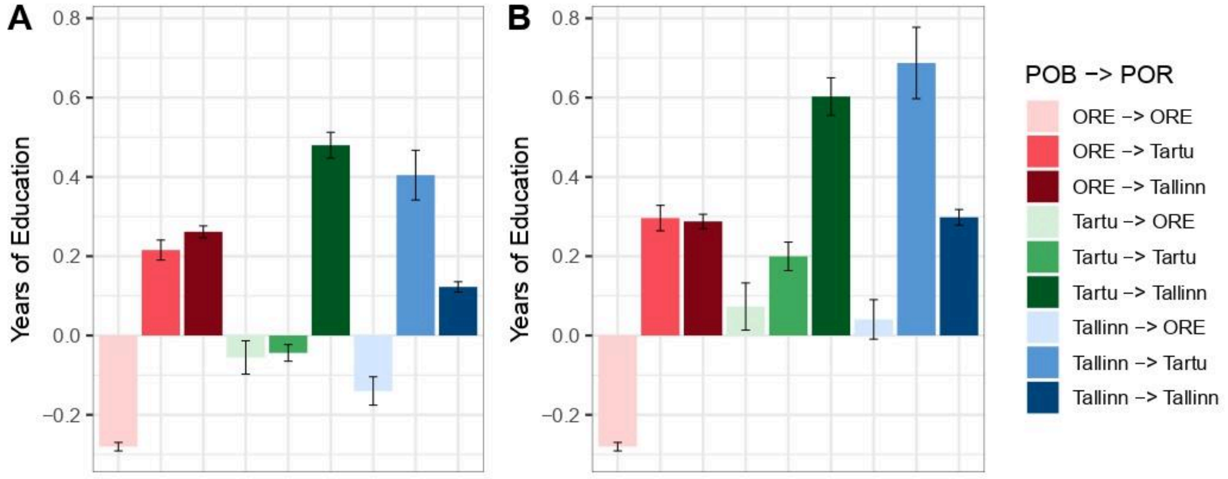

**Supplementary Figure 61. EA (years of education) in migration groups among unrelated Estonian participants by region of birth (POB) and residence (POR).** (A) County-based analysis where POB and POR refer to Tartu County ("Tartu"), Harju County ("Tallinn") and other counties ("ORE"). (B) City-based analysis, where POB and POR refer to Tartu City ("Tartu"), Tallinn ("Tallinn") and other counties ("ORE"). EA is adjusted for demographic and genetic ancestry covariates. Error bars correspond to 95% confidence intervals.

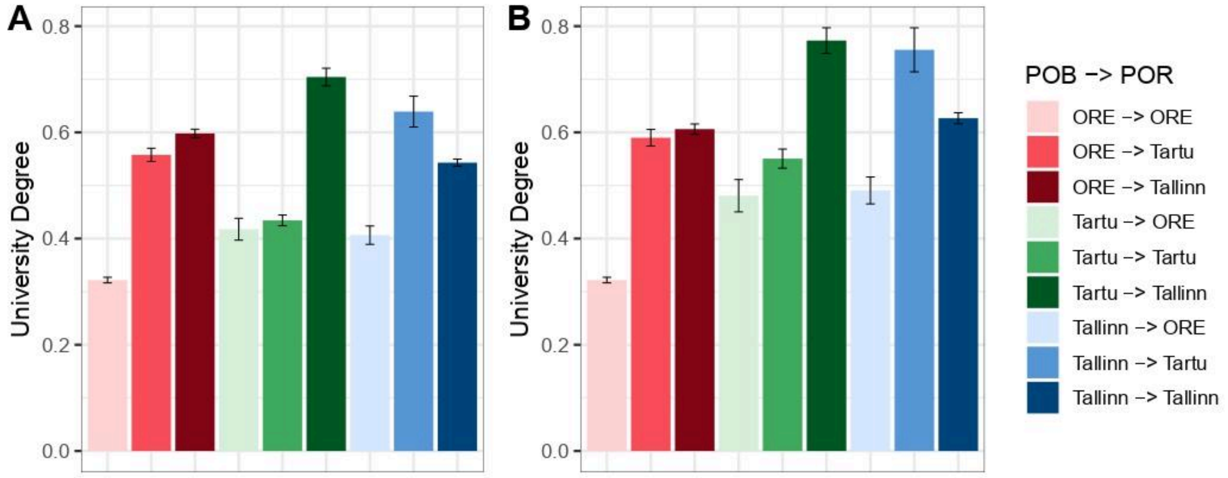

**Supplementary Figure 62. EA (university degree) in migration groups among unrelated Estonian participants by region of birth (POB) and residence (POR).** (A) County-based analysis where POB and POR refer to Tartu County ("Tartu"), Harju County ("Tallinn") and other counties ("ORE"). (B) City-based analysis, where POB and POR refer to Tartu City ("Tartu"), Tallinn ("Tallinn") and other counties ("ORE"). Error bars correspond to 95% confidence intervals.

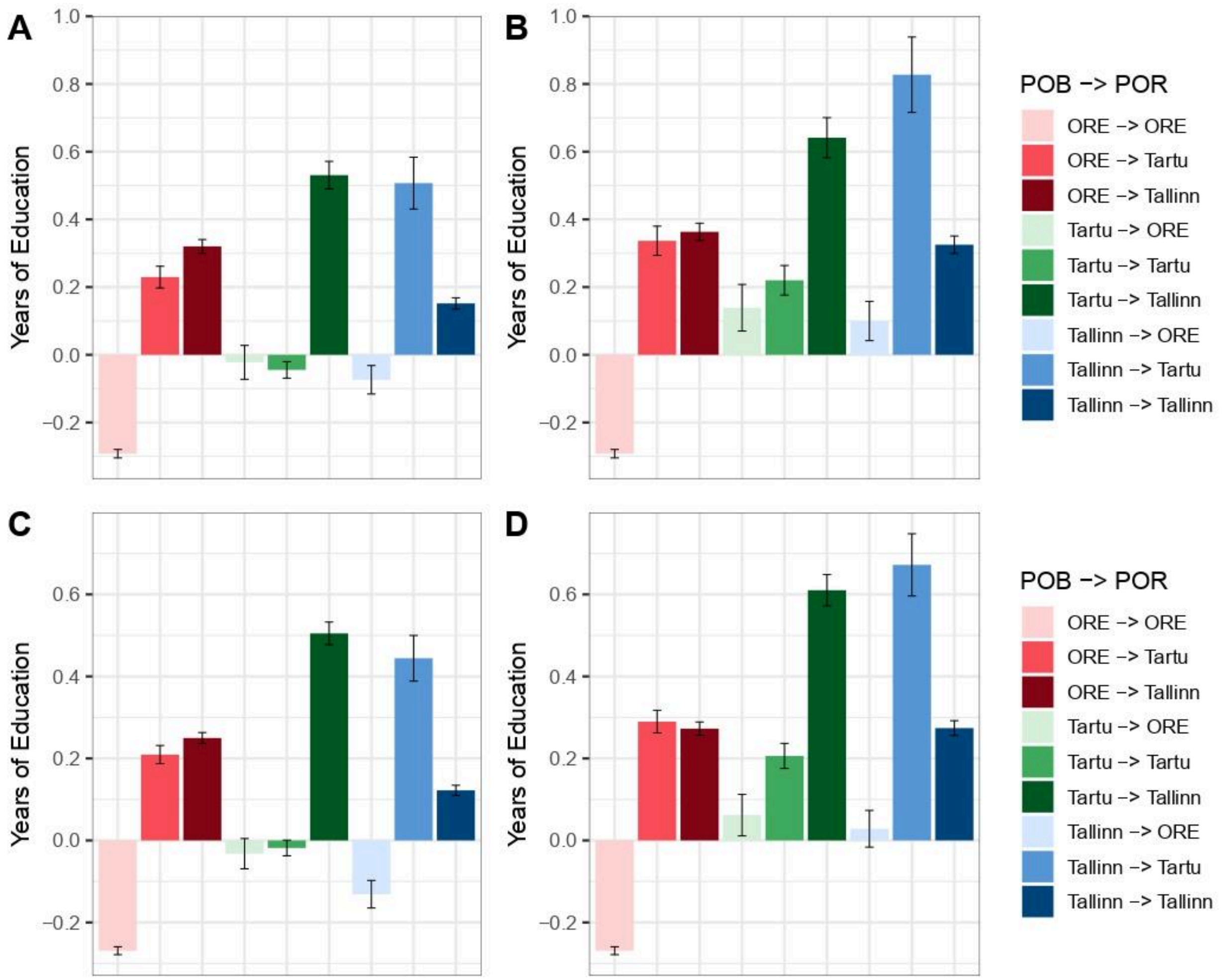

**Supplementary Figure 63. EA (years of education) in migration groups among (A-B) male and (C-D) female Estonian participants by region of birth (POB) and residence (POR).** (A, C) County-based analysis where POB and POR refer to Tartu County ("Tartu"), Harju County ("Tallinn") and other counties ("ORE"). (B, D) City-based analysis, where POB and POR refer to Tartu City ("Tartu"), Tallinn ("Tallinn") and other counties ("ORE"). EA is adjusted for demographic and genetic ancestry covariates. Error bars correspond to 95% confidence intervals.

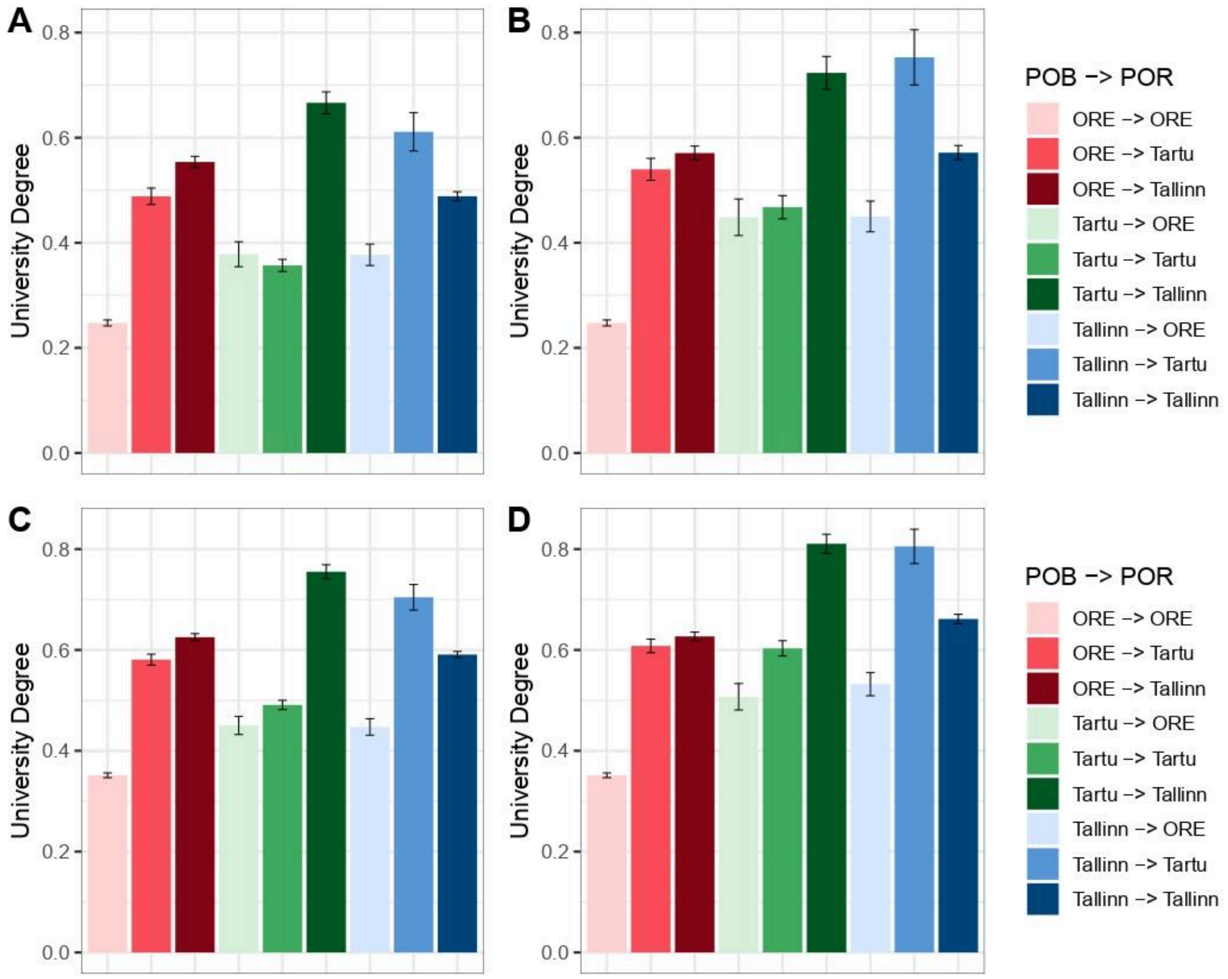

**Supplementary Figure 64. EA (university degree) in migration groups among (A-B) male and (C-D) female Estonian participants by region of birth (POB) and residence (POR).**(A, C) County-based analysis where POB and POR refer to Tartu County ("Tartu"), Harju County ("Tallinn") and other counties ("ORE"). (B, D) City-based analysis, where POB and POR refer to Tartu City ("Tartu"), Tallinn ("Tallinn") and other counties ("ORE"). Error bars correspond to 95% confidence intervals.

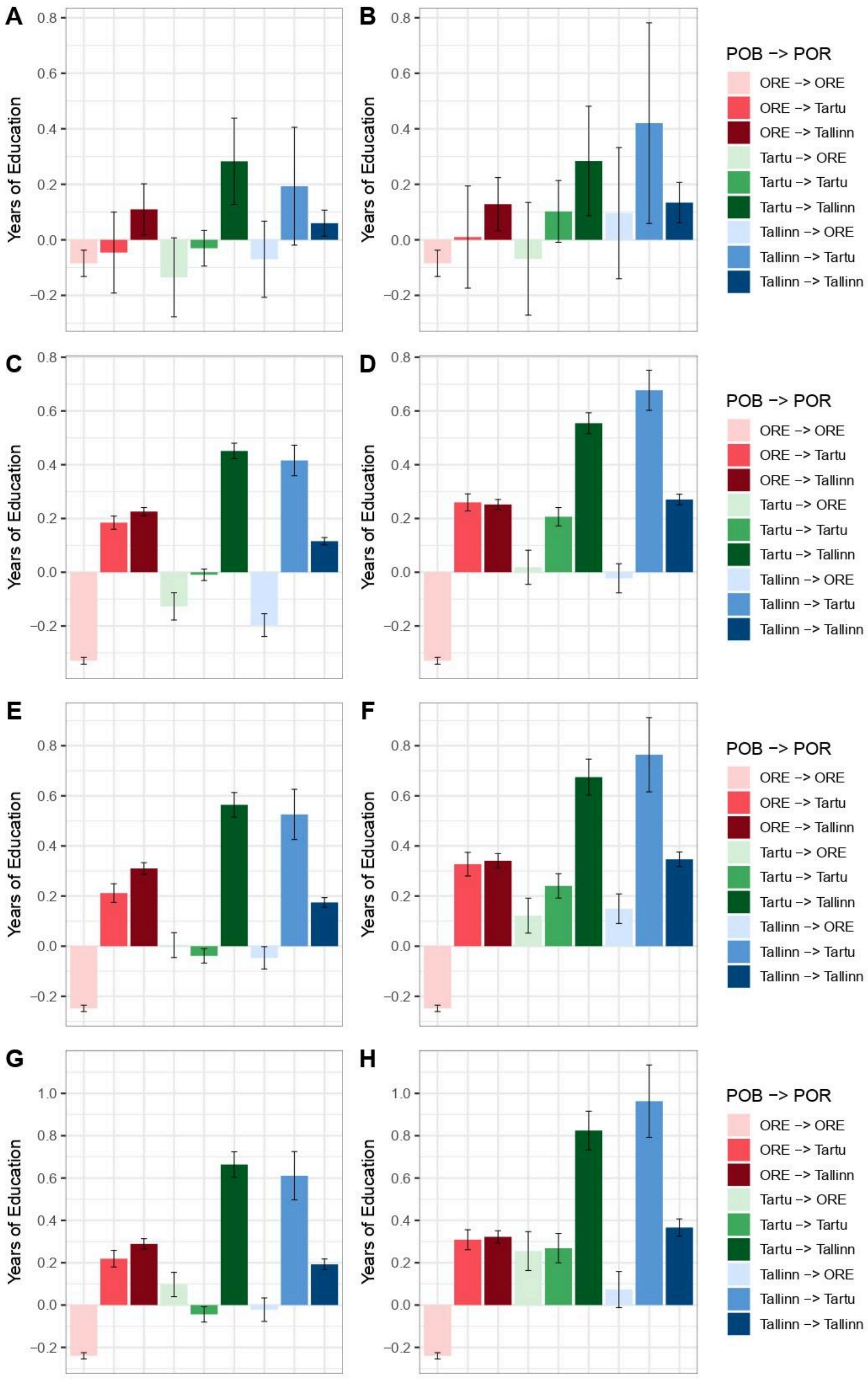

A
B
C
D
E
F
G
H
Years of Education
POB -> POR
ORE -> ORE
ORE -> Tartu
ORE -> Tallinn
Tartu -> ORE
Tartu -> Tartu
Tartu -> Tallinn
Tallinn -> ORE
Tallinn -> Tartu
Tallinn -> Tallinn

**Supplementary Figure 65. EA (years of education) in migration groups among Estonian participants stratified by age by region of birth (POB) and residence (POR).** Age groups were defined as (A-B) 18-24, (C-D) 25-48, (E-F) 49-64, (G-H) 65+. (A, C, E, G) County-based analysis where POB and POR refer to Tartu County ("Tartu"), Harju County ("Tallinn") and other counties ("ORE"). (B, D, F, H) City-based analysis, where POB and POR refer to Tartu City ("Tartu"), Tallinn ("Tallinn") and other counties ("ORE"). EA is adjusted for demographic and genetic ancestry covariates. Error bars correspond to 95% confidence intervals.

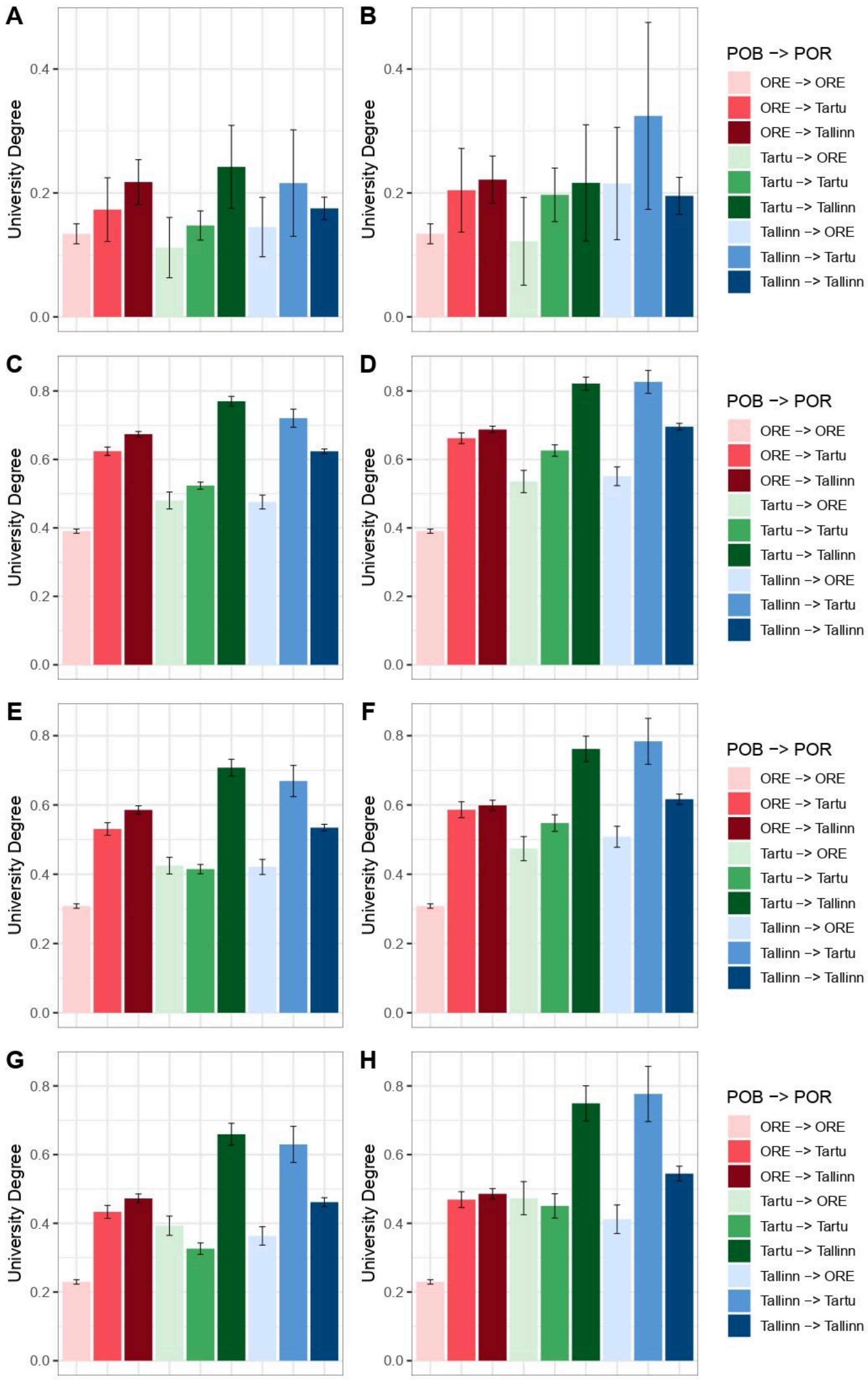
A
B
C
D
E
F
G
H
University Degree
0.0
0.2
0.4
0.6
0.8
POB -> POR
ORE -> ORE
ORE -> Tartu
ORE -> Tallinn
Tartu -> ORE
Tartu -> Tartu
Tartu -> Tallinn
Tallinn -> ORE
Tallinn -> Tartu
Tallinn -> Tallinn

**Supplementary Figure 66. EA (university degree) in migration groups among Estonian participants stratified by age by region of birth (POB) and residence (POR).** Age groups were defined as (A-B) 18-24, (C-D) 25-48, (E-F) 49-64, (G-H) 65+. (A, C, E, G) County-based analysis where POB and POR refer to Tartu County ("Tartu"), Harju County ("Tallinn") and other counties ("ORE"). (B, D, F, H) City-based analysis, where POB and POR refer to Tartu City ("Tartu"), Tallinn ("Tallinn") and other counties ("ORE"). Error bars correspond to 95% confidence intervals.

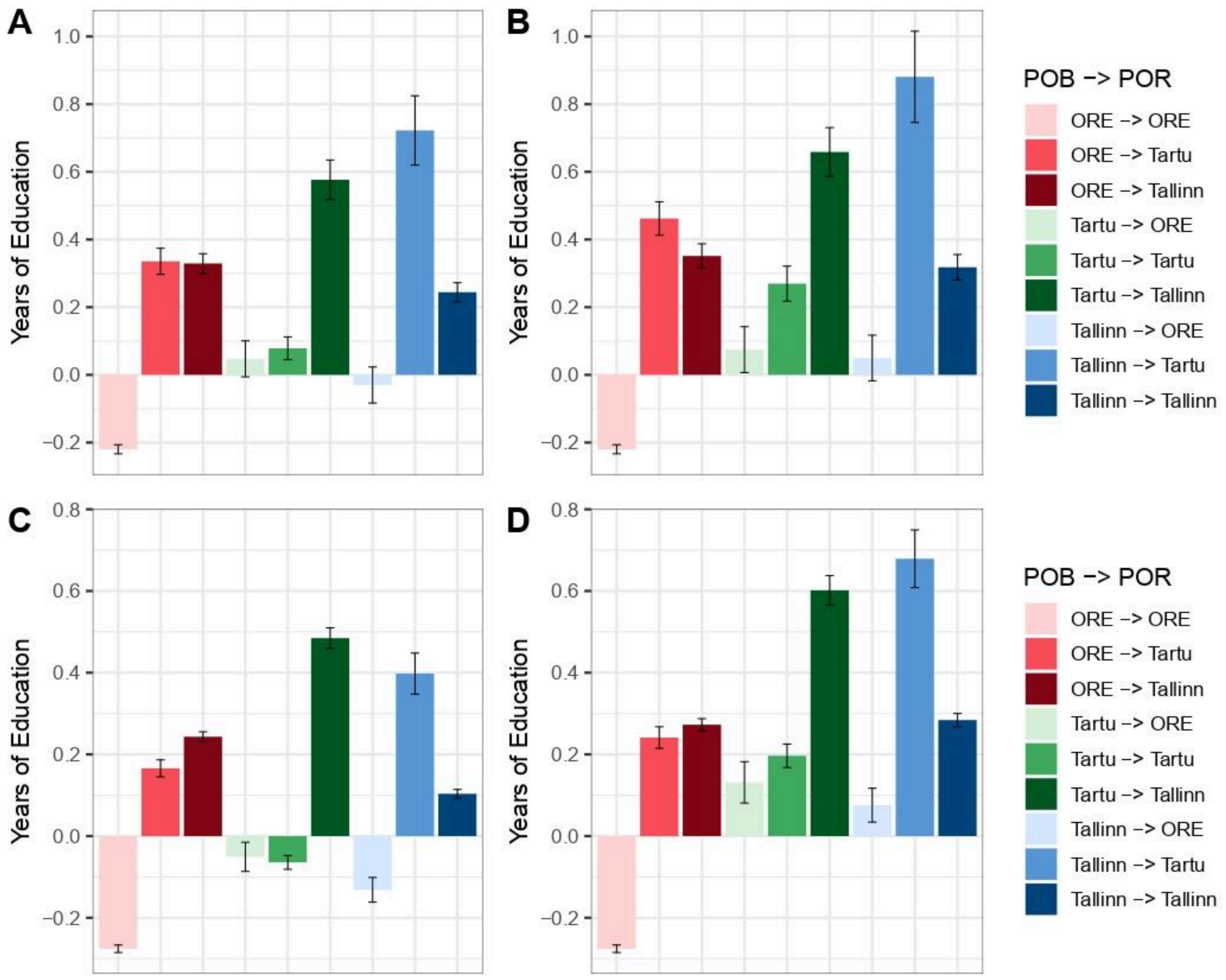

**Supplementary Figure 67. EA (years of education) in migration groups among Estonian participants stratified by year of joining the biobank by region of birth (POB) and residence (POR).** The periods of joining are (A-B) 2001-2016 and (C-D) 2017-2021. (A, C) County-based analysis where POB and POR refer to Tartu County ("Tartu"), Harju County ("Tallinn") and other counties ("ORE"). (B, D) City-based analysis, where POB and POR refer to Tartu City ("Tartu"), Tallinn ("Tallinn") and other counties ("ORE"). EA is adjusted for demographic and genetic ancestry covariates. Error bars correspond to 95% confidence intervals.

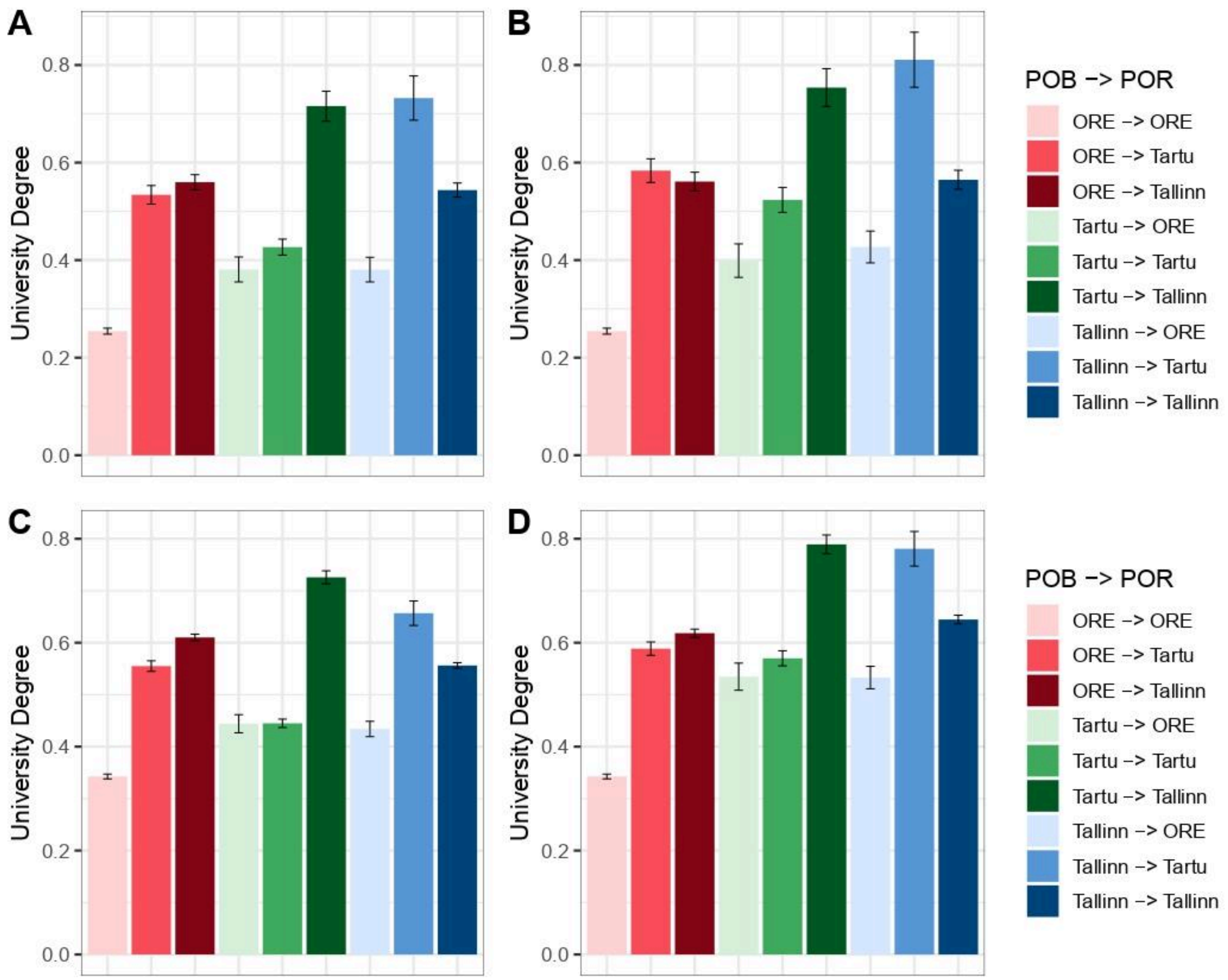

**Supplementary Figure 68. EA (university degree) in migration groups among Estonian participants stratified by year of joining the biobank by region of birth (POB) and residence (POR).** The periods of joining are (A-B) 2001-2016 and (C-D) 2017-2021. (A, C) County-based analysis where POB and POR refer to Tartu County ("Tartu"), Harju County ("Tallinn") and other counties ("ORE"). (B, D) City-based analysis, where POB and POR refer to Tartu City ("Tartu"), Tallinn ("Tallinn") and other counties ("ORE"). Error bars correspond to 95% confidence intervals.

#### (s)$PGS_{EA}$ with EA regressed out in groups with different migration profiles

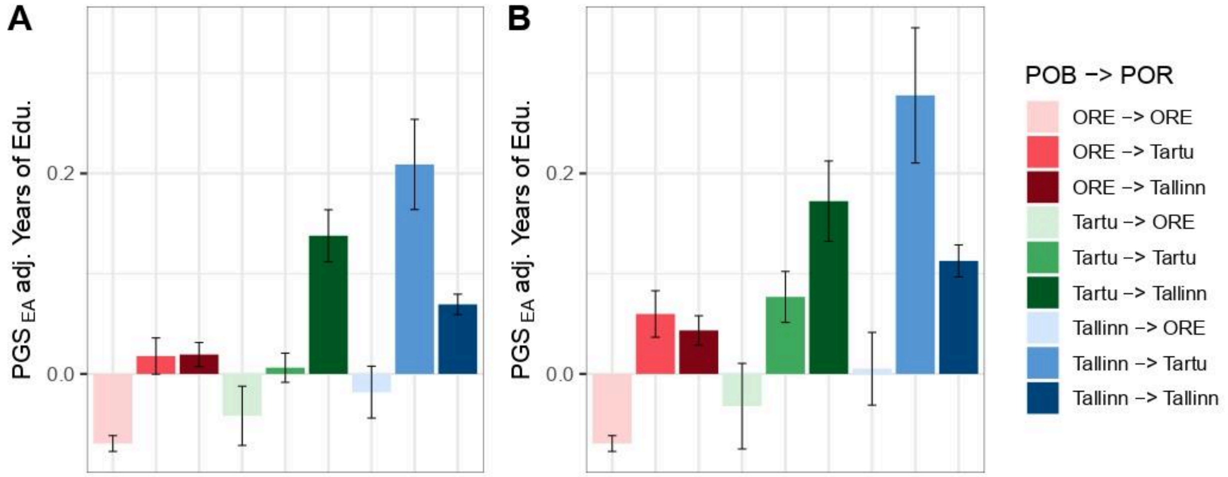

**Supplementary Figure 69. $PGS_{EA}$ with EA (years of education) regressed out in migration groups among Estonian participants by region of birth (POB) and residence (POR).** (A) County-based analysis where POB and POR refer to Tartu County ("Tartu"), Harju County ("Tallinn") and other counties ("ORE"). (B) City-based analysis, where POB and POR refer to Tartu City ("Tartu"), Tallinn ("Tallinn") and other counties ("ORE"). $PGS_{EA}$ is adjusted also for demographic and genetic ancestry covariates. Error bars correspond to 95% confidence intervals.

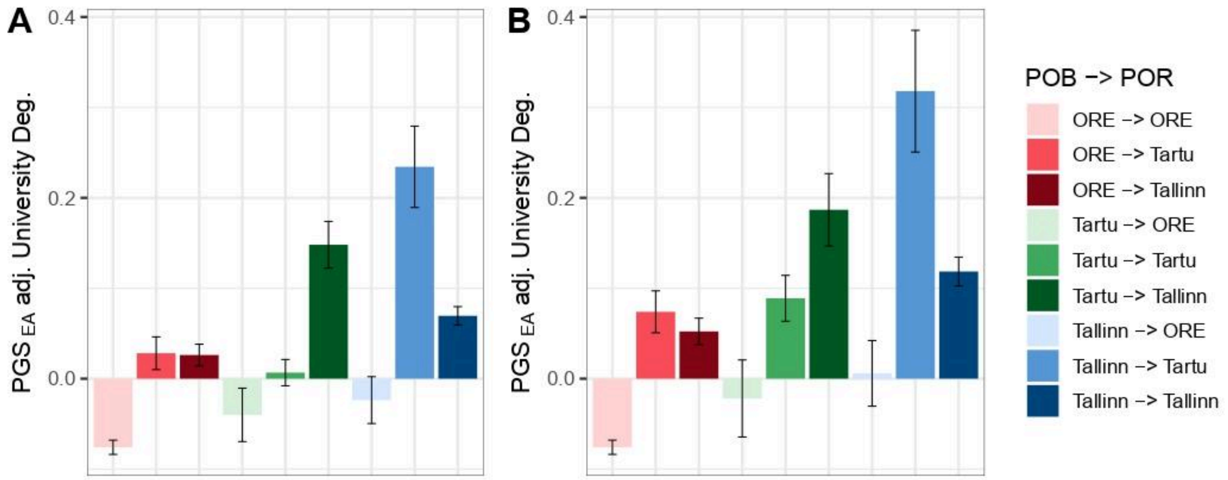

**Supplementary Figure 70. $PGS_{EA}$ with EA (university degree) regressed out in migration groups among Estonian participants by region of birth (POB) and residence (POR).** (A) County-based analysis where POB and POR refer to Tartu County ("Tartu"), Harju County ("Tallinn") and other counties ("ORE"). (B) City-based analysis, where POB and POR refer to Tartu City ("Tartu"), Tallinn ("Tallinn") and other counties ("ORE"). $PGS_{EA}$ is adjusted also for demographic and genetic ancestry covariates. Error bars correspond to 95% confidence intervals.

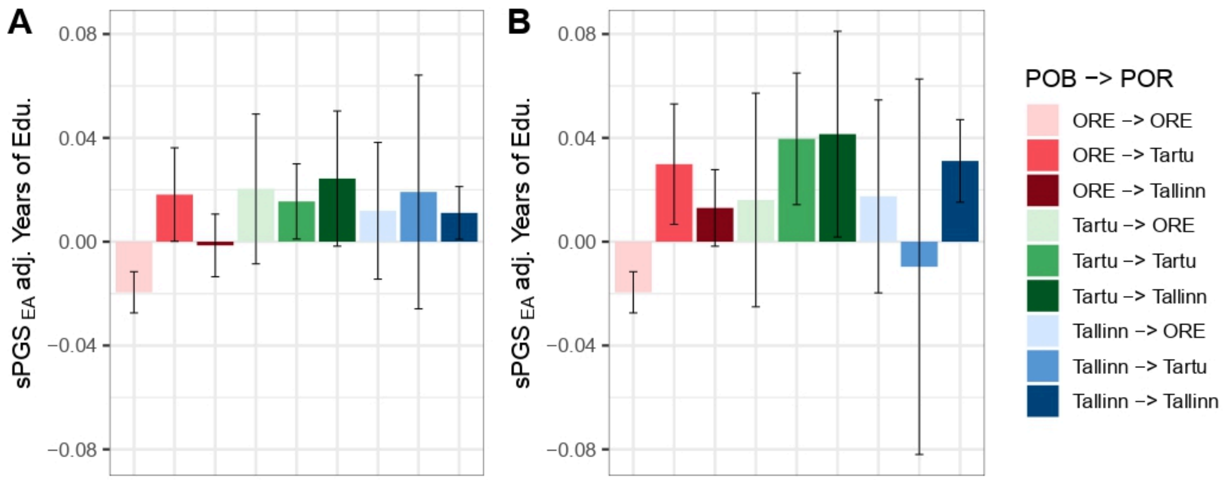

**Supplementary Figure 71. sPGS$_{EA}$ with EA (years of education) regressed out in migration groups among Estonian participants by region of birth (POB) and residence (POR).** (A) County-based analysis where POB and POR refer to Tartu County ("Tartu"), Harju County ("Tallinn") and other counties ("ORE"). (B) City-based analysis, where POB and POR refer to Tartu City ("Tartu"), Tallinn ("Tallinn") and other counties ("ORE"). sPGS$_{EA}$ is adjusted also for demographic and genetic ancestry covariates. Error bars correspond to 95% confidence intervals.

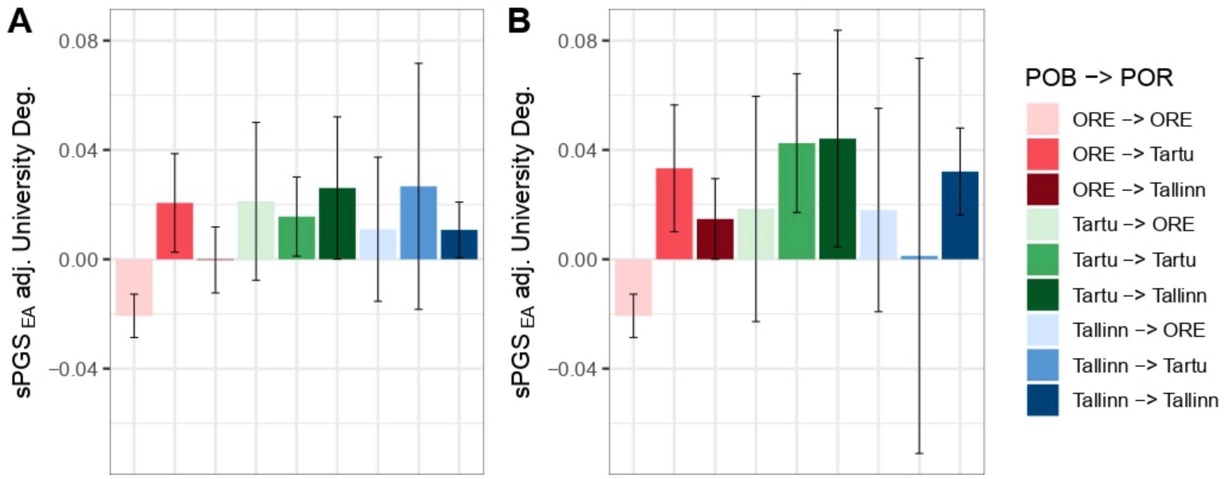

**Supplementary Figure 72. sPGS$_{EA}$ with EA (university degree) regressed out in migration groups among Estonian participants by region of birth (POB) and residence (POR).** (A) County-based analysis where POB and POR refer to Tartu County ("Tartu"), Harju County ("Tallinn") and other counties ("ORE"). (B) City-based analysis, where POB and POR refer to Tartu City ("Tartu"), Tallinn ("Tallinn") and other counties ("ORE"). sPGS$_{EA}$ is adjusted also for demographic and genetic ancestry covariates. Error bars correspond to 95% confidence intervals.

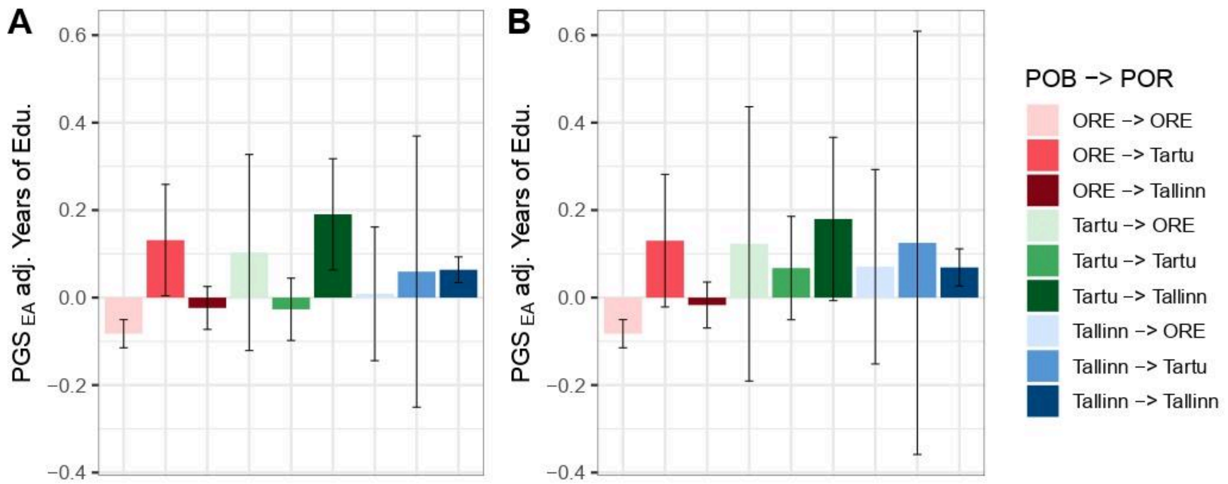

**Supplementary Figure 73. $PGS_{EA}$ with EA (years of education) regressed out in migration groups among Russian participants by region of birth (POB) and residence (POR).** (A) County-based analysis where POB and POR refer to Tartu County ("Tartu"), Harju County ("Tallinn") and other counties ("ORE"). (B) City-based analysis, where POB and POR refer to Tartu City ("Tartu"), Tallinn ("Tallinn") and other counties ("ORE"). $PGS_{EA}$ is adjusted also for demographic and genetic ancestry covariates. Error bars correspond to 95% confidence intervals.

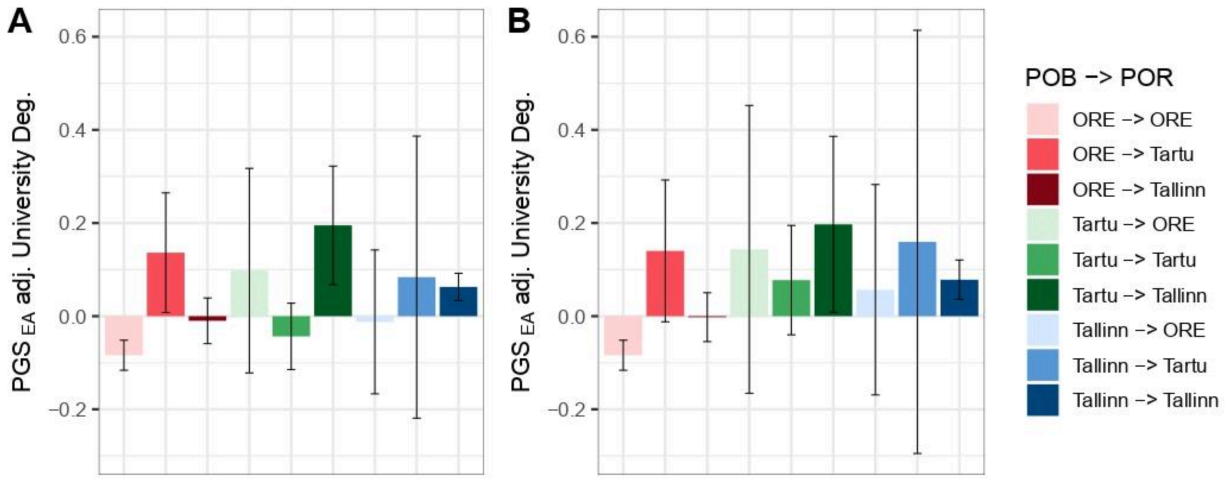

**Supplementary Figure 74. $PGS_{EA}$ with EA (university degree) regressed out in migration groups among Russian participants by region of birth (POB) and residence (POR).** (A) County-based analysis where POB and POR refer to Tartu County ("Tartu"), Harju County ("Tallinn") and other counties ("ORE"). (B) City-based analysis, where POB and POR refer to Tartu City ("Tartu"), Tallinn ("Tallinn") and other counties ("ORE"). $PGS_{EA}$ is adjusted also for demographic and genetic ancestry covariates. Error bars correspond to 95% confidence intervals.

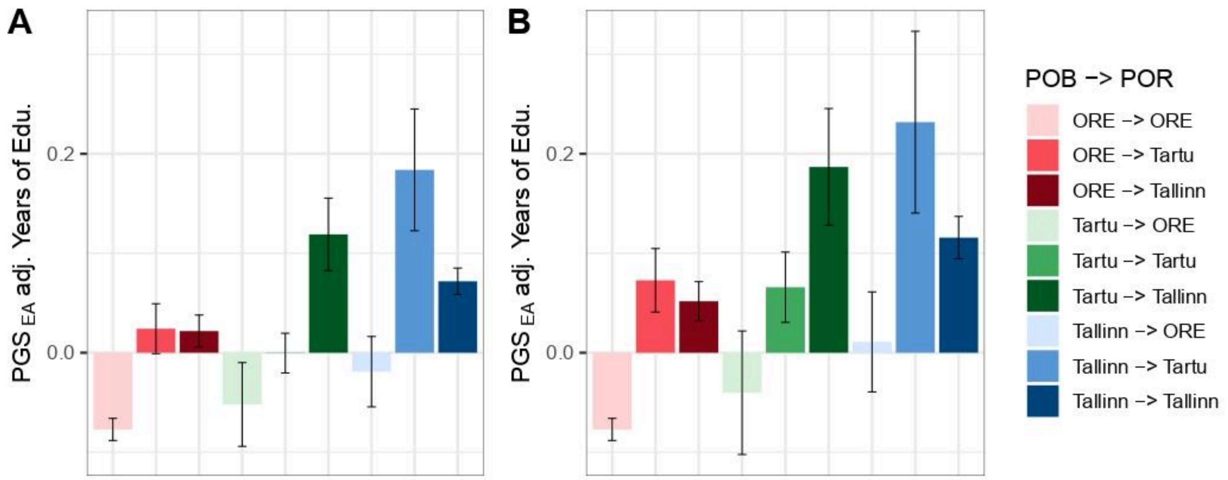

**Supplementary Figure 75. $PGS_{EA}$ with EA (years of education) regressed out in migration groups among unrelated Estonian participants by region of birth (POB) and residence (POR).** (A) County-based analysis where POB and POR refer to Tartu County ("Tartu"), Harju County ("Tallinn") and other counties ("ORE"). (B) City-based analysis, where POB and POR refer to Tartu City ("Tartu"), Tallinn ("Tallinn") and other counties ("ORE"). $PGS_{EA}$ is adjusted also for demographic and genetic ancestry covariates. Error bars correspond to 95% confidence intervals.

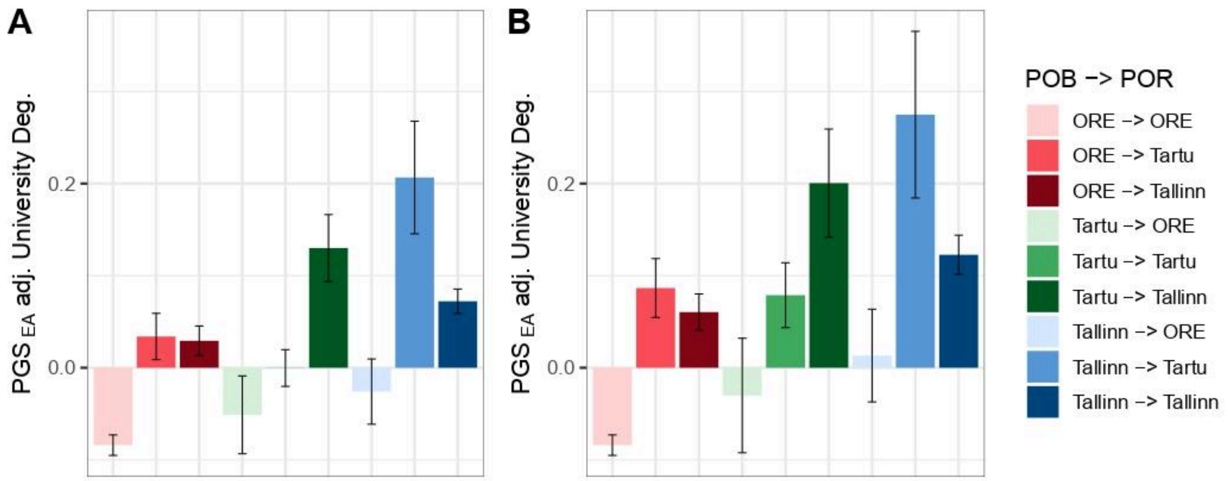

**Supplementary Figure 76. $PGS_{EA}$ with EA (university degree) regressed out in migration groups among unrelated Estonian participants by region of birth (POB) and residence (POR).** (A) County-based analysis where POB and POR refer to Tartu County ("Tartu"), Harju County ("Tallinn") and other counties ("ORE"). (B) City-based analysis, where POB and POR refer to Tartu City ("Tartu"), Tallinn ("Tallinn") and other counties ("ORE"). $PGS_{EA}$ is adjusted also for demographic and genetic ancestry covariates. Error bars correspond to 95% confidence intervals.

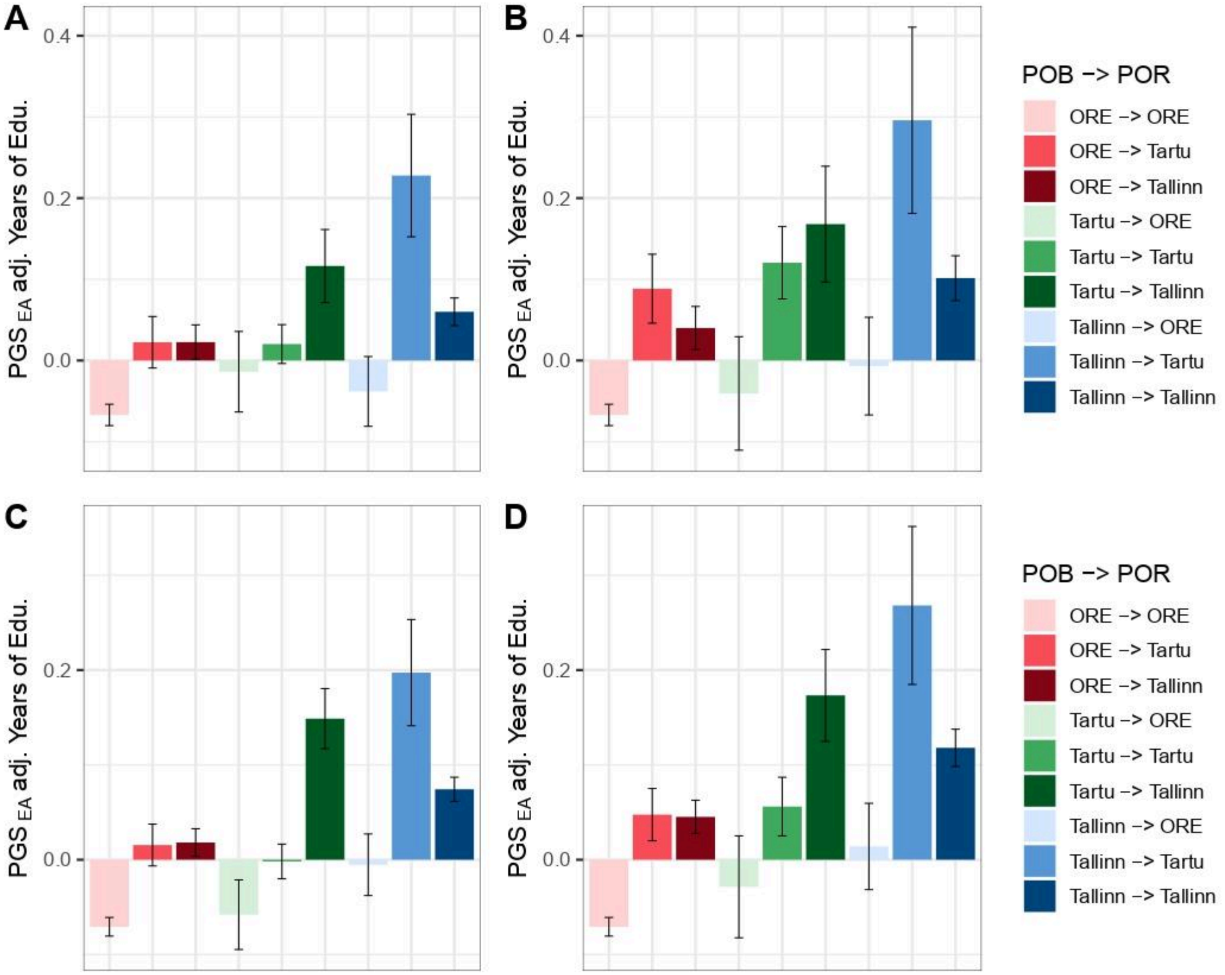

**Supplementary Figure 77. $PGS_{EA}$ with EA (years of education) regressed out in migration groups among (A-B) male and (C-D) female Estonian participants by region of birth (POB) and residence (POR).** (A, C) County-based analysis where POB and POR refer to Tartu County ("Tartu"), Harju County ("Tallinn") and other counties ("ORE"). (B, D) City-based analysis, where POB and POR refer to Tartu City ("Tartu"), Tallinn ("Tallinn") and other counties ("ORE"). $PGS_{EA}$ is adjusted also for demographic and genetic ancestry covariates. Error bars correspond to 95% confidence intervals.

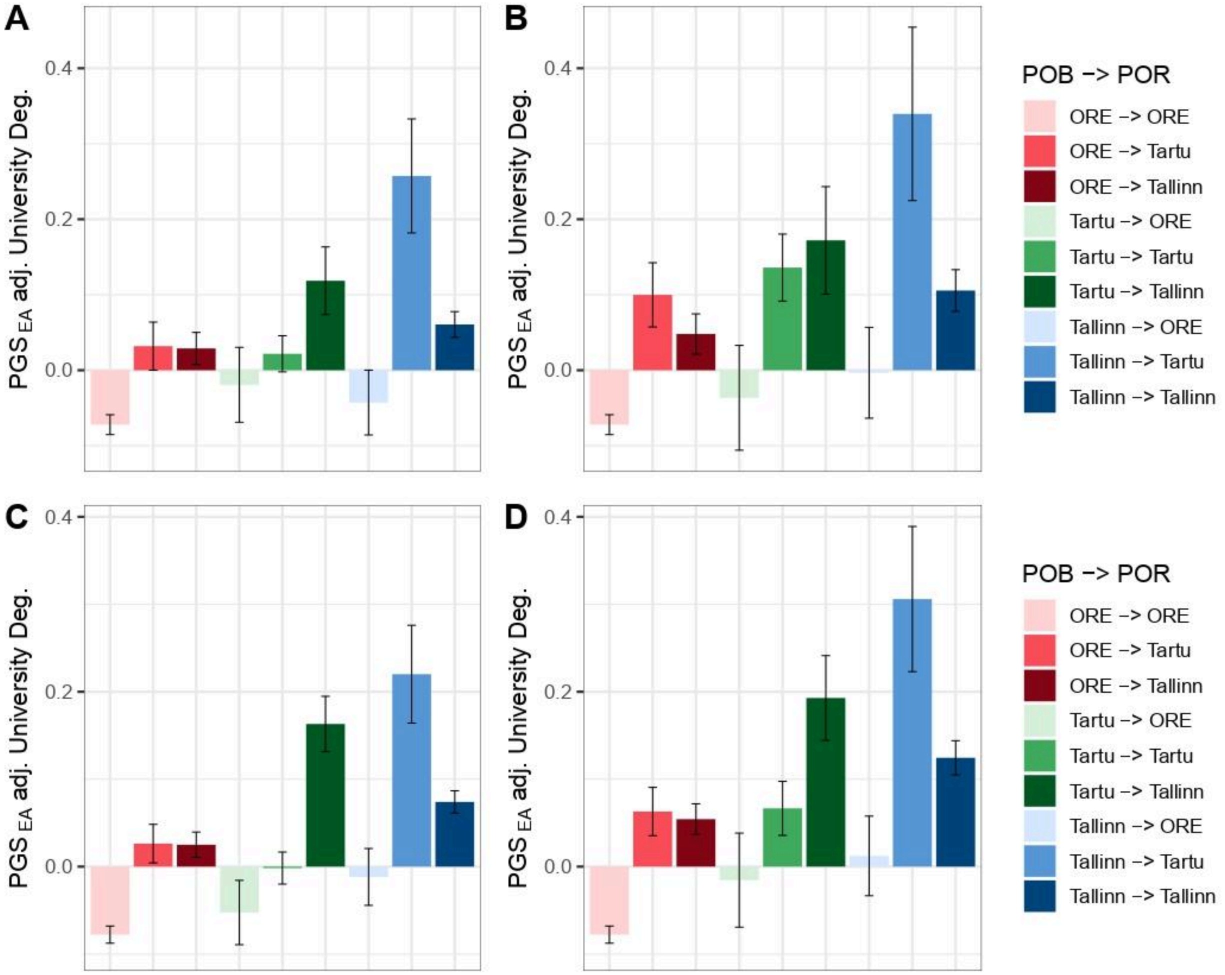

**Supplementary Figure 78. $PGS_{EA}$ with EA (university degree) regressed out in migration groups among (A-B) male and (C-D) female Estonian participants by region of birth (POB) and residence (POR).** (A, C) County-based analysis where POB and POR refer to Tartu County ("Tartu"), Harju County ("Tallinn") and other counties ("ORE"). (B, D) City-based analysis, where POB and POR refer to Tartu City ("Tartu"), Tallinn ("Tallinn") and other counties ("ORE"). $PGS_{EA}$ is adjusted also for demographic and genetic ancestry covariates. Error bars correspond to 95% confidence intervals.

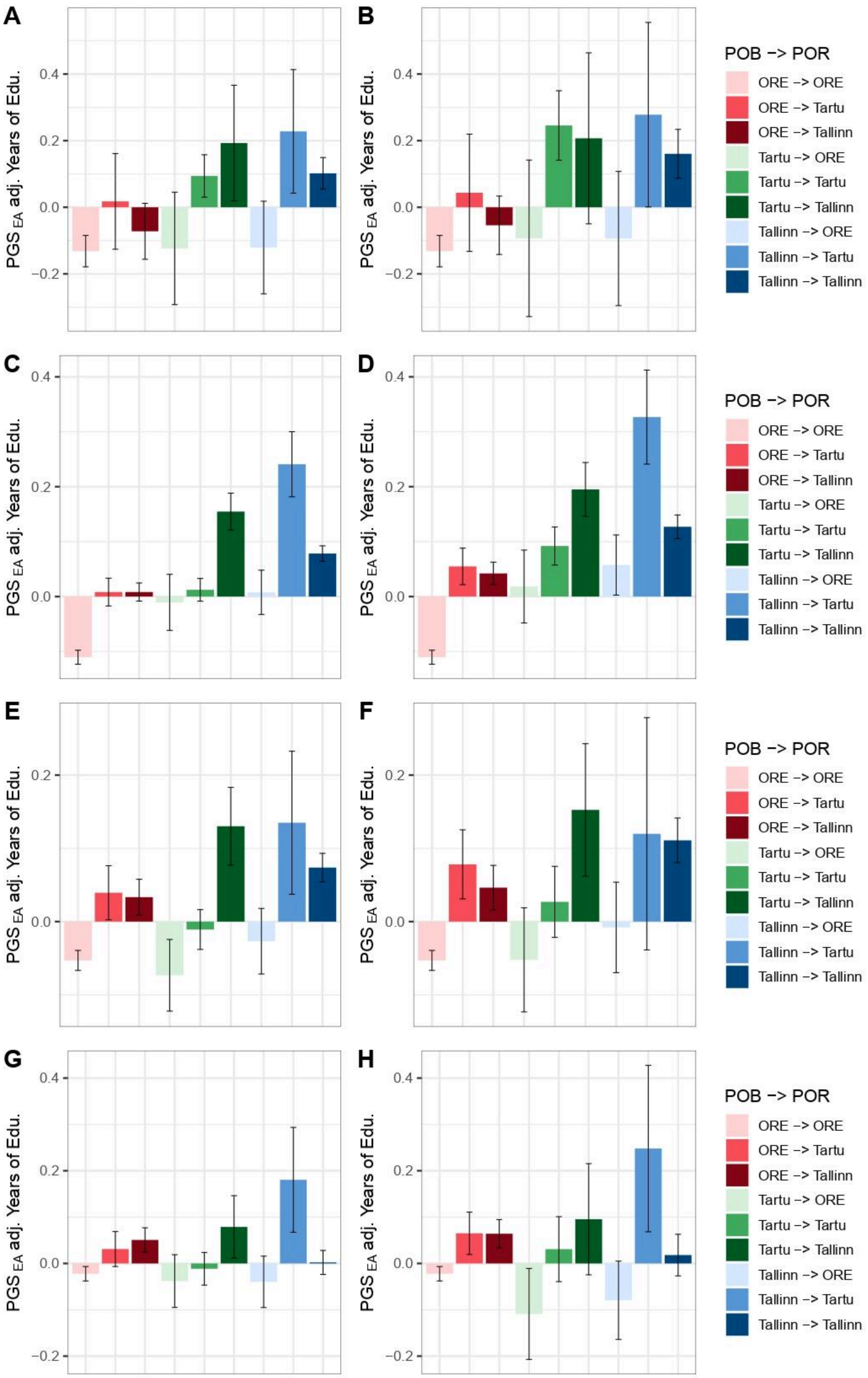
A
B
C
D
E
F
G
H
PGS $_{EA}$ adj. Years of Edu.
0.4
0.2
0.0
−0.2
POB −> POR
ORE −> ORE
ORE −> Tartu
ORE −> Tallinn
Tartu −> ORE
Tartu −> Tartu
Tartu −> Tallinn
Tallinn −> ORE
Tallinn −> Tartu
Tallinn −> Tallinn

**Supplementary Figure 79. $PGS_{EA}$ with EA (years of education) regressed out in migration groups among Estonian participants stratified by age by region of birth (POB) and residence (POR).** Age groups were defined as (A-B) 18-24, (C-D) 25-48, (E-F) 49-64, (G-H) 65+. (A, C, E, G) County-based analysis where POB and POR refer to Tartu County (“Tartu”), Harju County (“Tallinn”) and other counties (“ORE”). (B, D, F, H) City-based analysis, where POB and POR refer to Tartu City (“Tartu”), Tallinn (“Tallinn”) and other counties (“ORE”). $PGS_{EA}$ is adjusted also for demographic and genetic ancestry covariates. Error bars correspond to 95% confidence intervals.

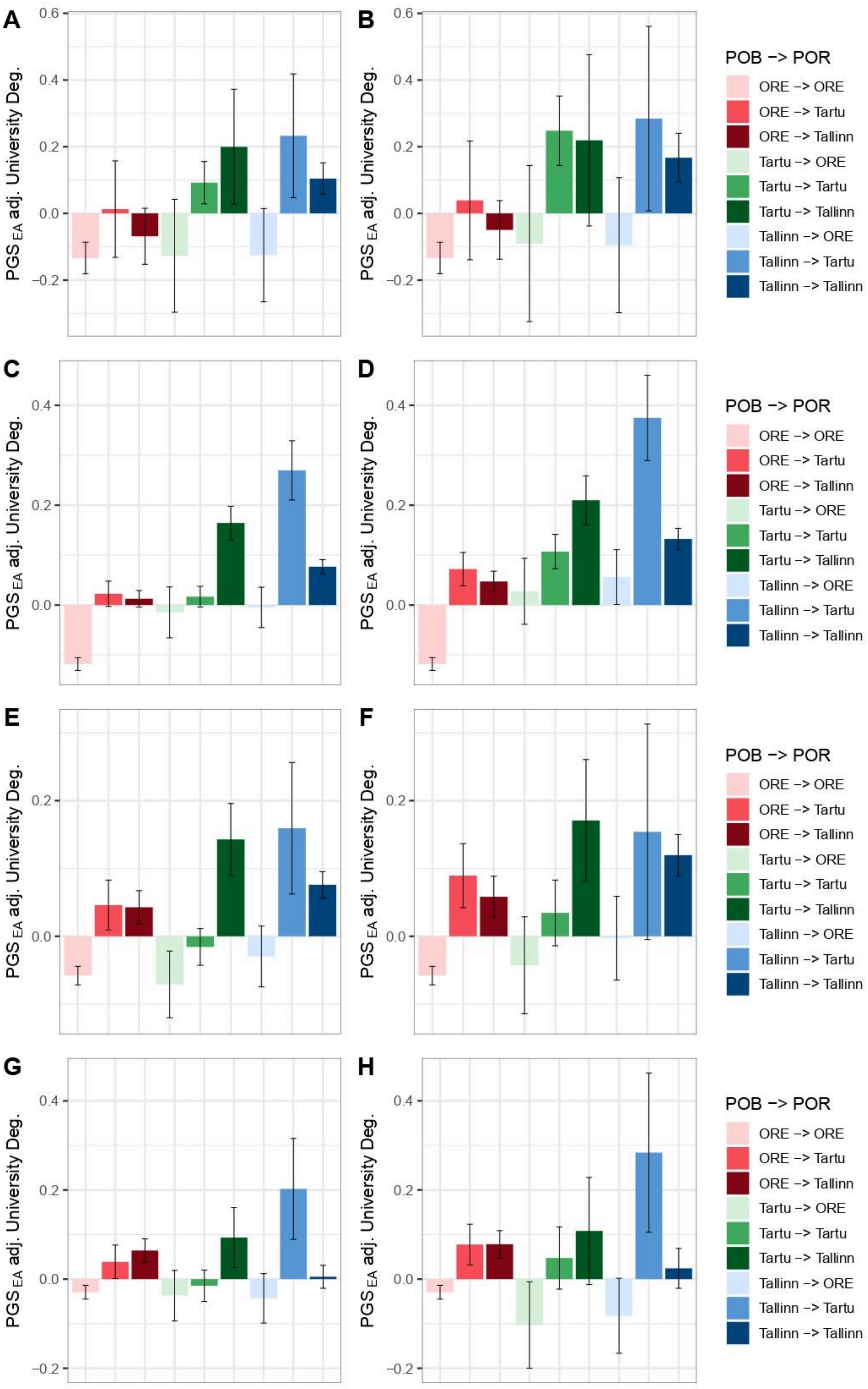
A
B
C
D
E
F
G
H
PGS EA adj. University Deg.
POB -> POR
ORE -> ORE
ORE -> Tartu
ORE -> Tallinn
Tartu -> ORE
Tartu -> Tartu
Tartu -> Tallinn
Tallinn -> ORE
Tallinn -> Tartu
Tallinn -> Tallinn

**Supplementary Figure 80. $PGS_{EA}$ with EA (university degree) regressed out in migration groups among Estonian participants stratified by age by region of birth (POB) and residence (POR).** Age groups were defined as (A-B) 18-24, (C-D) 25-48, (E-F) 49-64, (G-H) 65+. (A, C, E, G) County-based analysis where POB and POR refer to Tartu County ("Tartu"), Harju County ("Tallinn") and other counties ("ORE"). (B, D, F, H) City-based analysis, where POB and POR refer to Tartu City ("Tartu"), Tallinn ("Tallinn") and other counties ("ORE"). $PGS_{EA}$ is adjusted also for demographic and genetic ancestry covariates. Error bars correspond to 95% confidence intervals.

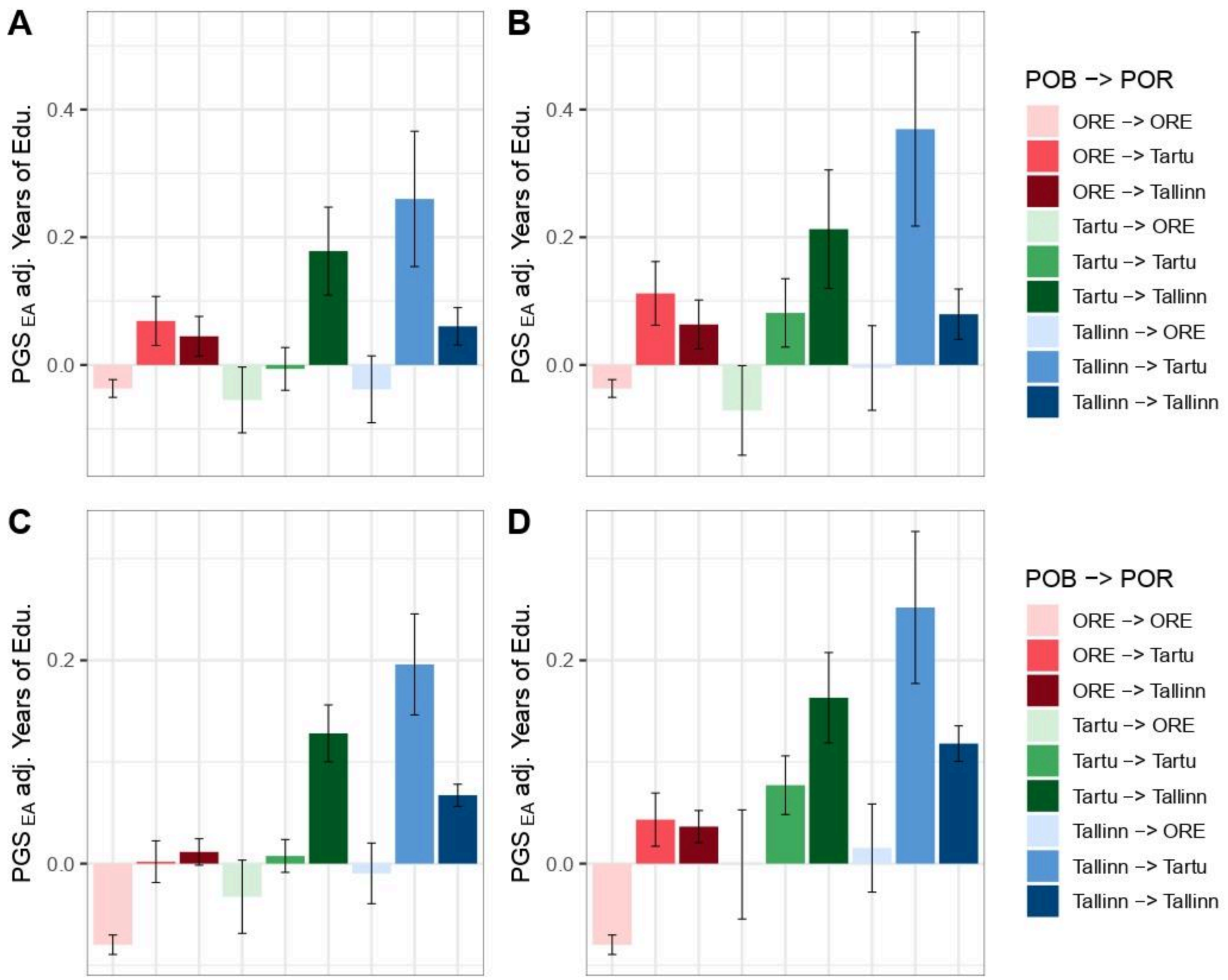

**Supplementary Figure 81. $PGS_{EA}$ with EA (university degree) regressed out in migration groups among Estonian participants stratified by year of joining the biobank by region of birth (POB) and residence (POR).** The periods of joining are (A-B) 2001-2016 and (C-D) 2017-2021. (A, C) County-based analysis where POB and POR refer to Tartu County ("Tartu"), Harju County ("Tallinn") and other counties ("ORE"). (B, D) City-based analysis, where POB and POR refer to Tartu City ("Tartu"), Tallinn ("Tallinn") and other counties ("ORE"). $PGS_{EA}$ is adjusted also for demographic and genetic ancestry covariates. Error bars correspond to 95% confidence intervals.

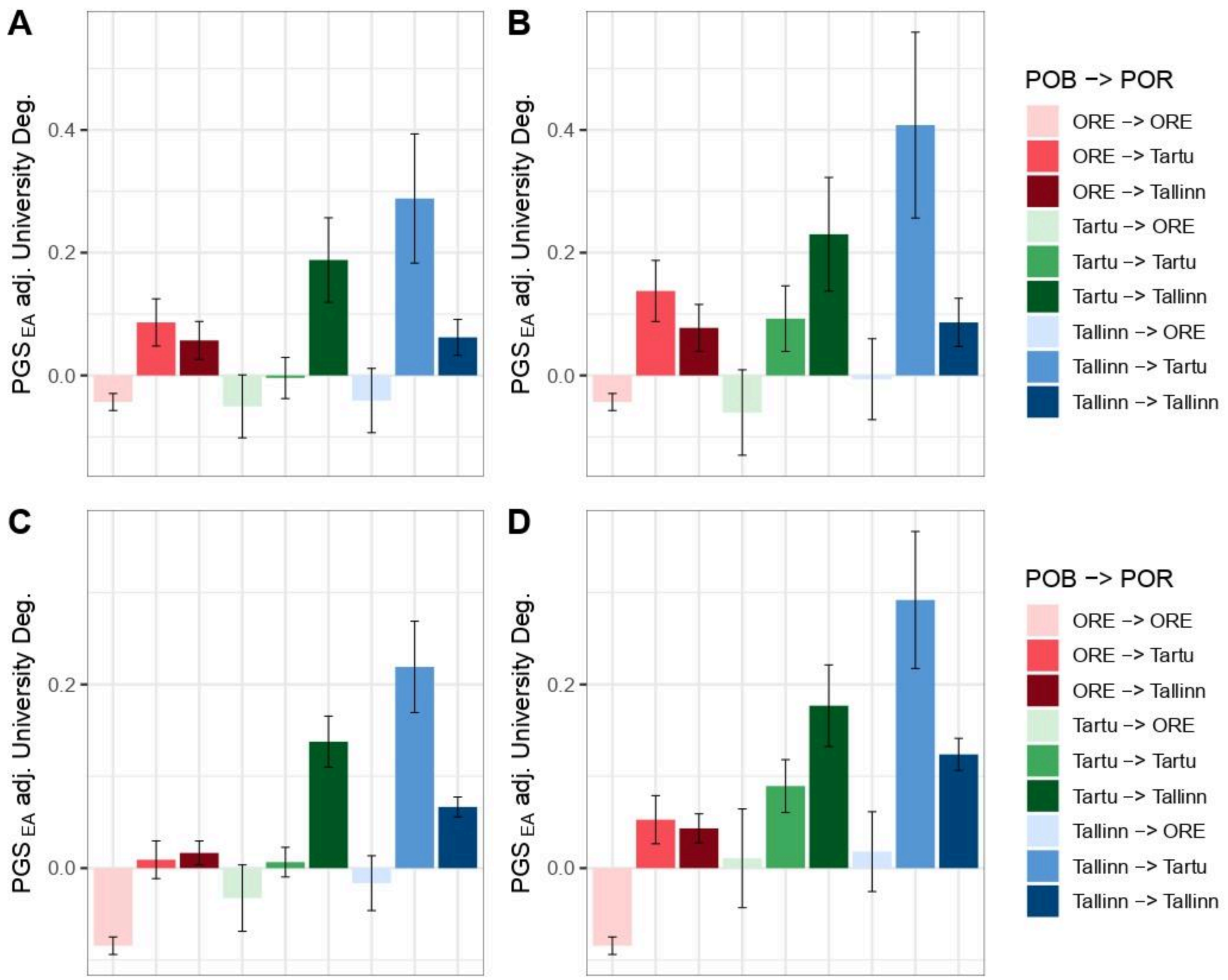

**Supplementary Figure 82. PGS$_{EA}$ with EA (years of education) regressed out in migration groups among Estonian participants stratified by year of joining the biobank by region of birth (POB) and residence (POR).** The periods of joining are (A-B) 2001-2016 and (C-D) 2017-2021. (A, C) County-based analysis where POB and POR refer to Tartu County ("Tartu"), Harju County ("Tallinn") and other counties ("ORE"). (B, D) City-based analysis, where POB and POR refer to Tartu City ("Tartu"), Tallinn ("Tallinn") and other counties ("ORE"). PGS$_{EA}$ is adjusted also for demographic and genetic ancestry covariates. Error bars correspond to 95% confidence intervals.
